# Proteome and functional decline as platelets age in the circulation

**DOI:** 10.1101/2021.06.02.446106

**Authors:** Harriet E. Allan, Melissa A. Hayman, Simone Marcone, Melissa V. Chan, Matthew L. Edin, Tania Maffucci, Abhishek Joshi, Laura Menke, Marilena Crescente, Manuel Mayr, Darryl C. Zeldin, Paul C. Armstrong, Timothy D. Warner

## Abstract

Anucleate platelets circulate in the blood of healthy individuals for approximately 7-10 days during which time their protein composition may change. We hypothesized such changes would be linked to altered structure and function. Here, we separated platelets of different ages based on mRNA content and characterised them using proteomics, immunofluorescence and functional assays. Total protein content was 45±5% (n=4) lower in old platelets compared to young platelets. Predictive proteomic pathway analysis identified associations with 28 biological processes, notably increased haemostasis in young platelets and apoptosis in old platelets. Further studies confirmed platelet ageing was linked to a reduction decrease in cytoskeletal proteins, a reduction in mitochondria number, and lower calcium dynamics and granule secretion. This work delineates physical and functional changes in platelets as they age and serves as a base to examine differences associated with altered mean age of platelet populations in conditions such as immune thrombocytopenia and diabetes.

## Introduction

Blood platelets are cellular fragments produced from megakaryocytes which circulate for approximately 10 days in healthy individuals.^1–3^ These anucleate fragments contain a complex array of extracellular proteins, signalling pathways and intracellular machinery including storage granules, canalicular systems, mitochondria, and contractile proteins. Platelets are dynamic and metabolically active, responding very rapidly as central players in haemostasis. Analyses of the platelet proteome have revealed a high degree of similarity in protein content among healthy individuals, with over 3000 proteins detected.^4^ Being cellular fragments, platelets lack a nucleus and have only limited capacity to synthesize new proteins. Proteomic analyses to date have largely been conducted on platelets separated as a single cell type from normal whole blood, and so represent the collective proteome across platelets of all ages.

Despite lacking a nucleus, newly formed platelets contain an array of the messenger ribonucleic acids (mRNA) which were present in their progenitor megakaryocyte.^5–7^ These residual mRNAs can be used as an indicator of platelet age as they are lost from platelets as they circulate and platelets have limited capacity to generate new mRNA.^8–12^ Numerous studies have reported that newly formed platelets, i.e. those with the highest levels of mRNA, also known as reticulated platelets or the ‘immature platelet fraction’, are hyper-reactive with an increased thrombotic potential.^13–18^ This hyper-reactivity has been linked to a number of pathological states, including diabetes mellitus and chronic kidney disease, in which there is both increased platelet turnover and higher incidence of acute coronary syndromes associated with reduced effectiveness of standard anti-platelet therapies.^19–23^

A hypothetical explanation for the age-related difference in reactivity is that younger platelets have a greater array of functional pathways to bring to bear to haemostatic processes; they have the full complement of proteins derived from the progenitor megakaryocytes, while these have become degraded or lost in older platelets without the general possibility for replacement. To test this idea, we developed and validated protocols to separate platelets according to circulatory age and carried out proteomic, immunofluorescence and functional assays to delineate physical and functional changes in platelets as they normally age within the circulation.

## Results

### Platelets can be separated by circulatory age using cell sorting based upon thiazole orange fluorescence

To validate assays to use in samples from healthy human individuals, we began with experiments in mice in which we could employ *in vivo* labelling. Temporal *in-vivo* antibody labelling in mice and cell sorting followed by quantitative real time polymerase chain reaction (qRT-PCR) demonstrated that the newest circulating platelets, i.e. young platelets (<24 hours old), had significantly lower mean cycle threshold (Ct) values for *ITGA2B*, *PF4* and *TUBB1* than old platelets (2-5 days). This is consistent with the loss of megakaryocytic mRNAs as platelets age and demonstrates that the levels of these megakaryocytic mRNAs are indicators of platelet circulatory age (Figure 1A, p<0.05, n=4). Next, as we cannot conduct *in vivo* labelling in human subjects, we assessed to what extent thiazole orange (TO) staining correlated with temporal antibody staining in mice. Analysis of these experiments demonstrated significantly higher TO fluorescence in young platelets compared to older platelets (Figure 1B, n=4). Following this we similarly sorted human platelets on the basis of TO fluorescence intensity and defined three platelet populations: young platelets (the top 10% of TO fluorescence), old platelets (the bottom 30%; Figure 1C), and for completeness intermediate-aged platelets (defined as the 50% between young and old platelets). To further validate this approach for the separation of human platelets on the basis of circulatory age, we conducted qRT-PCR as we had for mouse platelets and demonstrated the same pattern of mRNAs; i.e. that young platelets had significantly lower mean cycle threshold (Ct) values for *ITGA2B*, *PF4*, *TUBB1* than both old platelets (Figure 1D; p<0.05, n=3) and intermediate-aged platelets (*ITGA2B*, 27.0±0.3 vs. 31.9±1.6; *PF4*, 23.9±0.4 vs. 27.9±1.7; *TUBB1*, 25.8±1.4 vs. 30.4±2.8; young vs. intermediate platelets, p<0.05 for all, n=3). There were highly significant correlations between TO-determined platelet age and log2 fold differences in mRNA, as determined from Ct values, for ITGA2B (r^2^ 0.53, p<0.03), PF4 (r^2^ 0.55, p<0.02) and TUBB1 (r^2^ 0.65, p<0.008) (Figure 1E). The levels of a further ten mRNAs relevant to platelet function were also noted to decline strongly, although for these Ct values >40 in old platelets precluded full analyses (Supplemental Table 1). From these data we can conclude that our TO staining and sorting protocols allow the separation of human platelets from healthy volunteers on the basis of circulatory age as confirmed by decline in megakaryocytic mRNAs. Importantly, activation status (determined by P-selectin expression and PAC-1 and Annexin V binding) did not change during the staining and sorting protocol (Supplemental Figure 1) indicating that platelets were not adversely activated by these processes.

**Figure 1:**
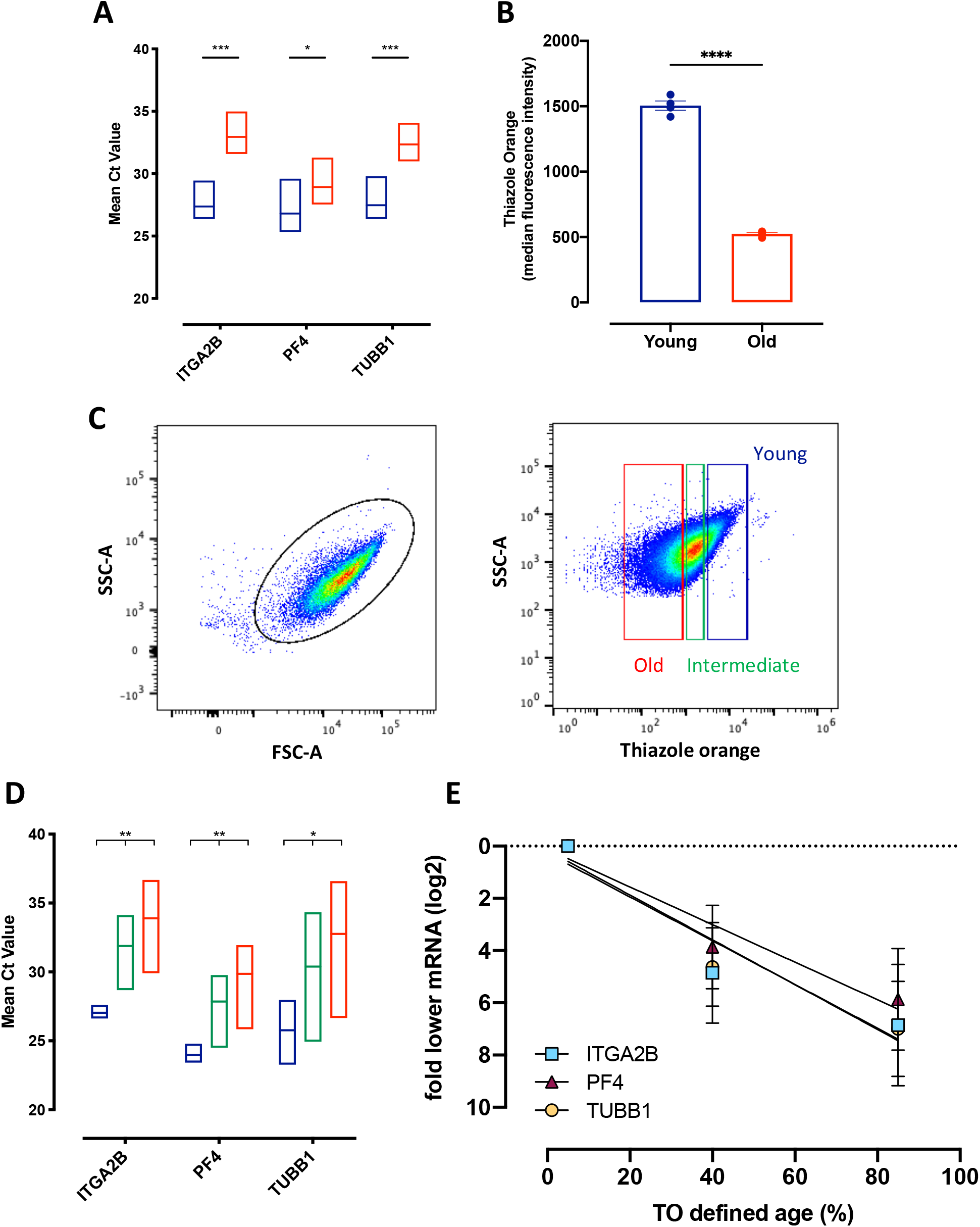
RNA expression in temporally labelled murine platelets and thiazole orange sorted human platelets. **A.** qRT-PCR of platelet-specific mRNAs; *ITGA2B, PF4, TUB*B1 showing mean Ct value for young (blue) and old (red) murine platelets sorted on the basis of *in vivo* antibody labelling. **B.** Thiazole orange staining in young (blue), and old (red) temporally labelled murine platelets**. C.** Gating strategy for two human platelet subpopulations based on thiazole orange/SSC-A; young (blue), top 10%; intermediate (green) middle 50% and old (red) bottom 30%. **D.** qRT-PCR of platelet-specific mRNAs; *ITGA2B, PF4, TUBB1* showing mean Ct value for young (blue), intermediate (green) and old (red) platelets sorted on the basis of thiazole orange staining. **E**. mRNA levels across TO-defined age in human sorted platelets. Data presented as min-max, with line at mean, or mean ±SEM (*p<0.05, ** p<0.01, *** p<0.005, ****p<0.001; n=3-4).

### Platelet ageing is associated with a decline in total protein content

We noted significant correlation (r^2^ 0.49, p<0.01, n=12) between TO-defined platelet age and protein content per platelet (young, 8.7±2.6pg; intermediate-aged, 6.6±0.7pg; old, 4.7±1.2pg; Figure 2A; p<0.05, n=4). In addition, there were strong correlations between total protein content per platelet and log2 fold differences in mRNA, determined from Ct values, for ITGA2B (r^2^ 0.55, p<0.02), PF4 (r^2^ 0.57, p<0.02) and TUBB1 (r^2^ 0.68, p<0.006 (Figure 2B). Notably these correlations are incompatible with the notion that mRNA and protein vary by platelet size rather than age, as protein varies across the TO-defined populations linearly and mRNA by powers of 2. As further confirmation, immunofluorescence revealed there was no difference in the cross-sectional area of our sorted young and old platelets (young platelets, 8.9±0.2μm^2^; old platelets, 8.5±0.5μm^2^; Figure 2C, p>0.05, n=4).

**Figure 2:**
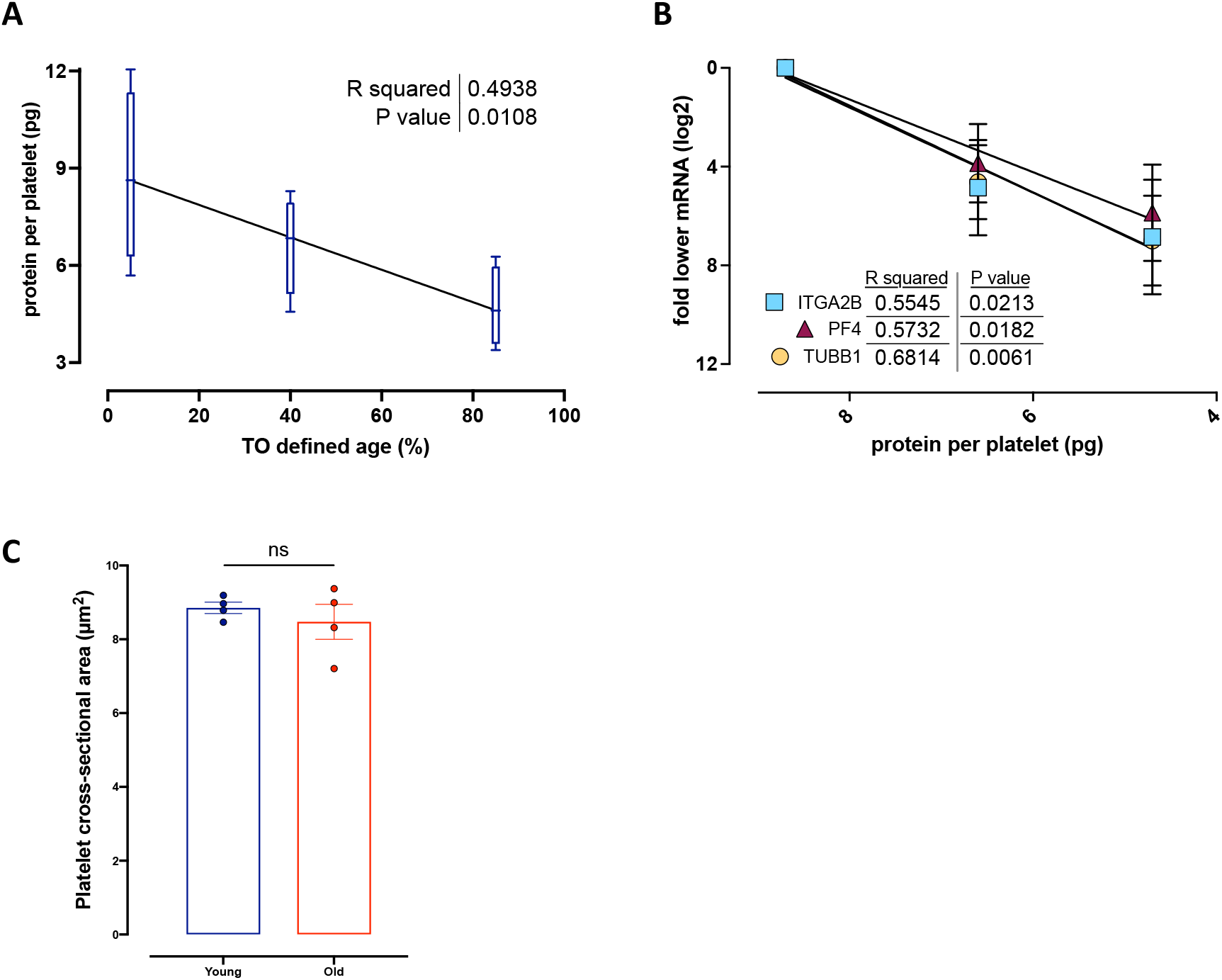
Proteomic and transcriptomic correlations in equally sized sorted platelets. **A.** Quantification of protein content per platelet correlated with mean TO-defined ‘age’. **B.** Correlation between protein content and log-2 fold decreases of individual mRNAs ITGA2B, PF4 and TUBB1. **C.** Quantification of platelet cross-sectional area in sorted young (blue) and old (red) platelets.

### Proteomic analysis identified differences in proteins affecting fundamental biological processes

Proteomic analysis identified 583 proteins within the sorted platelets (Supplemental Table 2) of which 94 proteins were significantly modulated among the three subpopulations (p<0.05; Supplemental Table 3). Targeted analysis between young and old platelets identified relative differences in the levels of 78 proteins (Table 1, Figure 3A).

**Table 1:**
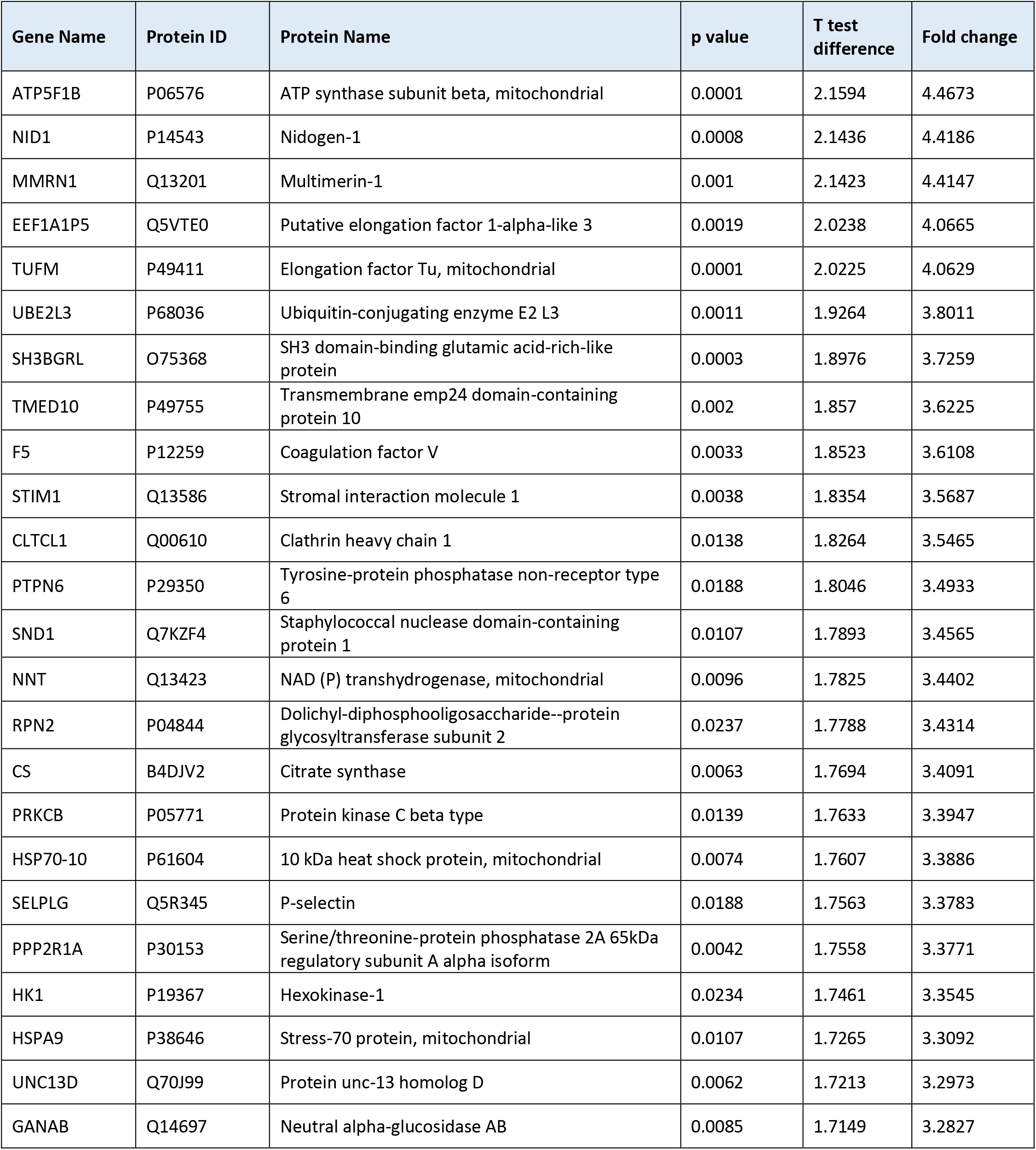
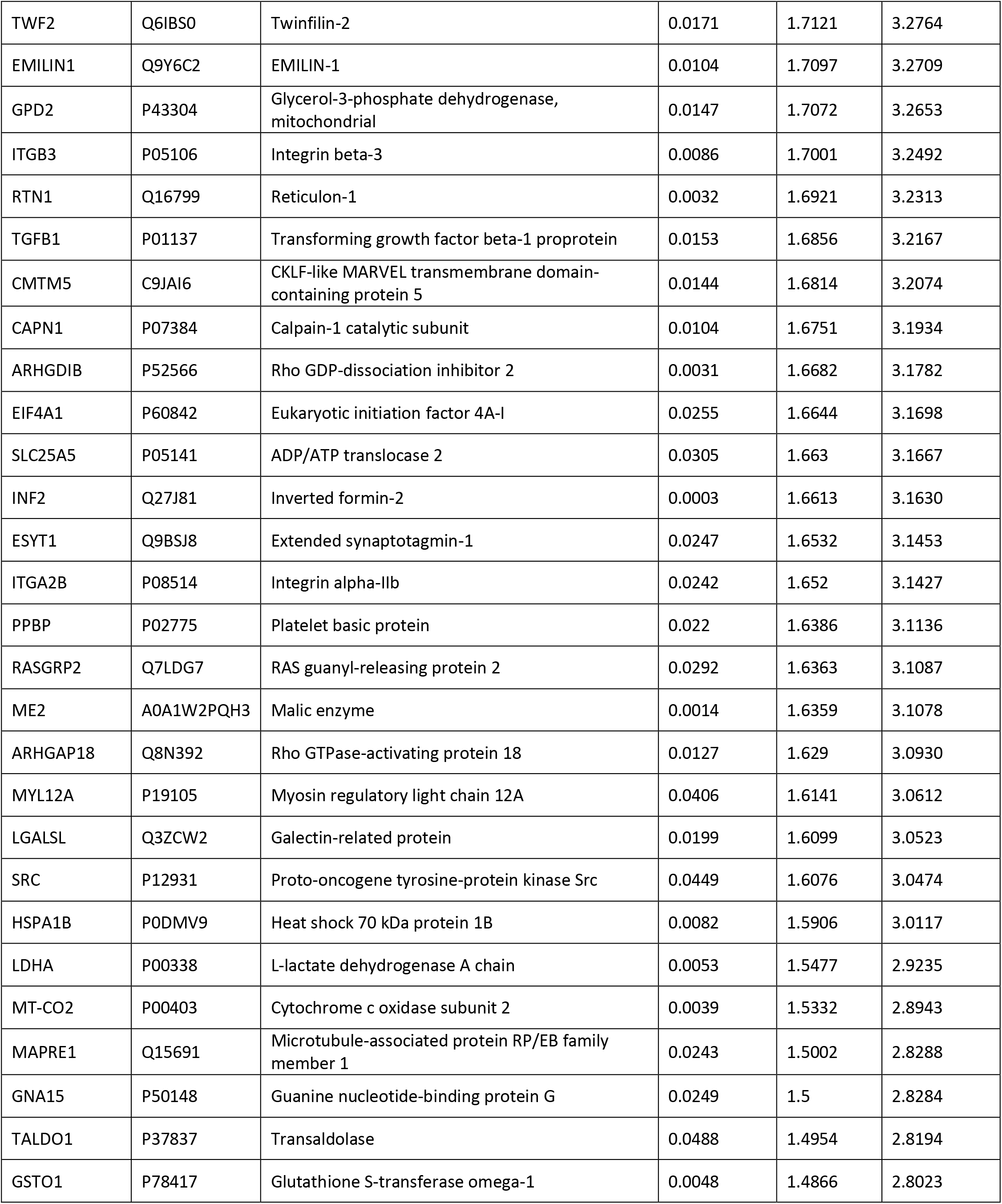
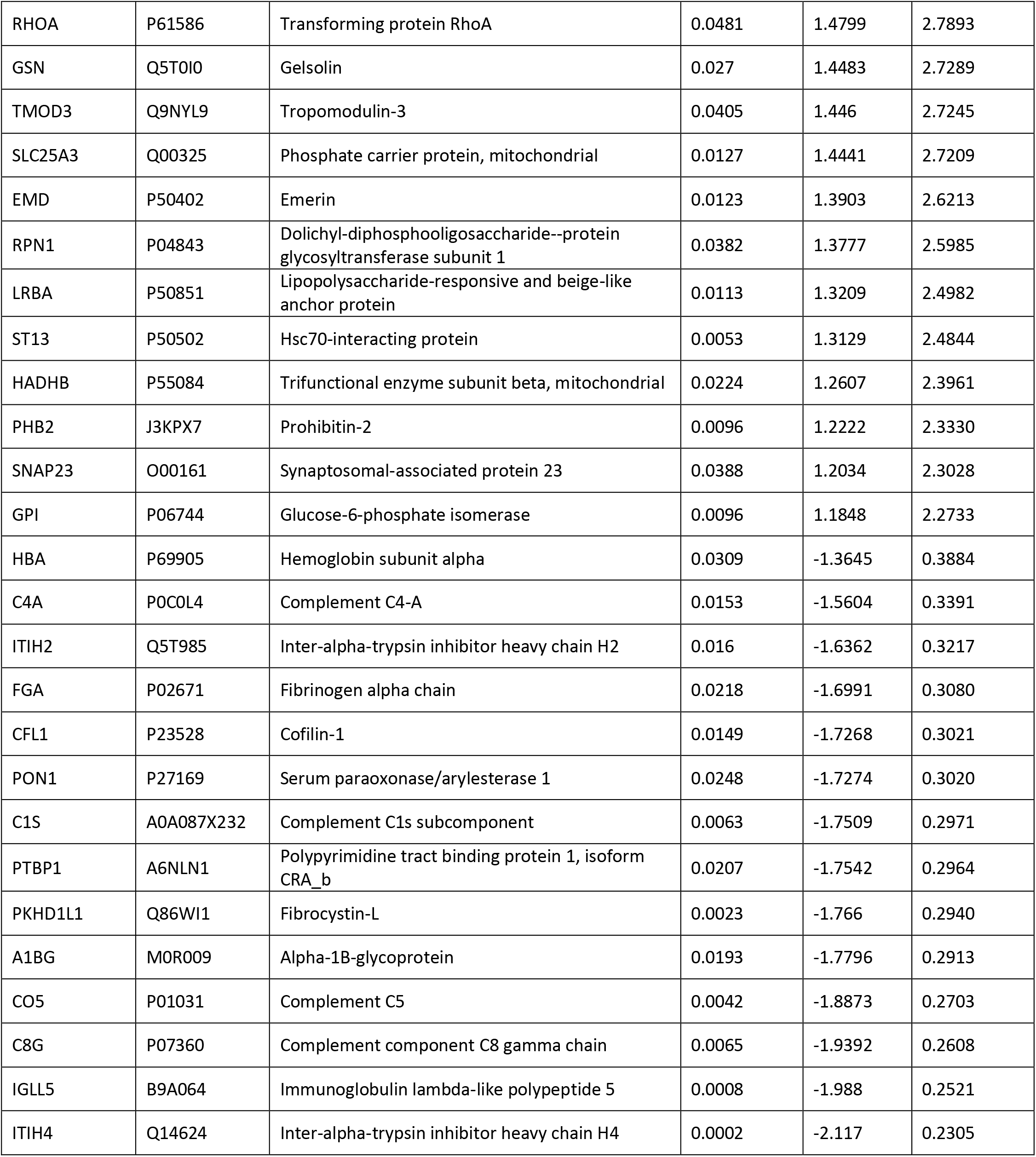
Proteins with significantly altered expression in young and old platelets.

**Figure 3:**
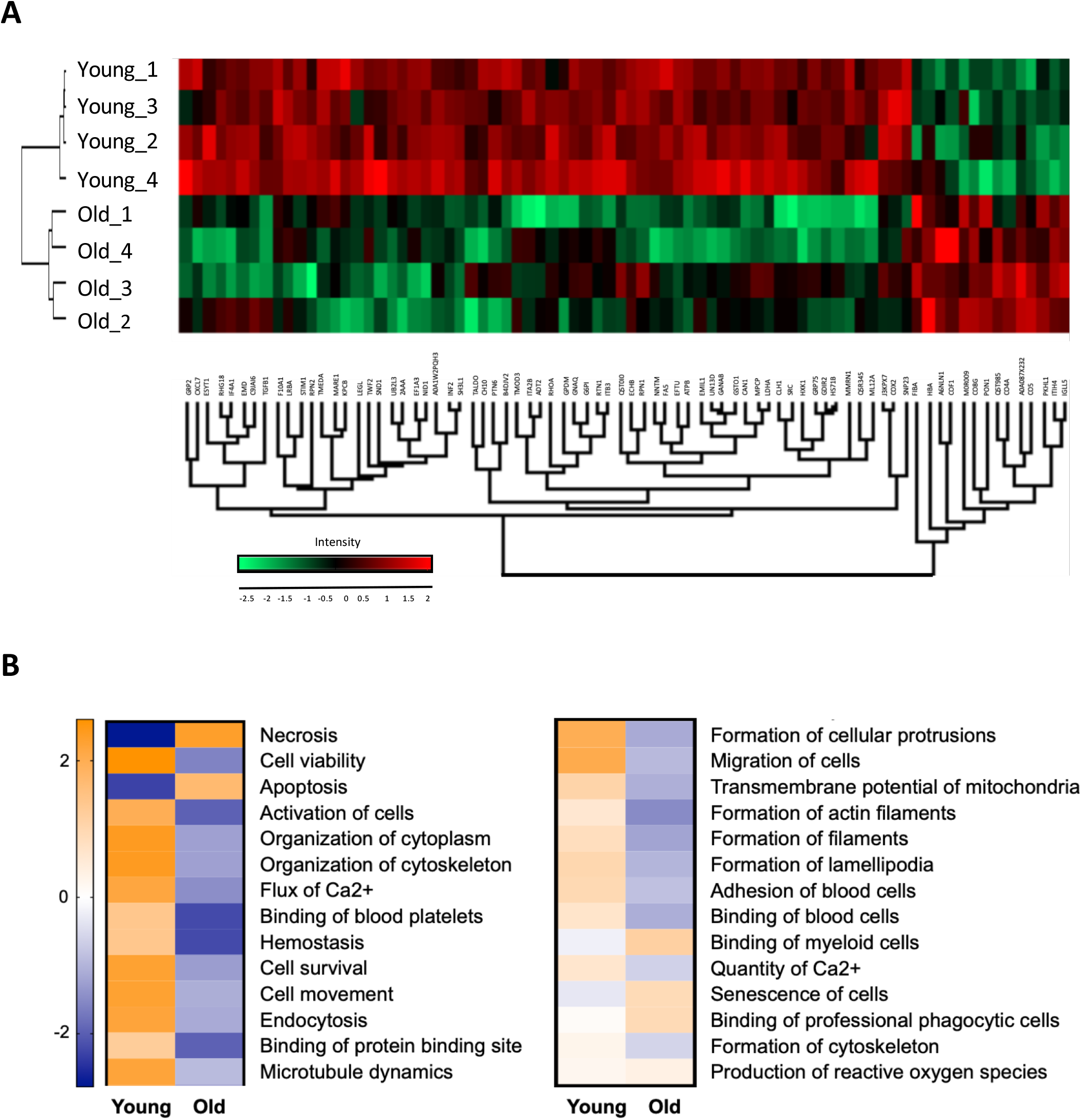
Proteomic and Ingenuity Pathway Analysis of young and old platelets. **A.** Hierarchical clustering heatmap showing the 78 proteins that were significantly modulated between young and old platelets; for each protein the log_2_ (intensity) was measured in samples from 4 individuals (noted 1 - 4) with low content indicated in green and high content indicated in red. **B.** Ingenuity pathway analysis showing biological functions predicted to be affected by the differential protein content of young platelets relative to old platelets. The biological processes are listed according to their Z-score: orange indicates a predicted upregulation, blue a predicted downregulation. Data presented as mean z-score (n=4).

Ingenuity pathway analysis (IPA) predicted an association between the 78 altered proteins and 28 biological processes and functions (Figure 3B). Twenty-two of these processes were predicted as higher in young platelets including haemostasis, binding of platelets and calcium flux, as well as transmembrane potential of mitochondria. The remaining six functions were predicted as higher in old platelets including apoptosis and senescence (Figure 3B).

### Old platelets have reduced mitochondrial number and activity

Proteomic analysis demonstrated a relative reduction in the levels of key mitochondrial proteins in old platelets, notably citrate synthase and ADP/ATP translocase 2 (Table 1). Further investigation using immunofluorescence and confocal microscopy for the mitochondrial protein, TOM20, established that the reduction in mitochondrial proteins was accompanied by a decrease in the number of mitochondria per platelet from 11±1 in young platelets to 5±1 in old platelets (Figure 4A-C; p<0.05, n=4). This observed reduction was not due to mitochondrial fusion as there was no change in mitochondrial cross-sectional area (young platelets, 177±10nm^2^; old platelets, 156±20nm^2^; Figure 4D; p<0.05, n=4). In agreement with the IPA, we identified a reduction in mitochondrial membrane potential in old platelets compared to young platelets (Figure 4E, F; p<0.05, n=4). Furthermore, phosphatidylserine exposure, commonly associated with a reduction in mitochondrial membrane potential, was higher basally in old platelets as indicated by higher basal levels of annexin V binding (20±4% vs 3±1% positive in old vs. young platelets; Figure 4G-I; p<0.05, n=6).

**Figure 4:**
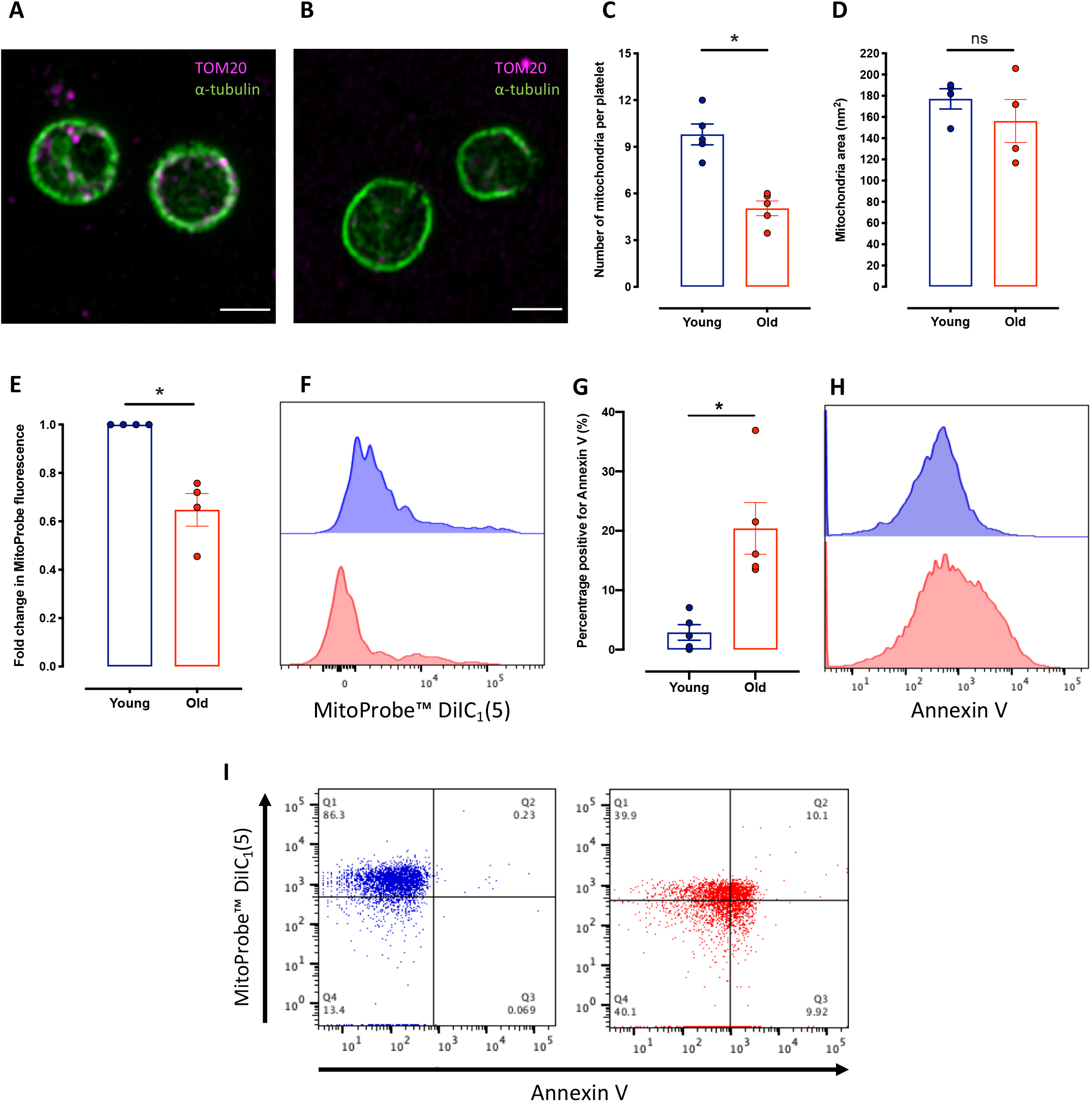
Analysis of mitochondria characteristics in young and old platelets. **A and B.** Representative confocal microscopy images (Zeiss LSM880 with Airyscan; 63x objective, 1.4 Oil DICII) showing mitochondria; TOM20 (magenta) and α-tubulin (green) in (**A**) young platelets and (**B**) old platelets, scale bar represents 2μm.. **C.** Quantification of the average number of mitochondria per platelet. **D.** Quantification of cross-sectional mitochondrial area in young and old platelets. **E.** Quantification of mitochondrial membrane potential in young and old platelets. **F.** Representative histogram of MitoProbe™ DiIC_1_(5) fluorescence in young (blue) and old (red) platelets. **G.** Quantification of phosphatidylserine exposure; annexin V binding. **H.** Representative histogram of annexin V binding in young (blue) and old (red) platelets. **I.** Representative dot plots showing MitoProbe DiIC_1_(5) fluorescence vs. Annexin V binding in young (blue) and old (red) platelets. Data presented as mean±SEM (*p<0.05, ***p<0.005, n=4).

### Old platelets have an altered cytoskeleton resulting in reduced platelet spreading

Proteomics also indicated a reduction in the amounts of a number of cytoskeletal binding proteins including twinfilin-2, emerin and gelsolin, suggesting age related alterations to cytoskeletal structure (Table 1). Immunofluorescence demonstrated a significant decrease in the fluorescence intensity for α-tubulin (2435±354 arbitrary units (AU) vs. 1034±79AU; Figure 5A-C; p<0.05, n=4) and F-actin (2166±110AU vs. 1364±139AU; Figure 5D-F; p<0.05, n=5) in young vs. old platelets, respectively. Furthermore, western blotting confirmed the reduction in the expression of α-tubulin and β-actin (Supplemental Figure 2A-B). Notably, the reduction in cytoskeletal proteins was not accompanied by a change in the cross-sectional area of resting platelets (Figure 2C).

**Figure 5:**
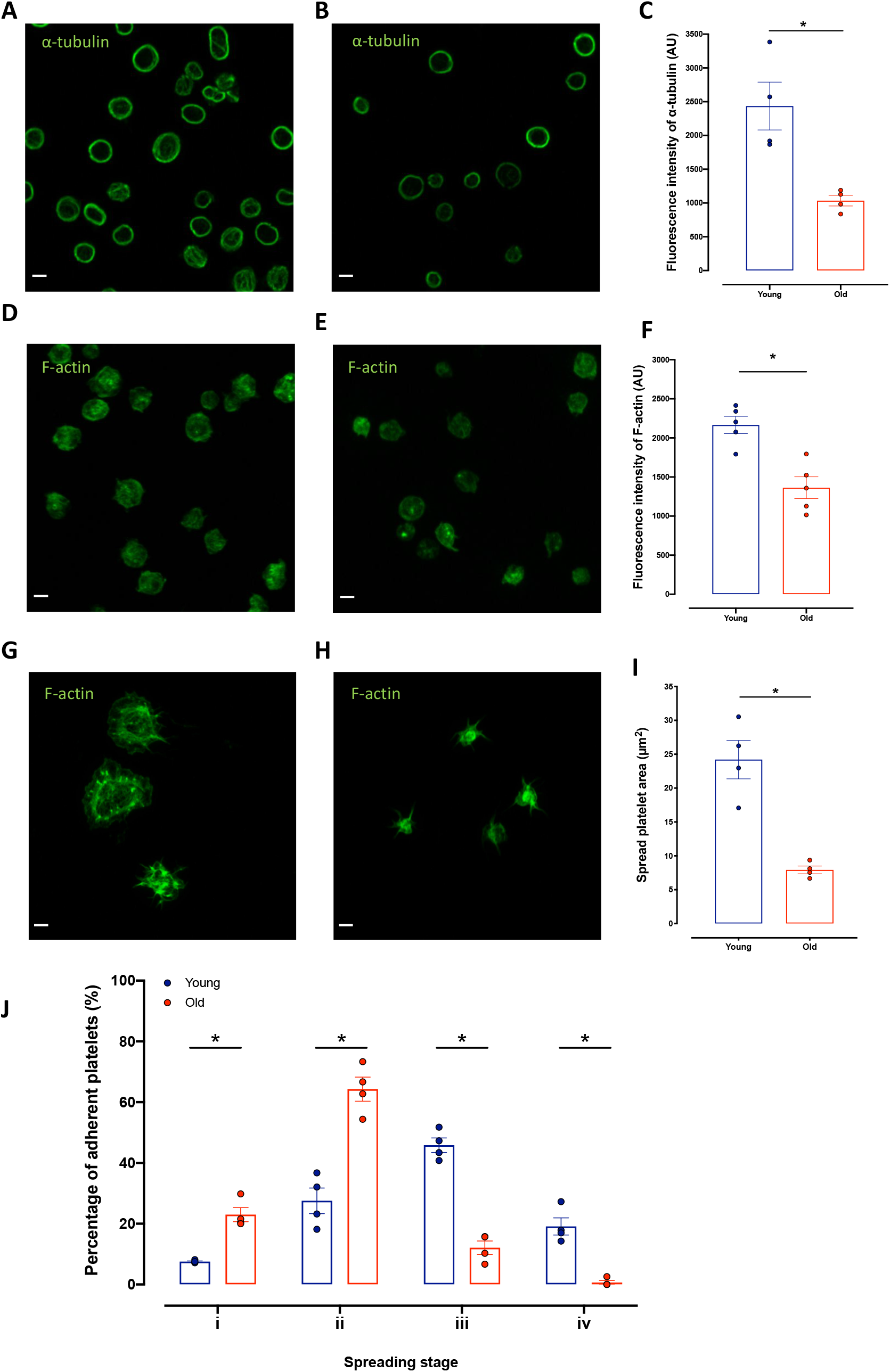
Analysis of platelet cytoskeletal structure. **A and B.** Representative confocal microscopy images visualizing α-tubulin in (**A**) young and (**B**) old platelets. **C.** Quantification of α-tubulin fluorescence intensity. **D and E.** Representative confocal microscopy images visualizing F-actin in (**D**) young and (**E**) old platelets. **F.** Quantification of F-actin fluorescence intensity. **G and H.** Representative confocal microscopy images of platelets spread on fibrinogen stained with F-actin in (**G**) young and (**H**) old platelets. **i.** Quantification of spread platelet area. **J.** Quantification of stage of platelet spreading: i, adhered but not spread; ii, filopodia; iii, lamellipodia; and iv, fully spread in young (blue) and old (red) platelets. Images acquired on a Zeiss LSM880 with Airyscan confocal microscope; 63x objective, 1.4 Oil DICII, scale bar represents 2μm; data presented as mean±SEM (*p<0.05, n=4).

Following spreading on fibrinogen, old platelets showed a marked reduction in their spread area (24.2±2.8μm vs. 7.9±0.6μm; young vs. old platelets respectively; p<0.05, n=4; Figure 5G-I). Categorization of spreading stage demonstrated that old platelets were able to form filopodia but unlike young platelets did not transition into lamellipodia and fully spread (Figure 5J). Furthermore, the reduction in spreading was accompanied by a decrease in the percentage of platelets forming actin nodules (63±1% vs. 24±1%; young vs. old platelets; p<0.05, n=4; Supplemental Figure 2C-E).

### Platelet ageing is associated with alterations in intracellular protein components

Proteomics also identified a relative increase in the levels of a small number of circulating proteins as platelets age (complement C5, C1s and C4a, and fibrinogen alpha chain; Table 1). Microscopy confirmed an increased abundance of complement C4 and fibrinogen in old platelets (respectively, 661±150AU vs. 1519±181AU and 580±93AU vs. 961±179AU, in young vs. old platelets; Figure 6C, F; p<0.05, n=4) localized both at the cell periphery and within the cytoplasm consistent with internalization (Figure 6A-F).

**Figure 6:**
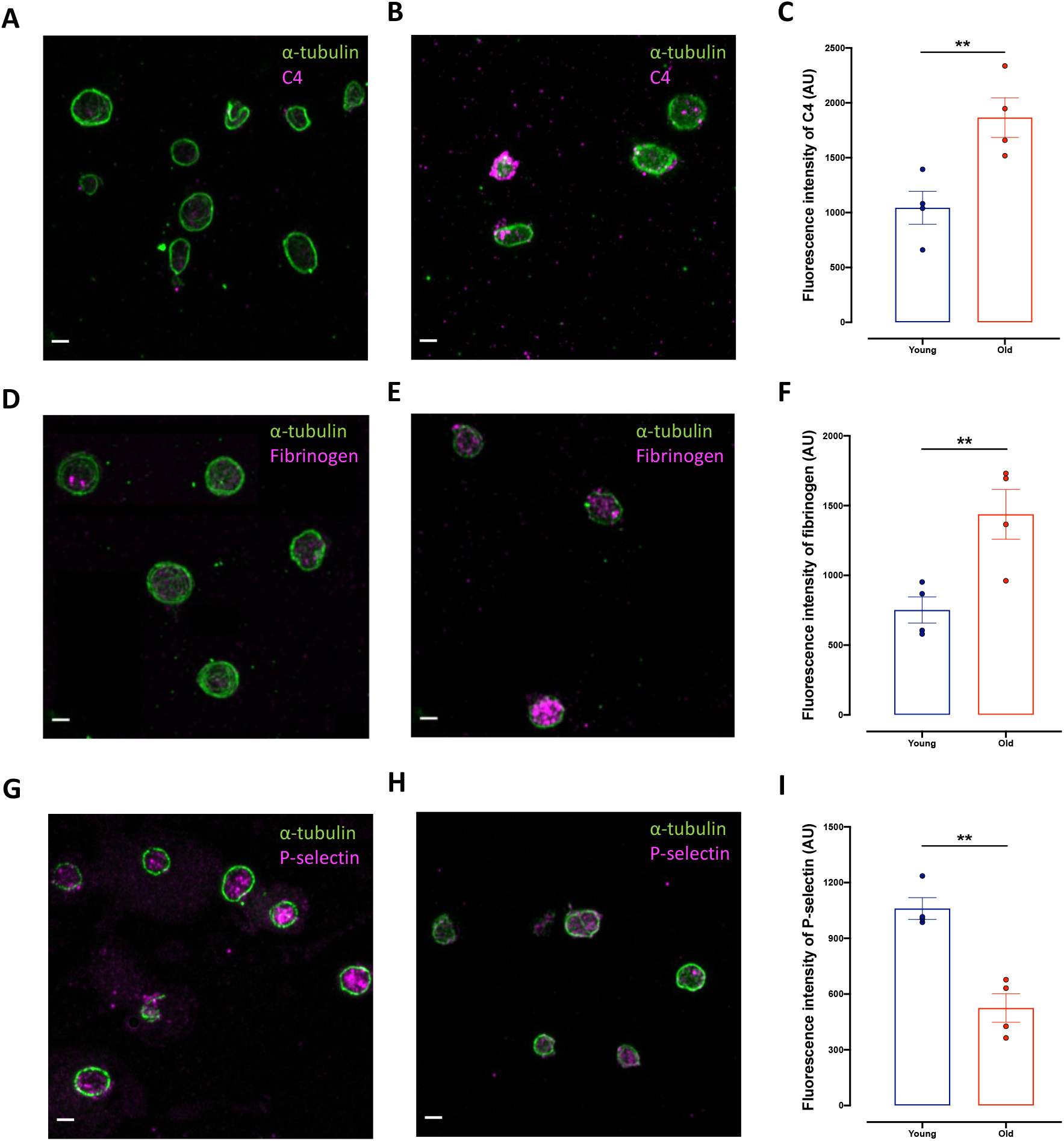
Analysis of intracellular components in young and old platelets. **A and B.** Representative confocal microscopy images visualizing α-tubulin (green) and complement protein C4 (magenta) in (**A**) young and (**B**) old platelets. **C.** Quantification of complement C4 fluorescence intensity. **D and E.** Representative confocal microscopy images visualizing α-tubulin (green) and fibrinogen (magenta) in (**D**) young and (**E**) old platelets. **F.** Quantification of fibrinogen fluorescence intensity. **G and H.** Representative confocal microscopy images visualizing α-tubulin (green) and P-selectin (magenta) in (**G**) young and (**H**) old platelets. **I.** Quantification of P-selectin fluorescence intensity.

### Platelet ageing is associated with a reduction in activation and secretory pathways

To fairly assess the relative activations of young and old platelets, imaging flow cytometry of fabricated mixed populations of 50% young and 50% old platelets demonstrated that young platelets responded more rapidly to stimulation, forming the core of aggregates with old platelets binding to the periphery (Figure 7A). Young platelets were present in 95±2% of aggregates compared with 42±6% of old platelets (Figure 7B). Notably, 57±6% of aggregates were composed exclusively of young platelets but only 4±2% were composed exclusively of old platelets (Figure 7C; p<0.05, n=3). These different aggregatory responses were associated with further distinct functional differences. Young platelets demonstrated a stronger increase in intracellular calcium following exposure to thrombin receptor activating peptide (TRAP-6) than old platelets (AUC; 7712±1684AU vs. 2532±1221AU). Similarly, ionomycin elicited a significantly stronger calcium response in young platelets than old platelets (AUC; 69830±12864AU vs. 27973±10195AU) suggesting young platelets have larger calcium stores (Figure 7D-G; p<0.05, n=4). Consistent with a reduction in activation pathways associated with ageing, old platelets had reduced granule secretion indicated by a significant reduction in P-selectin expression and ATP release following TRAP-6 stimulation (respectively, 0.39±0.05 and 0.36±0.04-fold changes compared to young platelets; Figure 7H-I; p<0.05, n=5). This reduction in P-selectin exposure may be due to a decline in P-selectin content identified in the proteomics (Table 1) and immunofluorescence (1061±59 vs. 524±76 in young vs. old platelets; Figure 6G-I; p<0.05, n=4;). Furthermore, synthesis and release of eicosanoids in response to collagen stimulation was altered between the two platelet subpopulations; old platelets released significantly lower quantities of thromboxane B_2_ (TXB_2_), prostaglandin E_2_ (PGE_2_) and 12-hydroxyeicosatetraenoic acid (12-HETE) than young platelets (Figure 7J-L; p<0.05, n=8).

**Figure 7:**
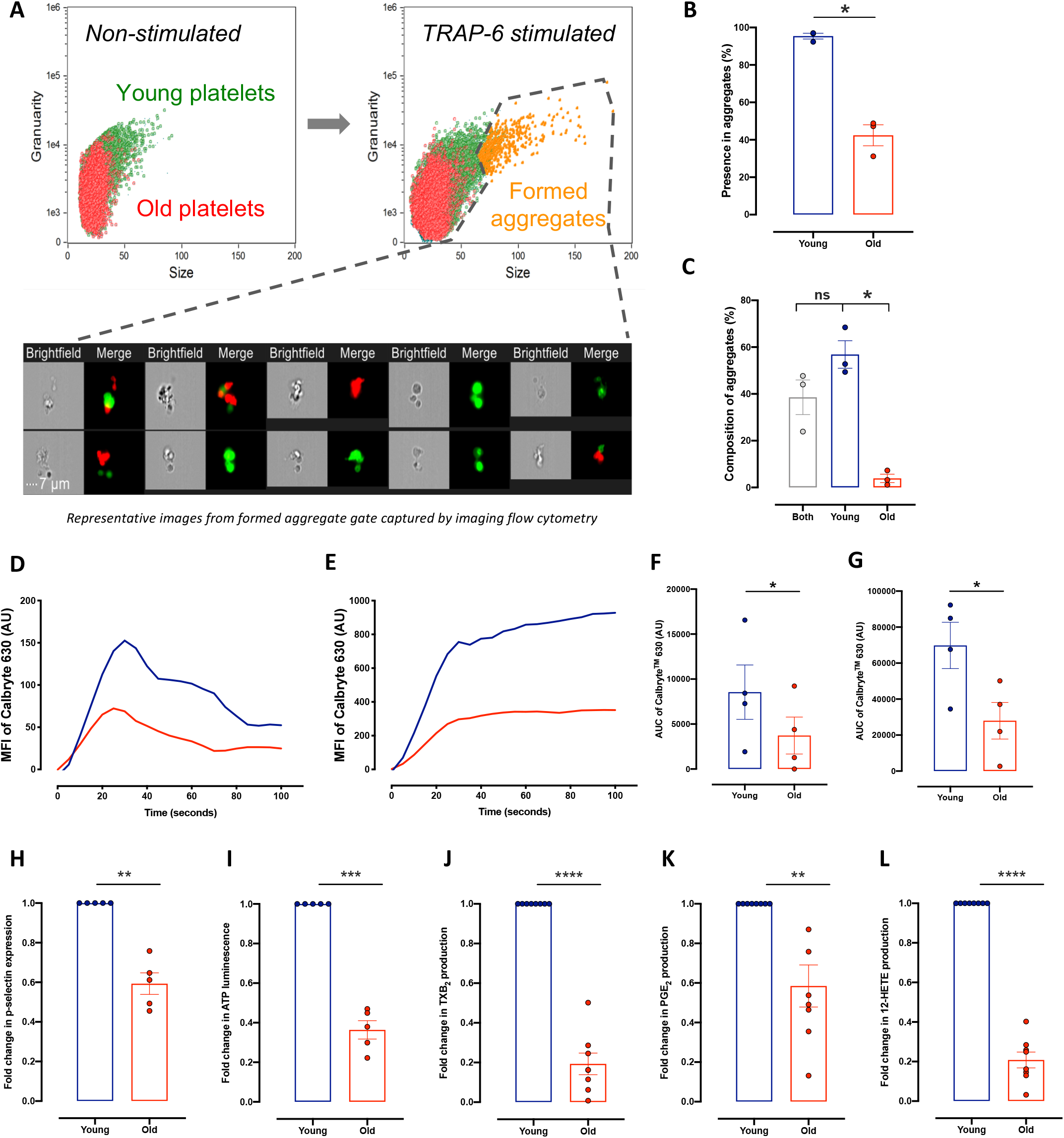
Characterization of function in young and old platelets. **A.** Representative imaging flow cytometry dot plots and images showing young platelets (green) and old platelets (red) in non-stimulated and TRAP-6 stimulated samples. **B.** Quantification of the percentage of young and old platelets within the formed aggregates. **C.** Quantification of the composition of aggregates. **D.** Representative calcium dynamic traces in young (blue) and old (red) platelets following TRAP-6 stimulation. **E.** Representative calcium dynamic traces in young (blue) and old (red) platelets following ionomycin stimulation **F.** Quantification of area under the curve of the calcium trace following TRAP-6 stimulation. **G.** Quantification of area under the curve of the calcium trace following ionomycin stimulation. **H.** Quantification of P-selectin expression following TRAP-6 stimulation fold change of old relative to young platelets. **I.** Quantification of ATP release following TRAP-6 stimulation fold change of old relative to young platelets. Quantification of eicosanoid synthesis following collagen stimulation fold change of old relative to young platelets: **J** thromboxane B_2_; **K** prostaglandin E_2_ (PGE_2_); **L** 12-hydroxyeicosatetraenoic acid (12-HETE). Data presented as mean±SEM (*p<0.05, **p<0.01, ***p<0.005, ****p<0.001, n=3-8).

## Discussion

Changes in the age profile of platelet populations have been associated with many disease states and increased risk of thrombotic events. However, to date there has been very limited systematic investigation into the changes that platelets undergo as they age naturally. To address this need we have isolated, phenotyped and functionally characterized platelets as they normally age within the healthy human circulation.

To separate human platelets by age, we used TO fluorescence intensity as a surrogate marker of mRNA content, corroborating the relevance of this to platelet ageing by determining the levels of particular megakaryocyte-derived platelet-specific mRNAs with qRT-PCR. This approach was validated in mice by the use of temporal antibody labelling to show the same patterns of mRNA expression and TO intensity in young and old platelets, and is supported by recent research using nucleic acid dye staining with TO and SYTO13 to sort platelet subpopulations for transcriptomic analyses.^24,25^ We found highly significant correlations between the TO-determined platelet age and platelet levels of mRNA for ITGA2B, PF4, TUBB1, and between TO-determined platelet age and total platelet protein; the former being a log2 function and the latter a linear relationship.

It was particularly notable that as platelets aged there was a marked reduction in total protein content, such that old platelets had lost almost half of the protein present in young platelets. As platelets are anucleate, with a limited translational capacity, this is perhaps unsurprising and could be explained by protein degradation as a result of basal cellular processes and the release of proteins, perhaps encapsulated within platelet microvesicles.^7,26–29^ While our data show very clearly that there is a large general loss of proteins, this does not mean that individual proteins are not more subtly regulated. Platelets have the capacity to replenish their protein levels through endocytosis via the open canalicular system, as well as through limited protein synthesis.^30,31^ The latter of these processes has been shown to be engaged in response to stimulation, suggesting a controlled mechanism of synthesis to support the increased demand on platelets during the activation process, that might help counter the general protein loss. ^27,32^ Given the rate of mRNA decay in platelets, it can be speculated that ongoing protein synthesis may occur primarily within the first few days of platelet lifespan.^9^

In our work, we were able to identify 583 proteins in our proteomic analysis and establish significant differences in the levels of 78 proteins between young and old platelets. These differences were determined against the background of a general loss of proteins noted above and so demonstrate particular variations rather than general changes. There were 64 proteins that were significantly reduced in old platelets when compared to young platelets, indicating even greater loss during ageing. A further 14 proteins were proportionately more abundant in old platelets compared to young platelets, indicating selective retention, synthesis or uptake from the circulation. Among the proteins which demonstrated accelerated reduction in old platelets there was a particular signal for those involved in dynamic processes, such as in mitochondria (12 proteins) and the cytoskeleton (6 proteins).^4,33^ Ingenuity pathway analyses of the significantly modulated proteins predicted strong associations between older platelet age and reductions in cell activation and calcium flux, cytoskeletal organization, microtubule dynamics, formation of filaments and lamellipodia, binding of platelets and haemostasis, which we substantiated in the functional studies discussed below. The pathway analysis also associated old platelets to reduced cell viability and increased necrosis, apoptosis, and senescence.

Platelets are highly metabolically active cells, containing a small number of mitochondria, which under basal conditions generate approximately 40% of their ATP through oxidative phosphorylation.^34,35^ Following stimulation the energy demand on the platelet increases, causing an enhancement of oxidative phosphorylation.^36^ Our analyses demonstrated that the loss of multiple mitochondrial and metabolic proteins was associated with more than a halving of the number of mitochondria in old platelets, lessening their metabolic capacity and contributing to their reduced haemostatic function. The mechanisms governing the reduction in mitochondria number across platelet lifespan remain unclear, but recent research has highlighted the importance of the inner mitochondrial membrane mitophagy receptor, prohibitin 2, in mediating mitophagy in platelets. Interestingly, we identified a significant reduction in the abundance of prohibitin 2 in old platelets, suggesting accelerated degradation which may be as a result of an increase in mitophagy pathways and thus accelerated consumption.^37^

Cytoskeletal rearrangement is intimately linked to activation and adhesion pathways, thus the reduction in cytoskeletal proteins such as emerin, gelsolin, and twinfilin-2 identified via proteomics, coupled with the decline in α-tubulin and β-actin demonstrated by immunofluorescence and western blotting, will cause impairments in activation and adhesion pathways in old platelets. During platelet activation the membrane-bound actin network is severed, promoting membrane expansion and the formation of protrusions.^38^ The subsequent assembly of actin monomers allows the protrusions to elongate forming filopodia and lamellipodia, facilitating firm adhesion.^39^ Thus, with a reduction in the content of a number of cytoskeletal-associated proteins, the dynamic cycling of actin and tubulin will be diminished in old platelets. Indeed, in line with the loss of the key cytoskeletal proteins, we observed that old platelets do not fully spread, arresting at the filopodia stage of adhesion, consistent with defects in the later stages of actin assembly.

In addition to a weakened adhesion capacity, proteomics highlighted reductions in granule proteins and those involved in granule exocytosis suggesting there may be defects in granule secretion associated with ageing. Indeed, we found that old platelets have diminished dense and α-granule secretion upon activation, consistent with reports of higher P-selectin expression on activated reticulated platelets.^40^ Our study indicates two contributing changes in old platelets. Firstly, old platelets have reduced intracellular P-selectin levels and secondly, loss of cytoskeletal proteins together with decreases in SNAP23, part of the SNARE-fusion machinery, may cause a reduction in granule exocytosis.^41^

Reductions in cytoskeletal and mitochondrial proteins may also influence apoptotic pathways. Intrinsic apoptosis initiated by disruption to mitochondrial integrity has been described in platelets, highlighting the importance of mitochondrial membrane depolarization along with BCL-X_L_ and Bax/Bak proteins in mediating mitochondrial damage, subsequent caspase activation and phosphatidylserine exposure.^42–46^ Our data suggest that mitochondrial loss during normal platelet ageing may be due to an accumulation of damage triggering mitophagy pathways to remove damaged mitochondria. A key step within the mitophagy pathways is depolarization of the mitochondrial membrane, which may subsequently cause release of pro-apoptotic proteins into the platelet cytoplasm and promote the exposure of phosphatidylserine on the platelet surface.^2,47,48^ This exposure of phosphatidylserine would be enhanced by the loss of cytoskeletal proteins which normally maintain the asymmetric distribution of lipids within the platelet plasma membrane.^49–51^ With the degradation of cytoskeletal proteins, an imbalance in the membrane lipid asymmetry can arise facilitating the flipping of phosphatidylserine onto the outer leaflet.^52^ Exposed phosphatidylserine has been shown to be a key component of programmed cell death in platelets, acting as an ‘eat-me’ signal, triggering the clearance and destruction of old platelets from the circulation.^44,47,53^ Interestingly, platelet-specific knockout of the actin binding protein, twinfilin-2, a protein we identified in higher abundance in young platelets, results in accelerated platelet turnover due to increased clearance in the spleen, supporting a potential role for the cytoskeleton and cytoskeletal-associated proteins in the maintenance of platelet lifespan.^54^

Previous research into platelet ageing has produced contradictory evidence as to whether platelet size changes with age, with some groups suggesting platelet size to be independent of age whilst others report an association between mean platelet volume and thrombotic risk.^55–61^ Indeed, the use of TO to identify platelets containing elevated levels of mRNA as ‘young’ has been questioned on the basis that larger platelets may uptake more dye and so skew subsequent analysis.^62^ However, this cannot explain our data in healthy individuals as we noted a log2 relationship between particular megakaryocytic mRNAs and TO-determined platelet age rather than a linear one; i.e. platelets would have to vary in size by around 32-64 fold for such a relationship to hold. Furthermore, we found no differences in the cross-sectional areas of young and old platelets. We would emphasize, however, that we have not examined associations between platelet age and size in any pathological conditions in which different relationships may well exist. Interestingly, it was hypothesized some 40 years ago that buoyant density is a more accurate indicator of platelet age than size, with the observation that high density platelets are more metabolically active.^63^ This idea is supported by our data, since general loss of protein content with no change in size will cause older platelets to have lower buoyant densities, while young platelets with higher protein content contained more mitochondria consistent with higher metabolic activity. Consistent with this, our data also support the related and much more recent suggestion that mitochondria number is a better indicator of platelet age than platelet size.^64^

The discussion above has focused upon proteins that are decreased at accelerated rates as platelet age, however our proteomic analysis also demonstrated significant increases in the relative levels of a number of proteins in old platelets. These can largely be categorized as circulating proteins, including complement proteins and fibrinogen, which are most likely being bound and endocytosed by platelets as they circulate. Supporting this notion, research has implicated anionic phospholipids, such as phosphatidylserine, as promotors of complement protein activation.^65,66^ Interestingly, however, recent transcriptomic research has indicated old platelets have an increase in complement C5 transcripts compared to young platelets, therefore the observed upregulation in complement proteins may indeed be due to protein synthesis.^25^

In conclusion, our work demonstrates the changes in total and relative protein content as platelets age and the associated alterations in haemostatic function. We propose that young platelets are rapid haemostatic responders, due to their more robust cytoskeleton and higher mitochondria number, forming the core of aggregates and recruiting older platelets to the periphery. On the other hand, old platelets have blunted haemostatic responses, accumulate circulating proteins, and bear the indicators of cells marked for clearance from the circulation. Our research provides a detailed characterization of protein and functional changes as platelets normally age within the circulation providing key information for studies of platelet function in health and disease.

## Materials and Methods

### Ethical statement: Murine studies

Animal procedures were conducted under UK Home Office project licence authority (PPL/8422) in accordance with “The Animals (Scientific Procedures) Act 1986”, EU directive 2010/63/EU, and were subject to local approval from Queen Mary University of London and Imperial College London Ethical Review Panel.

### *In vivo* labelling and flow cytometric sorting of murine young and old platelets

Male C57Bl/6 wild-type (WT) mice were purchased from Charles River UK. All mice were 8-12 weeks old (20-25g) and housed for a minimum of 7 days before commencement of experiments. They were housed on a 12-hour light-dark cycle, at a temperature of 22 to 24°C with access to water and food *ad libitum*. Injectable fluorescently conjugated anti-platelet antibodies (anti-CD42c DyLight-x488 or x649, Emfret) were administered (intravenous; i.v.) to C57BL/6 mice as per supplier guidance. Briefly, anti-CD42c-x488 was injected at 0 hours, followed by anti-CD42c-x649 at 23 hours later. Blood was collected from the inferior vena cava into sodium citrate (0.32%; Sigma) from mice anesthetized with ketamine (Narketan^®^, 100 mg/kg; Vetoquinol) and xylazine (Rompun^®^, 10 mg/kg; Bayer) and PRP was isolated as previously published.^67^ Briefly, whole blood was diluted 1:1 with HEPES-tyrodes buffer (37mM NaCl, 20mM HEPES, 5.6mM glucose, 1 g/l BSA, 1mM MgCl_2_, 2.7mM KCl, 3.3mM NaH_2_PO_4_) before centrifugation (100 *x g*, 8 minutes).

Murine platelets were sorted as previously stated for human platelets; old platelets were gated as CD42c-x488/CD42c-x649 dual positive, and young platelets as CD42c-x649 positive/-x488 negative events. Subsequently, the platelets were pelleted at 1000 *x g* for 10 minutes in the presence of prostacyclin (PGI_2_, 2μM) and re-suspended in 500μl of Qiazol (QIAGen).

### RNA extraction, cDNA synthesis and qRT-PCR of sorted murine platelets

RNA was extracted from sorted platelet pellets (2.5 million platelets per population) using the miRNAEasy mini kit (QIAGen), following manufacturer’s instructions. cDNA was synthesized from the extracted platelet RNA using VILO SuperScript for RT-PCR (Life Technologies). Target genes were rationally selected as candidates based on their specificity to platelets; TaqMan anti-mouse FAM-labelled probes were obtained from Fisher Scientific; Integrin alpha-IIb (*ITGA2B*; Mm00439741_m1), platelet factor 4 (*PF4*; Mm00451315_g1), tubulin beta-1 chain (*TUBB1*; Mm01239914_g1). qPCR was performed on a ViiA 7 Real-Time PCR System (Applied Biosystems). Data was analyzed using ViiA 7 Software (Applied Biosystems)

### Ethical statement: Human studies

All studies were conducted according to the principles of the Declaration of Helsinki and approved by St Thomas’s Hospital Research Ethics Committee (Ref. 07/Q0702/24). All volunteers were screened prior to entering the study to confirm their platelet count was in the normal range and subsequently gave written informed consent.

### Blood collection and isolation of platelets

Blood was collected by venepuncture into tri-sodium citrate (3.2%; Sigma) from healthy volunteers (aged 25-40, 60% female), who had abstained from non-steroidal anti-inflammatory drug consumption for the preceding 14 days. Platelet rich plasma (PRP) was obtained by centrifugation of whole blood (175 *× g,* 15 minutes, 25°C) and samples were processed immediately following collection.

### Thiazole orange staining and flow cytometric sorting of platelet subpopulations

Thiazole orange (200ng/mL, Sigma) was incubated with platelet rich plasma (PRP) at room temperature for 30 minutes. Thiazole orange stained PRP was diluted 1 in 8 using (i) saline with apyrase (0.04U/mL, Sigma), prostaglandin E_1_ (PGE_1_, 2μmol/L, Sigma) and phosphodiesterase inhibitor 3-isobutyl-1-methylxanthine (IBMX, 0.5mM; Sigma) for quantitative real time polymerase chain reaction (qRT-PCR) experiments or (ii) modified Tyrode’s *N*-2-hydroxyethylpiperazine-*N*’-2-ethanesulfonic acid (HEPES) buffer (134mmol/L NaCl, 2.9mM KCl, 0.34mmol/L Na_2_HPO_4_, 12mmol/L NaHCO_3_, 20mmol/L HEPES and 1mmol/L MgCl_2_; pH 7.4; Sigma), with glucose (0.1% (w/v); Sigma), apyrase (0.02U/mL) and PGE_1_ (2μM).

Platelets were sorted using a BD FACS Aria IIIu Fusion Cell Sorter (70μm nozzle, 70 Ps; BD Bioscience) and gated according to thiazole orange fluorescence intensity (Figure 1A). Sorting efficiency was maintained above 85% and the purity of samples was confirmed following sorting. Subsequently, the platelets were pelleted at 1000 *x g* for 10 minutes in the presence of prostacyclin (PGI_2_, 2μmol/L; Tocris) and re-suspended in modified Tyrode’s HEPES (MTH) buffer, supplemented with calcium chloride (CaCl_2_, 2mmol/L; Sigma).

### RNA extraction, cDNA synthesis and qRT-PCR of sorted platelets

RNA was extracted from platelet pellets (10 million per population) using the RNeasy Mini kit according to manufacturer’s instructions (Qiagen). cDNA was synthesized from extracted platelet RNA using SuperScript^®^ III first-strand synthesis system for RT-PCR (Life Technologies). TaqMan anti-human FAM-labelled probes were obtained from Life Technologies; Integrin alpha-IIb (*ITGA2B*; Hs01116228_m1), platelet factor 4 (*PF4*; Hs00427220_g1), tubulin beta-1 chain (*TUBB1*; Hs00258236_m1). qRT-PCR was performed on an Applied Biosystems ABI 7900HT (Life Technologies) instrument and data was collected using SDS software (Life Technologies).

### Flow cytometric measurement of activation markers pre- and post-sorting

PRP was diluted in modified Tyrode’s HEPES buffer (1 in 8), supplemented with PGE_1_ (2μM) and subsequently sorted using a BD FACS Aria IIIu Fusion Cell Sorter (70μm nozzle, 70 Ps). The platelets were gated on forward scatter (FSC) and side scatter (SSC) and 5 million platelets were sorted. To assess the activation potential of pre- and post-sorted platelets, 20μl of pre- or post-sorted platelets were diluted in 20μl annexin binding buffer, incubated with phosphate buffered saline (PBS), Thrombin Receptor Activator Peptide-6 (TRAP-6; 30μM) or Adenosine Diphosphate (ADP; 30μM) along with CD42b-BV421 (1:60, Biolegend), PAC-1 FITC (1:12, D Bioscience) and annexin V APC (1:60, Biolegend) for 30 minutes. The samples were further diluted in 100μl of filtered PBS and subsequently analyzed on the ACEA Novocyte 3000 (ACEA Biosciences Inc.), collecting a minimum of 5,000 CD42b-BV421 positive events. Analysis was performed using NovoExpress 1.3.0 (ACEA Biosciences Inc.)

### Protein extraction and proteomic analysis of sorted human platelets

Protein was extracted from sorted platelet subpopulations pellets (65 million platelets per population) using lysis buffer (100mM TRIS pH 7.5, 2% sodium dodecyl sulfate (SDS), 1 cOmplete™ Mini Protease Inhibitor Cocktail Tablet per 10ml; all Sigma). The protein content of the sorted platelet subpopulations was measured using a Nanodrop Spectrophotometer ND-1000 (Thermo Fisher Scientific) and samples were stored at −80°C. Proteomic analysis was performed on 30μg of total protein in each platelet subpopulation.

Proteomic analysis of the subpopulations was performed on tryptic digests obtained using the Filter Aided Sample Preparation protocol using 30k filter units (Microcon YM-30, Millipore) and sequencing grade trypsin (Trypsin Gold, Promega) with an enzyme to protein ratio of 1:50.^68^ The concentration of tryptic peptides was estimated by UV spectrometer at 280 nm, and 10μg peptides were used for mass spectrometry analysis. C18 stage tips were used to desalt and purify peptides as previously described.^69^ After elution from stage tips the samples were evaporated in a CentriVap Concentrator for approximately 10-15 minutes, until approximately 5μl volume remained. The sample was then resuspended in 18μl of 2% acetonitrile/0.5% acetic acid and stored at 4°C until mass spectrometry analysis.

LC-MS/MS analysis was performed using an Ultimate3000 nano-LC system coupled to a hybrid quadrupole-orbitrap mass spectrometer (Q Exactive, Thermo Fisher Scientific). Peptides were separated using an increasing acetonitrile gradient from 2% to 33% in a linear LC gradient of 40 minutes on a C18 reverse phase chromatography column packed with 2.4μm particle size, 300 Å pore size C18 material (Dr. Maisch GmbH, Ammerbuch-Entringen) to a length of 120mm in a column with a 75μm ID, using a flow rate of 250nL/min. Data was acquired with the mass spectrometer operating in automatic data-dependent acquisition mode (DDA, shotgun). A full MS service scan at a resolution of 70,000, AGC target 3e6 and a range of m/z 350–1600 was followed by up to 12 subsequent MS/MS scan with a resolution of 17,500, AGC target 2e4, isolation window m/z 1.6 and a first fix mass of m/z 100. Dynamic exclusion was set to 40 seconds.

Raw mass spectrometry data files were processed in the MaxQuant software (v.1.3.0.547). MS/MS spectra were searched using the built-in search engine Andromeda against a human FASTA Uniprot database (release 2016_3). Trypsin/P was set as a protease with up to two missed cleavages allowed. Proteins were quantified by a minimum of 2 peptides and the maximum mass deviation tolerance in MS mode was set to 20 ppm for the initial search and 6 ppm for the main search, whereas the maximum deviation tolerance in MS/MS mode was set to 20 ppm. Thiomethylation of cysteine residues was set as fixed modification, while oxidation of methionine and N-terminal protein acetylation were selected as variable modifications. The false discovery rate (FDR), determined by reverse database searching, was set to 0.01 for the peptides and proteins. All other settings were kept at their default values. Label-free quantification was performed using the MaxLFQ engine integrated in MaxQuant. Statistical analysis was performed with Perseus software (Version 1.6.0.7).^70^ Only proteins present in at least 50% of the samples in at least one group of differently aged platelets were considered identified. The filtered, log-transformed label-free quantification intensity data was subsequently median normalised and missing values imputed with a downshifted normal distribution with Perseus. Differentially modulated proteins were identified by ANOVA (p<0.05) and subsequently a two-tailed, two-sample t-test analysis (p value<0.05). The results were subsequently filtered using the Benjamini-Hochberg procedure for FDR correction (FDR<0.05) using the built-in volcano plot function in Perseus. Proteins found to be differentially expressed between groups (p<0.05, FDR 0.05) were subjected to pathway mapping analysis and were distributed into categories according to their cellular component and biological process using Ingenuity Pathway Analysis (IPA; Qiagen). The mass spectrometry proteomics data have been deposited to the ProteomeXchange Consortium via the PRIDE partner repository with the dataset identifier PXD014490.^71^

### Immunofluorescence and confocal microscopy

Paraformaldehyde (PFA, 4%; VWR) fixed platelet subpopulations were centrifuged onto poly-l-lysine coverslips (VWR; 600 *x g*, 5 minutes), permeabilized with 0.2% Triton, 2% donkey serum, 1% bovine serum albumin (BSA) in phosphate buffered saline (PBS, all Sigma) followed by incubation with primary antibodies; mouse monoclonal anti-α-tubulin (1:200; Sigma) or rat monoclonal anti-α-tubulin (1:200; Invitrogen), mouse monoclonal anti-TOM20 (1:500; Santa Cruz), rabbit polyclonal anti-ERp57 (1:200; Abcam), rabbit monoclonal anti-P-selectin, rabbit polyclonal anti-fibrinogen (1:100; Thermo Fisher Scientific), rabbit monoclonal anti-complement C4 (1:200; Abcam), then secondary conjugated antibodies; anti-mouse Alexa Fluor 555 or anti-mouse Alexa Fluor 647, anti-rat Alexa Fluor 594, anti-rabbit Alexa Fluor 647 (1:500; Thermo Fisher Scientific) or Phalloidin-647 (1:100; Thermo Fisher Scientific). Samples were mounted with Prolong Diamond antifade mount (Thermo Fisher Scientific). Confocal microscopy was performed using an inverted Zeiss LSM880 with Airyscan confocal microscope; 63x objective, 1.4 Oil DICII (Zeiss). Analysis was conducted using Zen Software (2.3 SP1, Zeiss) and ImageJ (NIH). Mitochondria number and area were calculated using Image J, averaging the number and size of mitochondria of 5 fields of view per individual. Integrated fluorescence intensity was established using at least 5 fields of view, averaged from a minimum of 20 platelets per individual.

### Western blotting

Platelet subpopulations pellets (1.5 × 10^6^) were lysed with 2% SDS, supplemented with Laemmli buffer containing β-mercaptoethanol, boiled at 95°C for 5 minutes and separated by SDS-PAGE (10% gel), before transferring to nitrocellulose membranes. Subsequently, membranes were incubated with 5% skimmed milk in PBS supplemented with 0.05% (v/v) Tween 20 (Sigma, PBS-T) for 30 minutes at room temperature (RT), followed by incubation with primary antibodies; anti α-tubulin (1:10,000; Sigma) and anti β-actin (1:10,000; Santa Cruz Biotechnology), at 4°C overnight. Membranes were then washed with PBS-T and incubated with anti-mouse IgG, horseradish peroxidase-linked antibody (1:10,000, Cell Signaling Technology) for 1 hour, RT. After washing with PBS-T and PBS, membranes were developed using ECL reagent (Merck™ Immobilon™ Western Chemiluminescent HRP Substrate, Thermo Fisher Scientific), X-ray films (Scientific Laboratory Supplies Ltd) and a film processor. Densitometry was performed using Image J software.

### Quantification of mitochondrial membrane potential and phosphatidylserine exposure

Platelet subpopulations (3 × 10^6^) stained with CD42b-BV421 (1:300; Biolegend) in (i) 300μl MTH buffer along with MitoProbe™ DiIC_1_(5) (1:300; Invitrogen) or (ii) 300μl annexin V binding buffer stained with Annexin V-APC (1:300; Biolegend) for 20 minutes. MitoProbe™ DiIC_1_(5) and Annexin V-positive events were determined by flow cytometry (BD LSRII; BD Biosciences) and analyzed using FlowJo software v.10 (TreeStar Inc).

### Functional characterization of platelet subpopulations

#### Platelet adhesion and spreading

Platelet subpopulations (2 × 10^6^) were incubated on Horm collagen type I (100μg/ml; Takeda) or fibrinogen (100μg/ml, Sigma) coated coverslips for 90 minutes at 37°C. Adherent platelets were fixed with paraformaldehyde (0.2%) for 10 minutes, permeabilized with Triton (0.2%) for 5 minutes, stained with Phalloidin-647 (1:100) and mounted with Prolong Diamond antifade mount. Imaging was performed using an inverted Zeiss LSM880 with Airyscan confocal microscope; collecting at least 10 fields of view per sample, averaging at least 30 platelets from 4 individuals. Analysis was conducted using Zen Software and ImageJ; spreading stage was quantified as; i: adherent, not spread; ii: filopodia; iii: lamellipodia; iv: fully spread.

#### Aggregation of platelet subpopulations

Sorted platelets (2.5 × 10^6^) were stained with CD61-FITC or CD61-APC (young and old; 1 in 100, eBioscience) and centrifuged (1000 *x g*, 10 minutes). The pellets were re-suspended in MTH buffer and recombined in equal proportions. Aggregation was stimulated with TRAP-6 (25μM) in a 96-well plate (15 minutes, 750rpm, 37°C; BioShake IQ, Quantifoil Instruments) and fixed with 1% formalin (Sigma). Imaging flow cytometry (ImageStream^X^ Mark II; Amnis) was used to measure single platelets and formed aggregates. Image analysis was performed using IDEAS software (Amnis).

#### Quantification of calcium dynamics

Platelet subpopulations (1 × 10^7^/ml) were incubated with Calbryte 630^TM^ (2μM; Stratech) for 30 minutes at 37°C followed by staining with CD42b-BV421 (1:100; Biolegend) for 15 minutes at room temperature and supplementation with 2mM CaCl_2_. Baseline Calbryte 630^TM^ fluorescence was recorded for 30 seconds, followed by challenge with TRAP-6 (25μM) or ionomycin (10μM; Invitrogen) and subsequent recording for 2 minutes. Samples were acquired on a BD LSRII using FACSDiva acquisition software, and analyzed using FlowJo software v.10.

#### Quantification of platelet activation

Platelet subpopulations (1 × 10^7^/ml) were incubated with TRAP-6 (25 μM) or PBS (20 minutes, 37°C), stained with anti-human CD61-FITC and CD62P-APC (1:180; eBioscience) and fixed with 1% formalin. CD62P-postive events were determined by flow cytometry (FACS Calibur, BD Bioscience) using CellQuest Software and analyzed using FlowJo software v.8.

#### Quantification of ATP release

Sorted platelets (2.5 × 10^6^, 45μl MTH buffer) supplemented with fibrinogen (1mg/mL) were incubated with TRAP-6 (25μM) for 2 minutes, 1200rpm, 37°C) followed by 25μL of Chronolume luciferin-luciferase system reagent (Chrono-log) for 2 minutes (350rpm). ATP release was determined by assessing the luminescence emitted using a Tecan Infinite^®^ M200 plate reader (Tecan) and comparing to standard ATP concentrations.

#### LC-MS/MS analysis of eicosanoid release

Eicosanoid production was measured from the releasate of platelet subpopulations (1 × 10^7^/mL) supplemented with fibrinogen (1mg/mL) following incubation (30 minutes, 37°C) with PBS or collagen (30μg/mL, Takeda). LC-MS/MS was performed at the National Institute of Environmental Health Science, North Carolina, USA as described previously.^6^

## Statistical analyses

Data were expressed as mean±SEM. Graphs and statistical analysis were generated using GraphPad Prism 8 (GraphPad Software Inc.). Statistical analyses were performed with a paired t-test or a one-way ANOVA with Tukey’s post-test for multiple comparisons. Correlations were assessed by simple linear regression. Significance was defined as p<0.05.

## Acknowledgements

The authors would like to acknowledge the support the Flow Cytometry Core Facilities at the Blizard Institute and Charterhouse Square, Queen Mary University of London and the UCD Conway Institute mass spectrometry facilities.

