## Supplemental Figures for "Proteome and functional decline as platelets age in the circulation"

### Supplement Table 1: mRNA expression in thiazole orange sorted human platelets

| Gene | Cycle Threshold | | | | | |
|---|---|---|---|---|---|---|
| | Young | | Intermediate | | Old | |
| | Mean | SEM | Mean | SEM | Mean | SEM |
| **ADCY3** | 35.45 | 0.35 | 40.00 | 0.00 | 40.00 | 0.00 |
| **GP6** | 32.78 | 0.24 | 39.50 | 0.50 | 39.46 | 0.54 |
| **GUCY1A3** | 32.58 | 0.48 | 39.51 | 0.49 | 40.00 | 0.00 |
| **P2RY1** | 35.75 | 0.39 | 39.19 | 0.81 | 39.17 | 0.83 |
| **P2RY12** | 31.76 | 0.30 | 36.72 | 0.70 | 39.35 | 0.65 |
| **PEAR1** | 36.13 | 0.23 | 40.00 | 0.00 | 40.00 | 0.00 |
| **PLA2G4A** | 36.96 | 0.03 | 40.00 | 0.00 | 40.00 | 0.00 |
| **PTGIR** | 34.08 | 0.40 | 39.50 | 0.50 | 40.00 | 0.00 |
| **PTGS1** | 30.82 | 0.32 | 37.38 | 0.86 | 39.31 | 0.69 |
| **TBXAS1** | 34.60 | 0.71 | 39.38 | 0.62 | 40.00 | 0.00 |

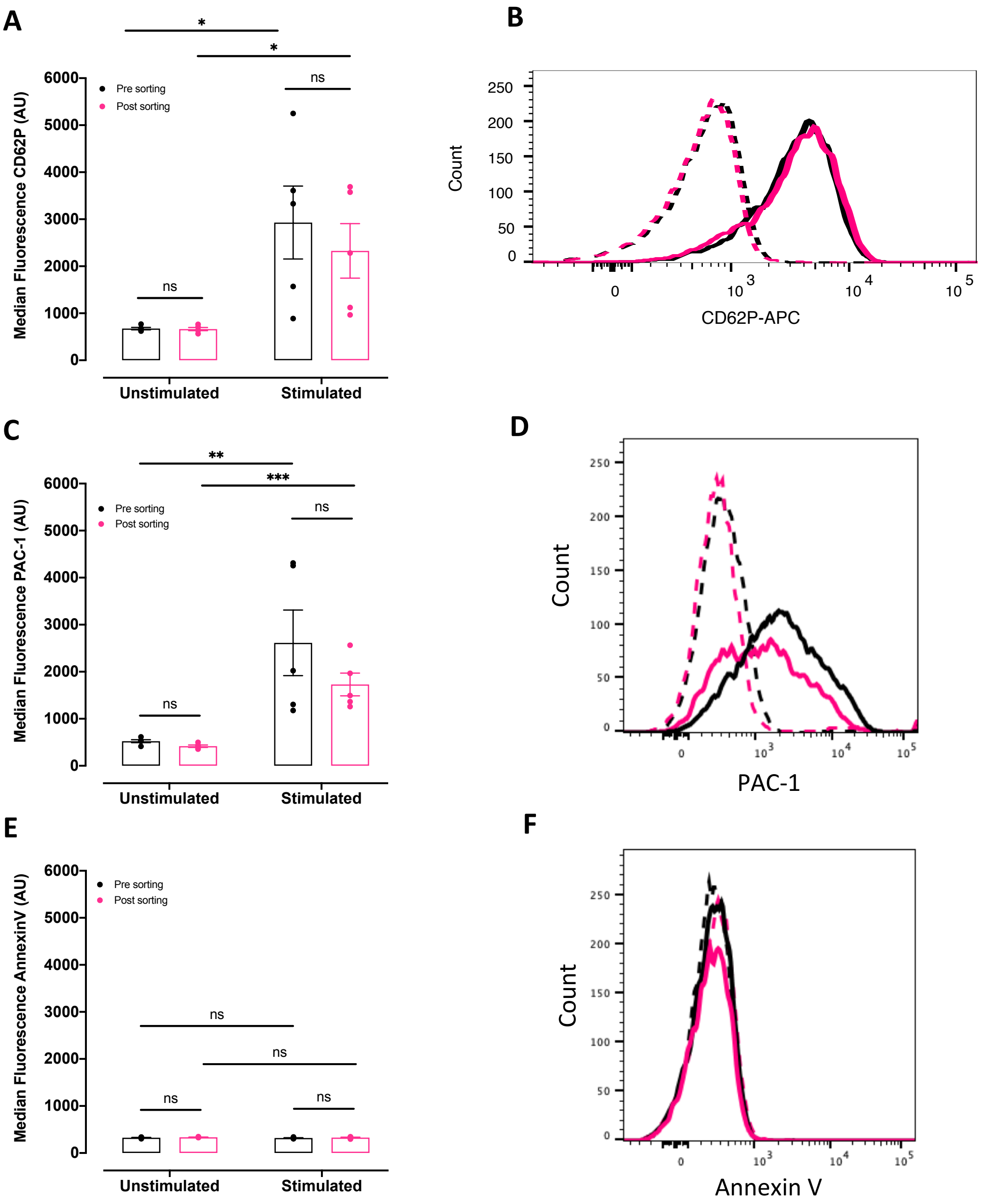


**Supplemental Figure 1: Expression of P-selectin, PAC-1 binding and Annexin V pre- and post-sorting**

**A.** Flow cytometric measurement of P-selectin in pre- (black) and post-sorted (pink) platelets under basal conditions and following stimulation with TRAP-6. **B.** Representative histograms of P-selectin expression under basal (dashed lines) and stimulated conditions in pre- (black) and post-sorted (pink) platelets. **C.** Flow cytometric measurement of PAC-1 binding in pre- (black) and post-sorted (pink) platelets under basal conditions and following stimulation with ADP. **D.** Representative histograms of PAC-1 binding under basal (dashed lines) and stimulated conditions in pre- (black) and post-sorted (pink) platelets. **E.** Flow cytometric measurement of Annexin V binding in pre- (black) and post-sorted (pink) platelets under basal conditions and following stimulation with ADP. **F.** Representative histograms of Annexin V binding under basal (dashed lines) and stimulated conditions in pre- (black) and post-sorted (pink) platelets Data presented as mean±SEM ($p>0.05$, $n=5$).

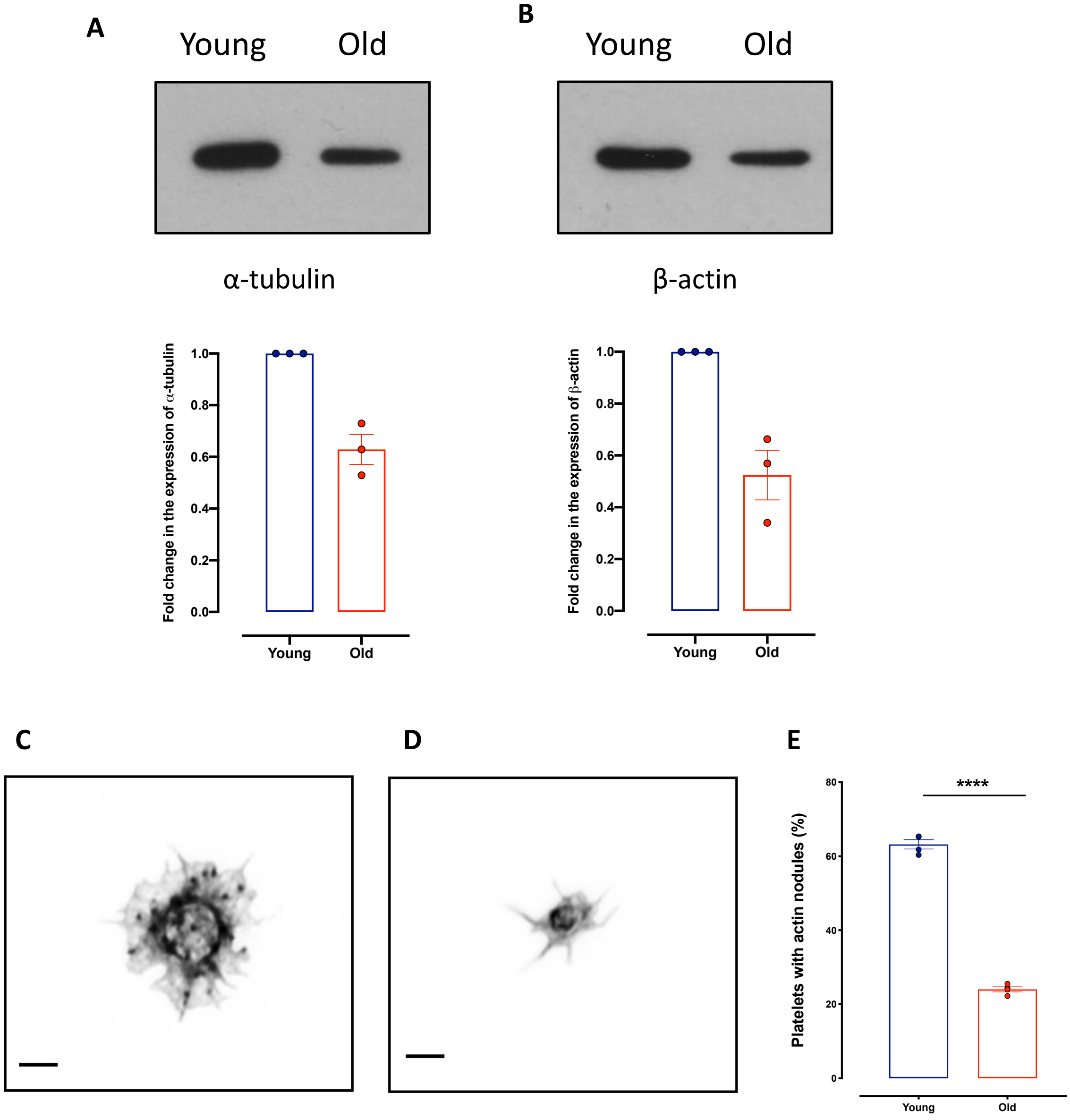


**Supplemental Figure 2: Analysis cytoskeletal structure and characteristics**

**A** and **B.** Representative western blots from lysates of 1.5 million young and old platelets analyzed using (**A**) α-tubulin and (**B**) β-actin antibodies, graphs indicate results from densitometry analysis; mean±SEM, n=3 independent lysates, expressed as fold change of protein levels in young platelets. **C and D.** Representative confocal microscopy images showing actin nodules in (**C**) young and (**D**) old platelets spread on fibrinogen. **E.** Quantification of the percentage of young and old platelets containing actin nodules. Data presented as mean±SEM (n=4, *p<0.05).
