## Supplemental Table 2 for "Proteome and functional decline as platelets age in the circulation"

Supplemental Table 2: Analysis of the proteome of young, intermediate and old platelets

| Gene Name | Protein ID | Protein | Protein Name | Young Z score | Intermediate Z Score | Old Z Score | Peptides | Razor + unique peptides | Unique peptides | Sequence coverage [%] | Unique + razor sequence coverage [%] | Unique sequence coverage [%] | Mol. weight [kDa] | Intensity | MS/MS count |
|---|---|---|---|---|---|---|---|---|---|---|---|---|---|---|---|
| A1BG | M0R009 | M0R009 | Alpha-1B-glycoprotein | -0.67424125 | -0.051267175 | 0.725508133 | 5 | 5 | 5 | 18 | 18 | 18 | 33.341 | 48636000 | 16 |
| A2M | P01023 | A2MG | Alpha-2-macroglobulin | -0.03147375 | -0.05733985 | 0.08881335 | 37 | 37 | 33 | 30.7 | 30.7 | 28 | 163.29 | 1518800000 | 210 |
| ABHD16A | O95870 | ABHGA | Phosphatidylserine lipase ABHD16A | 0.450428825 | 0.206688143 | -0.6571167 | 7 | 7 | 7 | 14.2 | 14.2 | 14.2 | 63.243 | 145530000 | 21 |
| ACADVL | P49748 | ACADV | Very long-chain specific acyl-CoA dehydrogenase, mitochondrial | 0.8999098 | -0.180869125 | -0.71904055 | 20 | 20 | 20 | 36.9 | 36.9 | 36.9 | 70.389 | 490210000 | 63 |
| ACLY | P53396 | ACLY | ATP-citrate synthase | -0.236072675 | 0.494700125 | -0.258627575 | 19 | 19 | 19 | 22.4 | 22.4 | 22.4 | 120.84 | 363230000 | 54 |
| ACO2 | Q99798 | ACON | Aconitate hydratase, mitochondrial | 0.267871475 | -0.198091675 | -0.069779713 | 15 | 15 | 15 | 21.9 | 21.9 | 21.9 | 85.424 | 200510000 | 37 |
| ACP1 | G5E9R5 | G5E9R5 | Acid phosphatase 1, soluble, isoform CRA_d | -0.45618505 | 0.685382475 | -0.229197665 | 3 | 3 | 2 | 38.8 | 38.8 | 27.5 | 8.9711 | 103510000 | 13 |
| ACTA1 | P68133 | ACTS | Actin, alpha skeletal muscle | 0.3085701 | -0.11016835 | -0.198401925 | 26 | 1 | 1 | 37.1 | 4.2 | 4.2 | 42.051 | 30878000000 | 115 |
| ACTB | P60709 | ACTB | Actin, cytoplasmic 1 | -0.430812 | 0.25788315 | 0.172928875 | 43 | 43 | 2 | 84 | 84 | 7.2 | 41.736 | 6.166E+11 | 3121 |
| ACTN1 | P12814 | ACTN1 | Alpha-actinin-1 | 0.8861307 | -0.26845555 | -0.617675135 | 75 | 75 | 6 | 81.8 | 81.8 | 10.2 | 103.06 | 69181000000 | 1534 |
| ACTN4 | O43707 | ACTN4 | Alpha-actinin-4 | 0.709413875 | -0.128750075 | -0.58066385 | 63 | 41 | 40 | 72.2 | 51.4 | 50.2 | 104.85 | 3308500000 | 242 |
| ACTR2 | P61160 | ARP2 | Actin-related protein 2 | 0.532034275 | 0.114199575 | -0.64623387 | 18 | 18 | 18 | 44.4 | 44.4 | 44.4 | 44.76 | 3132400000 | 142 |
| ACTR3 | P61158 | ARP3 | Actin-related protein 3 | 0.457590675 | -0.0147095 | -0.442881375 | 24 | 24 | 19 | 60 | 60 | 53.6 | 47.371 | 2668900000 | 166 |
| AGT | P01019 | ANGT | Angiotensinogen | -0.0544833 | -0.121524025 | 0.176007473 | 6 | 6 | 6 | 14.6 | 14.6 | 14.6 | 53.154 | 126820000 | 24 |
| AHSG | P02765 | FETUA | Alpha-2-HS-glycoprotein | 0.224993623 | 0.195547875 | -0.420541525 | 5 | 5 | 5 | 13.4 | 13.4 | 13.4 | 39.324 | 1091200000 | 62 |
| AK2 | F8W1A4 | F8W1A4 | Adenylate kinase 2, mitochondrial | 0.617298825 | -0.6181465 | 0.00084785 | 8 | 8 | 8 | 43.1 | 43.1 | 43.1 | 25.63 | 239890000 | 34 |
| AKR7A2 | O43488 | ARK72 | Aflatoxin B1 aldehyde reductase member 2 | 0.415893575 | 0.328643025 | -0.7445367 | 10 | 10 | 10 | 32.3 | 32.3 | 32.3 | 39.589 | 580320000 | 49 |
| ALDOA | P04075 | ALDOA | Fructose-bisphosphate aldolase A | 0.871562675 | -0.517267 | -0.35429591 | 33 | 33 | 29 | 87.1 | 87.1 | 75.3 | 39.42 | 16830000000 | 473 |
| ALOX12 | P18054 | LOX12 | Arachidonate 12-lipoxygenase, 12S-type | -0.29725386 | -0.308164225 | 0.6054183 | 20 | 20 | 20 | 33.6 | 33.6 | 33.6 | 75.693 | 1384500000 | 129 |
| AMPD2 | Q01433 | AMPD2 | AMP deaminase 2 | 0.798481225 | -0.527212625 | -0.271268575 | 16 | 16 | 16 | 17.6 | 17.6 | 17.6 | 100.69 | 313530000 | 46 |
| ANO6 | Q4KMQ2 | ANO6 | Anoctamin-6 | 0.267376225 | -0.363251468 | 0.095875383 | 13 | 13 | 13 | 17.8 | 17.8 | 17.8 | 106.16 | 299280000 | 25 |
| ANXA11 | P50995 | ANX11 | Annexin A11 | 0.98433215 | -0.561897475 | -0.422434625 | 9 | 9 | 9 | 19.8 | 19.8 | 19.8 | 54.389 | 341560000 | 37 |
| ANXA7 | P20073 | ANXA7 | Annexin A7 | -0.469838825 | -0.4199708 | 0.889809425 | 12 | 12 | 12 | 28.3 | 28.3 | 28.3 | 52.739 | 368220000 | 40 |
| AP1B1 | Q10567 | AP1B1 | AP-1 complex subunit beta-1 | 0.6683743 | -0.7397916 | 0.0714177 | 14 | 14 | 8 | 16.3 | 16.3 | 9.8 | 104.64 | 319380000 | 49 |
| AP1M1 | Q9BXS5 | AP1M1 | AP-1 complex subunit mu-1 | -0.10860005 | 0.357240885 | -0.24864065 | 10 | 10 | 10 | 29.1 | 29.1 | 29.1 | 48.586 | 229150000 | 36 |
| APOA1 | P02647 | APOA1 | Apolipoprotein A-I | -0.518682253 | 0.459292048 | 0.0593904 | 35 | 35 | 33 | 83.9 | 83.9 | 83.9 | 30.777 | 24195000000 | 629 |
| APOA2 | V9GYE3 | V9GYE3 | Apolipoprotein A-II | -0.395393525 | 0.4268612 | -0.031467625 | 6 | 6 | 6 | 42.3 | 42.3 | 42.3 | 5.8767 | 2546800000 | 84 |
| APOA4 | P06727 | APOA4 | Apolipoprotein A-IV | -0.914874975 | 0.431210975 | 0.48366405 | 29 | 29 | 28 | 65.9 | 65.9 | 65.9 | 45.398 | 1536200000 | 183 |
| APOB | P04114 | APOB | Apolipoprotein B-100 | -0.8011828 | 0.21291305 | 0.5882697 | 141 | 141 | 141 | 33 | 33 | 33 | 515.6 | 3422500000 | 616 |
| APOC1 | K7ERI9 | K7ERI9 | Apolipoprotein C-I | -0.0202885 | -0.0554997 | 0.07578825 | 3 | 3 | 3 | 26 | 26 | 26 | 8.647 | 405640000 | 19 |
| APOC3 | P02656 | APOC3 | Apolipoprotein C-III | -0.314292925 | -0.026141615 | 0.3404347 | 4 | 4 | 4 | 37.4 | 37.4 | 37.4 | 10.852 | 1188900000 | 63 |
| APOD | P05090 | APOD | Apolipoprotein D | 0.122915523 | -0.127088925 | 0.004173425 | 4 | 4 | 4 | 21.2 | 21.2 | 21.2 | 21.275 | 194720000 | 27 |
| APOE | P02649 | APOE | Apolipoprotein E | -0.82052705 | 0.0607567 | 0.759770325 | 22 | 22 | 22 | 64 | 64 | 64 | 36.154 | 1006300000 | 134 |
| APOL1 | O14791 | APOL1 | Apolipoprotein L1 | -0.0102577 | -0.332855175 | 0.343112775 | 6 | 6 | 6 | 14.8 | 14.8 | 14.8 | 43.974 | 170810000 | 20 |
| APP | A0A0A0MRG2 | A0A0A0MRG2 | Amyloid-beta precursor protein | 0.1229219 | 0.245701225 | -0.36862325 | 13 | 13 | 12 | 22.1 | 22.1 | 20.9 | 75.109 | 497550000 | 51 |
| APRT | P07741 | APT | Adenine phosphoribosyltransferase | 0.259699 | -0.532604458 | 0.272905595 | 8 | 8 | 8 | 47.8 | 47.8 | 47.8 | 19.608 | 322140000 | 44 |
| ARCN1 | P48444 | COPD | Coatomer subunit delta | 0.405941575 | 0.020087253 | -0.4260291 | 11 | 11 | 11 | 24.3 | 24.3 | 24.3 | 57.21 | 341620000 | 41 |
| ARF1 | P84077 | ARF1 | ADP-ribosylation factor 1 | 0.8529179 | -0.6341586 | -0.218759458 | 11 | 11 | 6 | 64.1 | 64.1 | 29.8 | 20.697 | 4036600000 | 153 |
| ARF4 | P18085 | ARF4 | ADP-ribosylation factor 4 | 0.97043345 | -0.173234315 | -0.797198925 | 10 | 6 | 6 | 64.4 | 35.6 | 35.6 | 20.511 | 422650000 | 50 |
| ARHGAP1 | Q07960 | RHG01 | Rho GTPase-activating protein 1 | 0.901240775 | -0.54852825 | -0.35271245 | 18 | 18 | 18 | 47.4 | 47.4 | 47.4 | 50.435 | 1446500000 | 83 |
| ARHGAP18 | Q8N392 | RHG18 | Rho GTPase-activating protein 18 | 0.8584712 | -0.26395615 | -0.594514975 | 27 | 27 | 27 | 39.8 | 39.8 | 39.8 | 74.976 | 865290000 | 105 |
| ARHGAP45 | Q92619 | HMHA1 | Rho GTPase-activating protein 45 [Cleaved into: Minor histocompatibility antigen HA-1 | 0.204941075 | -0.042137575 | -0.162803275 | 20 | 20 | 20 | 20.4 | 20.4 | 20.4 | 124.61 | 445100000 | 47 |
| ARHGAP6 | O43182 | RHG06 | Rho GTPase-activating protein 6 | 0.509521775 | -0.051340625 | -0.4581813 | 20 | 20 | 20 | 25.4 | 25.4 | 25.4 | 105.95 | 242030000 | 40 |
| ARHGDIA | J3KTF8 | J3KTF8 | Rho GDP-dissociation inhibitor 1 | 0.44920715 | -0.413259675 | -0.035947475 | 9 | 9 | 9 | 34.7 | 34.7 | 34.7 | 21.517 | 900840000 | 51 |
| ARHGDIB | P52566 | GDIR2 | Rho GDP-dissociation inhibitor 2 | 0.6935724 | 0.281072875 | -0.97464545 | 12 | 12 | 12 | 70.6 | 70.6 | 70.6 | 22.988 | 2007800000 | 99 |
| ARL6IP5 | O75915 | PRAF3 | PRA1 family protein 3 | 0.338963071 | 0.57126825 | -0.9102312 | 8 | 8 | 8 | 32.4 | 32.4 | 32.4 | 21.614 | 1306500000 | 131 |
| ARPC1B | O15143 | ARC1B | Actin-related protein 2/3 complex subunit 1B | 0.65741812 | -0.244700465 | -0.412717475 | 15 | 15 | 15 | 43.3 | 43.3 | 43.3 | 40.949 | 2285400000 | 117 |
| ARPC2 | O15144 | ARPC2 | Actin-related protein 2/3 complex subunit 2 | -0.423754568 | 0.346104289 | 0.077650475 | 21 | 21 | 21 | 58.7 | 58.7 | 58.7 | 34.333 | 3336500000 | 182 |
| ARPC3 | O15145 | ARPC3 | Actin-related protein 2/3 complex subunit 3 | 0.59621685 | 0.142954315 | -0.7391712 | 9 | 9 | 9 | 43.8 | 43.8 | 43.8 | 20.546 | 2458500000 | 98 |
| ARPC4 | P59998 | ARPC4 | Actin-related protein 2/3 complex subunit 4 | 0.37628945 | -0.249162325 | -0.127127225 | 9 | 9 | 9 | 51.8 | 51.8 | 51.8 | 19.667 | 5399600000 | 143 |
| ARPC5 | O15511 | ARPC5 | Actin-related protein 2/3 complex subunit 5 | -0.22735985 | -0.051765775 | 0.279125775 | 8 | 8 | 8 | 52.3 | 52.3 | 52.3 | 16.32 | 980290000 | 63 |
| ATL1 | Q8WXF7 | ATLA1 | Atlastin-1 | -0.414620605 | 0.21598805 | 0.198632525 | 4 | 4 | 4 | 8.6 | 8.6 | 8.6 | 63.543 | 51606000 | 15 |
| ATL3 | Q6DD88 | ATLA3 | Atlastin-3 | 0.59914 | -0.33637825 | -0.262761675 | 9 | 9 | 9 | 27.4 | 27.4 | 27.4 | 60.541 | 244340000 | 27 |
| ATP2A2 | P16615 | AT2A2 | Sarcoplasmic/endoplasmic reticulum calcium ATPase 2 | 0.067066718 | 0.2487831 | -0.31584975 | 30 | 22 | 16 | 30 | 24.2 | 17.1 | 114.76 | 1569100000 | 147 |
| ATP2A3 | Q93084 | AT2A3 | Sarcoplasmic/endoplasmic reticulum calcium ATPase 3 | 0.449649175 | 0.040040075 | -0.489689225 | 33 | 33 | 24 | 33.3 | 33.3 | 26.7 | 113.98 | 6111200000 | 385 |
| ATP5F1A | P25705 | ATPA | ATP synthase subunit alpha, mitochondrial | 0.299994325 | 0.328153375 | -0.62814775 | 29 | 29 | 16 | 51.7 | 51.7 | 34 | 59.75 | 3934500000 | 332 |
| ATP5F1B | P06576 | ATPB | ATP synthase subunit beta, mitochondrial | 1.167999975 | -0.176643198 | -0.991357025 | 22 | 22 | 22 | 50.5 | 50.5 | 50.5 | 56.559 | 6448900000 | 291 |
| ATP5F1C | P36542 | ATPG | ATP synthase subunit gamma, mitochondrial | 0.70184545 | -0.497859295 | -0.20398605 | 8 | 8 | 8 | 28.5 | 28.5 | 28.5 | 32.996 | 444460000 | 37 |
| ATP5MG | E9PN17 | E9PN17 | ATP synthase subunit g, mitochondrial | 0.845874025 | -0.44089665 | -0.40497745 | 5 | 5 | 5 | 60.5 | 60.5 | 60.5 | 8.4518 | 222330000 | 27 |
| ATP5PB | Q5QNZ2 | Q5QNZ2 | ATP synthase F | 0.6026074 | -0.395350005 | -0.207257178 | 5 | 5 | 5 | 29.7 | 29.7 | 29.7 | 22.275 | 298750000 | 23 |
| ATP5PO | P48047 | ATPO | ATP synthase subunit O, mitochondrial | 0.518930675 | -0.495265125 | -0.0236655 | 6 | 6 | 6 | 35.7 | 35.7 | 35.7 | 23.277 | 518340000 | 39 |
| ATP6V1A | P38606 | VATA | V-type proton ATPase catalytic subunit A | 0.77552525 | -0.257767375 | -0.517758 | 18 | 18 | 18 | 36 | 36 | 36 | 68.303 | 446970000 | 54 |
| B2M | P61769 | B2MG | Beta-2-microglobulin | 0.56036495 | 0.090143725 | -0.65050861 | 6 | 6 | 6 | 48.7 | 48.7 | 48.7 | 13.714 | 2997300000 | 84 |
| BCAP31 | P51572 | BAP31 | B-cell receptor-associated protein 31 | 0.509592325 | -0.21850454 | -0.2910881 | 14 | 14 | 14 | 51.2 | 51.2 | 51.2 | 27.991 | 1123200000 | 86 |
| BIN2 | Q9UBW5 | BIN2 | Bridging integrator 2 | -0.011378075 | -0.532119725 | 0.54349785 | 26 | 26 | 26 | 51.9 | 51.9 | 51.9 | 61.874 | 7624100000 | 293 |
| BTK | Q06187 | BTK | Tyrosine-protein kinase BTK | 0.837315625 | -0.897881125 | 0.06056557 | 15 | 15 | 14 | 24.4 | 24.4 | 23.2 | 76.28 | 1104100000 | 48 |
| C1QC | P02747 | C1QC | Complement C1q subcomponent subunit C | -0.43712985 | -0.220464275 | 0.6575944 | 4 | 4 | 4 | 21.2 | 21.2 | 21.2 | 25.773 | 128230000 | 45 |
| C1S | A0A087X232 | A0A087X232 | Complement C1s subcomponent | -0.826034775 | -0.275835325 | 1.101870133 | 6 | 6 | 6 | 9.5 | 9.5 | 9.5 | 75.905 | 45099000 | 16 |
| C3 | P01024 | CO3 | Complement C3 | -0.516737825 | 0.21110974 | 0.305628 | 94 | 94 | 93 | 53.5 | 53.5 | 53.1 | 187.15 | 16785000000 | 1143 |
| C4A | P0C0L4 | CO4A | Complement C4-A | -1.124835225 | 0.689237 | 0.43559805 | 40 | 40 | 1 | 25.6 | 25.6 | 0.7 | 192.78 | 1967100000 | 255 |
| C4BPA | P04003 | C4BPA | C4b-binding protein alpha chain | 0.635388378 | -0.237450975 | -0.3979373 | 11 | 11 | 11 | 14.4 | 14.4 | 14.4 | 67.033 | 663830000 | 51 |
| C5 | P01031 | CO5 | Complement C5 | -1.177975075 | 0.322472583 | 0.85550239 | 12 | 12 | 12 | 7.1 | 7.1 | 7.1 | 188.3 | 70077000 | 29 |
| C8G | P07360 | CO8G | Complement component C8 gamma chain | -0.88989085 | 0.1191765 | 0.7707143 | 4 | 4 | 4 | 29.2 | 29.2 | 29.2 | 22.277 | 37237000 | 11 |
| C9 | P02748 | CO9 | Complement component C9 | -0.170990608 | -0.2786326 | 0.449623325 | 10 | 10 | 10 | 17.5 | 17.5 | 17.5 | 63.173 | 126720000 | 30 |
| CA1 | E5RHP7 | E5RHP7 | Carbonic anhydrase 1 | -0.79875804 | 0.177443575 | 0.62131435 | 11 | 11 | 11 | 55.4 | 55.4 | 55.4 | 27.754 | 476150000 | 41 |
| CA2 | P00918 | CAH2 | Carbonic anhydrase 2 | 0.01638815 | -0.34631915 | 0.329931175 | 12 | 12 | 12 | 57.3 | 57.3 | 57.3 | 29.246 | 1819300000 | 94 |
| CALD1 | E7EX44 | E7EX44 | Caldesmon | 0.698760133 | -0.621807775 | -0.07695215 | 47 | 47 | 4 | 59.2 | 59.2 | 5.2 | 64.168 | 5484700000 | 257 |
| CALM3 | P0DP25 | CALM3 | Calmodulin-3 | 0.570008795 | 0.215463353 | -0.785472275 | 11 | 11 | 9 | 60.4 | 60.4 | 55.7 | 16.837 | 3971500000 | 155 |
| CALR | P27797 | CALR | Calreticulin | 0.331127675 | 0.259127375 | -0.590255 | 18 | 18 | 18 | 47 | 47 | 47 | 48.141 | 4677500000 | 168 |
| CALU | O43852 | CALU | Calumenin | 0.2445251 | -0.163814883 | -0.08071025 | 11 | 11 | 11 | 37.8 | 37.8 | 37.8 | 37.106 | 391910000 | 28 |
| CAND1 | Q86VP6 | CAND1 | Cullin-associated NEDD8-dissociated protein 1 | 0.24601005 | -0.2648701 | 0.01885995 | 20 | 20 | 20 | 19 | 19 | 19 | 136.37 | 410180000 | 72 |
| CANX | P27824 | CALX | Calnexin | 0.712720775 | -0.69173685 | -0.020984 | 26 | 26 | 26 | 37.3 | 37.3 | 37.3 | 67.567 | 3182300000 | 173 |
| CAP1 | Q01518 | CAP1 | Adenylyl cyclase-associated protein 1 | 0.5478223 | -0.865076275 | 0.3172538 | 40 | 40 | 40 | 69.7 | 69.7 | 69.7 | 51.901 | 20610000000 | 550 |
| CAPN1 | P07384 | CAN1 | Calpain-1 catalytic subunit | 1.048590525 | -0.42205305 | -0.626537405 | 35 | 35 | 29 | 53.1 | 53.1 | 41.9 | 81.889 | 3996200000 | 276 |
| CAPNS1 | P04632 | CPNS1 | Calpain small subunit 1 | 0.8142617 | -0.537825025 | -0.276436525 | 11 | 11 | 11 | 60.4 | 60.4 | 60.4 | 28.315 | 1879300000 | 110 |
| CAPZA1 | P52907 | CAZA1 | F-actin-capping protein subunit alpha-1 | 0.95671 | -0.678475675 | -0.2782343 | 13 | 13 | 9 | 64.3 | 64.3 | 50 | 32.922 | 2564400000 | 117 |
| CAPZA2 | P47755 | CAZA2 | F-actin-capping protein subunit alpha-2 | 1.12229492 | -0.64747285 | -0.474822125 | 11 | 7 | 7 | 52.4 | 38.1 | 38.1 | 32.949 | 633370000 | 49 |
| CAPZB | B1AK87 | B1AK87 | F-actin-capping protein subunit beta | 0.718662725 | 0.155197828 | -0.8738605 | 20 | 20 | 4 | 58.8 | 58.8 | 9.2 | 29.295 | 2543400000 | 165 |
| CASS4 | Q9NQ75 | CASS4 | Cas scaffolding protein family member 4 | 0.060593675 | -0.813180922 | 0.752587325 | 18 | 18 | 18 | 27.9 | 27.9 | 27.9 | 87.143 | 176090000 | 32 |
| CAT | P04040 | CATA | Catalase | 0.051808325 | -0.11802615 | 0.066217775 | 16 | 16 | 16 | 36.8 | 36.8 | 36.8 | 59.755 | 450480000 | 66 |
| CAVIN2 | O95810 | CAVN2 | Caveolae-associated protein 2 | -0.612371375 | 0.069577825 | 0.542793458 | 30 | 30 | 30 | 55.3 | 55.3 | 55.3 | 47.173 | 25673000000 | 537 |
| CBR1 | P16152 | CBR1 | Carbonyl reductase [NADPH] 1 | -0.312293 | -0.0600514 | 0.372344575 | 8 | 8 | 7 | 35.7 | 35.7 | 30 | 30.375 | 141080000 | 34 |
| CCT2 | P78371 | TCPB | T-complex protein 1 subunit beta | 1.030001875 | -0.871824493 | -0.158177468 | 29 | 29 | 29 | 64.1 | 64.1 | 64.1 | 57.488 | 1020400000 | 103 |
| CCT3 | P49368 | TCPG | T-complex protein 1 subunit gamma | 0.838662985 | -1.0077069 | 0.1690441 | 26 | 26 | 26 | 52.8 | 52.8 | 52.8 | 60.533 | 941530000 | 105 |
| CCT4 | P50991 | TCPD | T-complex protein 1 subunit delta | 0.7980121 | -0.765968528 | -0.032043458 | 19 | 19 | 19 | 40.3 | 40.3 | 40.3 | 57.924 | 562130000 | 79 |
| CCT5 | P48643 | TCPE | T-complex protein 1 subunit epsilon | 0.72509765 | -0.057784575 | -0.6673128 | 17 | 17 | 17 | 34.4 | 34.4 | 34.4 | 59.67 | 705400000 | 66 |
| CCT6A | P40227 | TCPZ | T-complex protein 1 subunit zeta | 0.57575335 | -0.05606456 | -0.51968887 | 14 | 14 | 11 | 28.2 | 28.2 | 23.2 | 58.024 | 762080000 | 100 |
| CCT7 | Q99832 | TCPH | T-complex protein 1 subunit eta | 0.706802175 | -0.5817485 | -0.125053775 | 19 | 19 | 19 | 36.8 | 36.8 | 36.8 | 59.366 | 662060000 | 88 |
| CCT8 | P50990 | TCPQ | T-complex protein 1 subunit theta | 0.71304072 | 0.125785243 | -0.838826175 | 21 | 21 | 21 | 44.2 | 44.2 | 44.2 | 59.62 | 1014800000 | 124 |
| CD36 | E7EU05 | E7EU05 | Platelet glycoprotein 4 | 0.501746873 | 0.140372213 | -0.64211885 | 14 | 14 | 14 | 24.2 | 24.2 | 24.2 | 52.062 | 3240900000 | 185 |
| CD47 | Q08722 | CD47 | Leukocyte surface antigen CD47 | 0.533653525 | -0.196899048 | -0.336754363 | 4 | 4 | 4 | 10.8 | 10.8 | 10.8 | 35.213 | 1022100000 | 48 |
| CD84 | Q9UIB8 | SLAF5 | SLAM family member 5 | 0.531626535 | -0.499158825 | -0.032467825 | 9 | 9 | 9 | 35.9 | 35.9 | 35.9 | 38.782 | 262430000 | 30 |
| CD9 | A6NNI4 | A6NNI4 | Tetraspanin | 0.396485615 | 0.511854775 | -0.908340375 | 7 | 7 | 7 | 20.8 | 20.8 | 20.8 | 17.764 | 4342600000 | 122 |
| CDC42 | P60953 | CDC42 | Cell division control protein 42 homolog | -0.17593395 | -0.02032185 | 0.196255675 | 7 | 7 | 7 | 36.6 | 36.6 | 36.6 | 21.258 | 1236100000 | 72 |
| CFB | P00751 | CFAB | Complement factor B | -0.165068025 | -0.2591156 | 0.424183775 | 14 | 14 | 14 | 18.8 | 18.8 | 18.8 | 85.532 | 169870000 | 39 |
| CFL1 | P23528 | COF1 | Cofilin-1 | -0.942733775 | 0.158658175 | 0.784075675 | 20 | 20 | 19 | 88.6 | 88.6 | 84.3 | 18.502 | 19883000000 | 333 |
| CKAP5 | Q14008 | CKAP5 | Cytoskeleton-associated protein 5 | -0.034016025 | -0.081600055 | 0.115616125 | 25 | 25 | 25 | 14.8 | 14.8 | 14.8 | 225.49 | 237100000 | 64 |
| CLIC1 | O00299 | CLIC1 | Chloride intracellular channel protein 1 | 0.391727375 | 0.472695475 | -0.86442275 | 17 | 17 | 17 | 81.3 | 81.3 | 81.3 | 26.922 | 6517800000 | 193 |
| CLINT1 | Q14677 | EPN4 | Clathrin interactor 1 | 0.750537089 | -0.832907925 | 0.082371025 | 12 | 12 | 12 | 21.8 | 21.8 | 21.8 | 68.259 | 342330000 | 33 |
| CLTA | P09496 | CLCA | Clathrin light chain A | 0.206117425 | -0.442703375 | 0.236585945 | 6 | 6 | 6 | 16.9 | 16.9 | 16.9 | 27.076 | 380940000 | 38 |
| CLTC | Q00610 | CLH1 | Clathrin heavy chain 1 | 0.99412135 | -0.11880867 | -0.875312775 | 65 | 65 | 65 | 44.2 | 44.2 | 44.2 | 191.61 | 4883400000 | 340 |
| CLU | P10909 | CLUS | Clusterin | -0.80307205 | 0.476846188 | 0.326225925 | 18 | 18 | 18 | 37 | 37 | 37 | 52.494 | 3529000000 | 202 |
| CMPK1 | P30085 | KCY | UMP-CMP kinase | 0.41555375 | 0.15264115 | -0.5681948 | 8 | 8 | 8 | 50.5 | 50.5 | 50.5 | 22.222 | 193960000 | 31 |
| CMTM5 | C9JAI6 | C9JAI6 | CKLF-like MARVEL transmembrane domain-containing protein 5 | 1.05197355 | -0.665463125 | -0.3865105 | 3 | 3 | 3 | 33.1 | 33.1 | 33.1 | 13.238 | 730350000 | 27 |
| CMTM6 | Q9NX76 | CKLF6 | CKLF-like MARVEL transmembrane domain-containing protein 6 | -0.61628275 | 0.867660075 | -0.25137735 | 5 | 5 | 5 | 29.5 | 29.5 | 29.5 | 20.419 | 309440000 | 27 |
| CNDP2 | Q96KP4 | CNDP2 | Cytosolic non-specific dipeptidase | 0.69673725 | -0.337227825 | -0.359509425 | 15 | 15 | 15 | 42.3 | 42.3 | 42.3 | 52.878 | 286450000 | 50 |
| CNN2 | B4DDF4 | B4DDF4 | Calponin | 0.74150703 | -0.761214875 | 0.019707775 | 20 | 20 | 17 | 71.8 | 71.8 | 69.5 | 32.616 | 3629000000 | 190 |
| CNST | Q6PJW8 | CNST | Consortin | 0.364189325 | -0.12804025 | -0.236149025 | 13 | 13 | 13 | 20.8 | 20.8 | 20.8 | 79.596 | 283150000 | 36 |
| COPA | P53621 | COPA | Coatomer subunit alpha | 0.134756 | 0.478836 | -0.613591963 | 25 | 25 | 25 | 23.4 | 23.4 | 23.4 | 138.34 | 351640000 | 56 |
| COPB2 | P35606 | COPB2 | Coatomer subunit beta' | 0.364947975 | -0.440832425 | 0.075884438 | 17 | 17 | 17 | 23.3 | 23.3 | 23.3 | 102.49 | 251320000 | 38 |
| CORO1A | P31146 | COR1A | Coronin-1A | 0.804659925 | -0.6709777 | -0.133682118 | 21 | 20 | 20 | 40.1 | 38.6 | 38.6 | 51.026 | 6931500000 | 167 |
| CORO1B | Q9BR76 | COR1B | Coronin-1B | 0.7586096 | -0.084216582 | -0.6743928 | 10 | 9 | 8 | 19.6 | 17.8 | 16.4 | 54.234 | 434410000 | 45 |
| CORO1C | Q9ULV4 | COR1C | Coronin-1C | 0.313078275 | -0.668874268 | 0.355796025 | 27 | 27 | 24 | 43.2 | 43.2 | 37.6 | 53.248 | 7612300000 | 262 |
| COTL1 | Q14019 | COTL1 | Coactosin-like protein | 0.30167925 | 0.20535905 | -0.507038375 | 15 | 15 | 15 | 86.6 | 86.6 | 86.6 | 15.945 | 3133300000 | 111 |
| COX4I1 | P13073 | COX41 | Cytochrome c oxidase subunit 4 isoform 1, mitochondrial | -0.039570818 | 0.831137525 | -0.7915668 | 8 | 8 | 8 | 41.4 | 41.4 | 41.4 | 19.576 | 728640000 | 57 |
| COX5A | P20674 | COX5A | Cytochrome c oxidase subunit 5A, mitochondrial | 0.42426335 | -0.245449253 | -0.178813878 | 7 | 7 | 7 | 38 | 38 | 38 | 16.762 | 481370000 | 49 |
| CP | E9PFZ2 | E9PFZ2 | Ceruloplasmin | 0.3836078 | -0.312508825 | -0.071099125 | 7 | 7 | 7 | 8.9 | 8.9 | 8.9 | 108.82 | 49376000 | 14 |
| CPT1A | P50416 | CPT1A | Carnitine O-palmitoyltransferase 1, liver isoform | -0.112957925 | -0.26371505 | 0.376672675 | 10 | 10 | 10 | 14.9 | 14.9 | 14.9 | 88.367 | 201060000 | 27 |
| CRKL | P46109 | CRKL | Crk-like protein | -0.1028433 | 0.1980647 | -0.09522145 | 11 | 11 | 11 | 47.2 | 47.2 | 47.2 | 33.777 | 409250000 | 41 |
| CS | B4DJV2 | B4DJV2 | Citrate synthase | 0.97847635 | -0.1852571 | -0.793219225 | 13 | 13 | 12 | 26.9 | 26.9 | 24.9 | 50.431 | 986350000 | 77 |
| CSK | P41240 | CSK | Tyrosine-protein kinase CSK | 0.29247812 | -0.54956905 | 0.257090925 | 15 | 15 | 15 | 38 | 38 | 38 | 50.704 | 440480000 | 49 |
| CSRP1 | P21291 | CSRP1 | Cysteine and glycine-rich protein 1 | 0.816093125 | -0.636413575 | -0.179679325 | 5 | 5 | 5 | 36.8 | 36.8 | 36.8 | 20.567 | 5143700000 | 69 |
| CTSA | P10619 | PPGB | Lysosomal protective protein | 0.157083725 | -0.160081825 | 0.002998025 | 5 | 5 | 5 | 11.9 | 11.9 | 11.9 | 54.465 | 173370000 | 24 |
| CTSD | A0A1B0GW44 | A0A1B0GW44 | Cathepsin D | 0.4187061 | -0.528676225 | 0.109970085 | 7 | 7 | 7 | 21.2 | 21.2 | 21.2 | 43.688 | 131430000 | 19 |
| CTTN | Q14247 | SRC8 | Src substrate cortactin | 0.595246825 | 0.0166468 | -0.611893775 | 34 | 34 | 34 | 54.9 | 54.9 | 54.9 | 61.585 | 5406000000 | 228 |
| CYB5R1 | Q9UHQ9 | NB5R1 | NADH-cytochrome b5 reductase 1 | -0.203002475 | 0.207101513 | -0.004098875 | 7 | 7 | 7 | 26.9 | 26.9 | 26.9 | 34.094 | 126180000 | 23 |
| CYB5R3 | P00387 | NB5R3 | NADH-cytochrome b5 reductase 3 | 0.4708411 | -0.580605933 | 0.109764885 | 18 | 18 | 18 | 60.1 | 60.1 | 60.1 | 34.234 | 2195900000 | 139 |
| CYCS | C9JFR7 | C9JFR7 | Cytochrome c | -0.133903 | 3.215E-05 | 0.133870975 | 5 | 5 | 5 | 57.4 | 57.4 | 57.4 | 11.333 | 185090000 | 22 |
| CYFIP1 | Q7L576 | CYFP1 | Cytoplasmic FMR1-interacting protein 1 | 0.766622286 | -0.539177388 | -0.2274449 | 32 | 32 | 18 | 27.7 | 27.7 | 16.4 | 145.18 | 1031400000 | 125 |
| DAD1 | F5GXX5 | F5GXX5 | Dolichyl-diphosphooligosaccharide--protein glycosyltransferase subunit DAD1 | 0.100293325 | 0.15044025 | -0.250733625 | 3 | 3 | 3 | 37.6 | 37.6 | 37.6 | 9.5541 | 218140000 | 31 |
| DBN1 | Q16643 | DREB | Drebrin | 0.825316675 | -0.71514117 | -0.110175275 | 18 | 18 | 18 | 31 | 31 | 31 | 71.428 | 706620000 | 70 |
| DBNL | Q9UJU6 | DBNL | Drebrin-like protein | 0.680538225 | -0.1940378 | -0.48650065 | 21 | 21 | 5 | 52.1 | 52.1 | 12.1 | 48.207 | 2504700000 | 148 |
| DCD | P81605 | DCD | Dermcidin | -0.542082975 | 0.41872505 | 0.1233579 | 6 | 6 | 6 | 36.4 | 36.4 | 36.4 | 11.284 | 240390000 | 22 |
| DDOST | A0A0C4DGS1 | A0A0C4DGS1 | Dolichyl-diphosphooligosaccharide--protein glycosyltransferase 48 kDa subunit | 0.92115045 | -0.7943827 | -0.126767525 | 13 | 13 | 13 | 29.4 | 29.4 | 29.4 | 48.799 | 428400000 | 45 |
| DECR1 | Q16698 | DECR | 2,4-dienoyl-CoA reductase, mitochondrial | 0.685763775 | -0.347302625 | -0.338461175 | 7 | 7 | 7 | 28.7 | 28.7 | 28.7 | 36.067 | 300400000 | 32 |
| DIAPH1 | A0A140T8Z0 | A0A140T8Z0 | Protein diaphanous homolog 1 | 0.81315995 | -0.485452975 | -0.327707106 | 37 | 37 | 37 | 30.3 | 30.3 | 30.3 | 140.3 | 1288600000 | 137 |
| DLST | P36957 | ODO2 | Dihydrolipoyllysine-residue succinyltransferase component of 2-oxoglutarate dehydrogenase complex, mitochondrial | -0.422120415 | 0.6882418 | -0.266121125 | 6 | 6 | 6 | 14.8 | 14.8 | 14.8 | 48.755 | 82609000 | 23 |
| DMTN | Q08495 | DEMA | Dematin | 0.402064448 | -0.323288025 | -0.078776225 | 15 | 15 | 15 | 44.9 | 44.9 | 44.9 | 45.514 | 1261500000 | 79 |
| DNM1 | A0A0U1RQP1 | A0A0U1RQP1 | Dynamin-1 | -0.192976599 | 0.5224418 | -0.329465375 | 13 | 13 | 8 | 18.2 | 18.2 | 13 | 96.396 | 297500000 | 49 |
| DNM1L | O00429 | DNM1L | Dynamin-1-like protein | 0.78498335 | -0.877539575 | 0.092556125 | 27 | 27 | 27 | 43.8 | 43.8 | 43.8 | 81.876 | 2042500000 | 157 |
| DOK2 | O60496 | DOK2 | Docking protein 2 | 0.340195425 | -0.366744275 | 0.02654915 | 5 | 5 | 5 | 17.7 | 17.7 | 17.7 | 45.378 | 270650000 | 25 |
| DUSP3 | P51452 | DUS3 | Dual specificity protein phosphatase 3 | 0.27369935 | 0.3024098 | -0.576109305 | 8 | 8 | 8 | 55.1 | 55.1 | 55.1 | 20.478 | 589610000 | 53 |
| ECH1 | Q13011 | ECH1 | Delta | 0.3499027 | 0.439753062 | -0.78965597 | 10 | 10 | 10 | 39.3 | 39.3 | 39.3 | 35.816 | 329270000 | 36 |
| EEF1A1P5 | Q5VTE0 | EF1A3 | Putative elongation factor 1-alpha-like 3 | 1.131717975 | -0.469615975 | -0.662102268 | 12 | 12 | 12 | 31.4 | 31.4 | 31.4 | 50.184 | 837020000 | 74 |
| EEF1B2 | P24534 | EF1B | Elongation factor 1-beta | 0.62887805 | -0.536874375 | -0.09200355 | 7 | 6 | 6 | 38.7 | 35.6 | 35.6 | 24.763 | 326250000 | 29 |
| EEF1D | E9PK01 | E9PK01 | Elongation factor 1-delta | 1.1153267 | -0.45259315 | -0.66273365 | 9 | 9 | 8 | 36.4 | 36.4 | 33.7 | 28.821 | 256870000 | 24 |
| EEF1G | P26641 | EF1G | Elongation factor 1-gamma | 0.7797416 | -0.227283725 | -0.55245765 | 17 | 17 | 17 | 38.7 | 38.7 | 38.7 | 50.118 | 1216800000 | 107 |
| EEF2 | P13639 | EF2 | Elongation factor 2 | -0.359266575 | -0.2274325 | 0.5866992 | 2 | 2 | 2 | 2.3 | 2.3 | 2.3 | 95.337 | 33707000 | 7 |
| EHD1 | Q9H4M9 | EHD1 | EH domain-containing protein 1 | 0.77360515 | -0.365555425 | -0.408049675 | 35 | 19 | 19 | 60.3 | 35.6 | 35.6 | 60.626 | 765500000 | 81 |
| EHD3 | Q9NZN3 | EHD3 | EH domain-containing protein 3 | 0.8105492 | -0.051999248 | -0.75854975 | 36 | 36 | 18 | 68.8 | 68.8 | 39.1 | 60.886 | 4145200000 | 270 |
| EIF2S2 | P20042 | IF2B | Eukaryotic translation initiation factor 2 subunit 2 | -0.510189125 | 0.153263275 | 0.35692583 | 3 | 3 | 3 | 9.9 | 9.9 | 9.9 | 38.388 | 16764000 | 7 |
| EIF4A1 | P60842 | IF4A1 | Eukaryotic initiation factor 4A-I | 0.932395525 | -0.256510745 | -0.675884875 | 17 | 17 | 2 | 43.6 | 43.6 | 6.2 | 46.153 | 840780000 | 71 |
| EMD | P50402 | EMD | Emerin | 0.940118225 | -0.588977675 | -0.3511402 | 11 | 11 | 11 | 58.7 | 58.7 | 58.7 | 28.994 | 526150000 | 48 |
| EMILIN1 | Q9Y6C2 | EMIL1 | EMILIN-1 | 1.03847845 | -0.388773175 | -0.649705425 | 21 | 21 | 20 | 25.6 | 25.6 | 25 | 106.69 | 1335000000 | 101 |
| ENDOD1 | O94919 | ENDD1 | Endonuclease domain-containing 1 protein | 0.535152925 | -0.181284925 | -0.353867708 | 14 | 14 | 14 | 31.6 | 31.6 | 31.6 | 55.016 | 828090000 | 77 |
| ENO1 | P06733 | ENOA | Alpha-enolase | 0.08960215 | -0.177527325 | 0.087925208 | 37 | 37 | 34 | 74.9 | 74.9 | 73.5 | 47.168 | 12515000000 | 511 |
| EPS15 | B1AUU8 | B1AUU8 | Epidermal growth factor receptor substrate 15 | -0.488061125 | 0.550090575 | -0.0620295 | 7 | 7 | 7 | 10.2 | 10.2 | 10.2 | 83.653 | 96674000 | 17 |
| EPS15L1 | M0R165 | M0R165 | Epidermal growth factor receptor substrate 15-like 1 | -0.536872025 | -0.124047025 | 0.66091882 | 11 | 11 | 11 | 18.4 | 18.4 | 18.4 | 83.474 | 95897000 | 27 |
| ERAP1 | Q9NZ08 | ERAP1 | Endoplasmic reticulum aminopeptidase 1 | 0.0388512 | 0.260943533 | -0.299794975 | 16 | 16 | 16 | 19.6 | 19.6 | 19.6 | 107.23 | 271140000 | 37 |
| ERP29 | P30040 | ERP29 | Endoplasmic reticulum resident protein 29 | 0.177872325 | 0.113631002 | -0.2915034 | 9 | 9 | 9 | 36.8 | 36.8 | 36.8 | 28.993 | 321300000 | 36 |
| ERP44 | Q9BS26 | ERP44 | Endoplasmic reticulum resident protein 44 | 0.6733579 | -0.269692625 | -0.4036651 | 8 | 8 | 8 | 19 | 19 | 19 | 46.971 | 167640000 | 29 |
| ESAM | Q96AP7 | ESAM | Endothelial cell-selective adhesion molecule | 0.217695125 | -0.419640775 | 0.201945545 | 5 | 5 | 5 | 22.1 | 22.1 | 22.1 | 41.176 | 193440000 | 26 |

| | | | | | | | | | | | | | | | |
|---|---|---|---|---|---|---|---|---|---|---|---|---|---|---|---|
| ESYT1 | Q9BSJ8 | ESYT1 | Extended synaptotagmin-1 | 0.728515859 | -0.1268696 | -0.60164623 | 28 | 28 | 28 | 29.6 | 29.6 | 29.6 | 122.85 | 623780000 | 86 |
| ESYT2 | H7BXI1 | H7BXI1 | Extended synaptotagmin-2 | 0.3308166 | -0.55456495 | 0.223748475 | 10 | 10 | 10 | 14.6 | 14.6 | 14.6 | 98.008 | 177260000 | 30 |
| ETFB | P38117 | ETFB | Electron transfer flavoprotein subunit beta | 0.426405275 | -0.269118825 | -0.157286425 | 5 | 5 | 5 | 23.1 | 23.1 | 23.1 | 27.843 | 92290000 | 19 |
| F11R | Q9Y624 | JAM1 | Junctional adhesion molecule A | 0.347470195 | -0.075743418 | -0.271727005 | 13 | 13 | 13 | 43.1 | 43.1 | 43.1 | 32.583 | 1907500000 | 90 |
| F13A1 | P00488 | F13A | Coagulation factor XIII A chain | 0.641987575 | 0.084847568 | -0.72683505 | 48 | 48 | 48 | 53.7 | 53.7 | 53.7 | 83.266 | 46618000000 | 776 |
| F2 | P00734 | THRB | Prothrombin | -0.71338911 | 0.302017888 | 0.41137115 | 7 | 7 | 7 | 10.8 | 10.8 | 10.8 | 70.036 | 233020000 | 48 |
| F5 | P12259 | FA5 | Coagulation factor V | 1.048784325 | -0.245233948 | -0.803550525 | 49 | 49 | 39 | 24.5 | 24.5 | 20 | 251.7 | 2339200000 | 257 |
| FAH | P16930 | FAAA | Fumarylacetoacetase | 0.128267275 | 0.14988965 | -0.278156875 | 7 | 7 | 7 | 20.8 | 20.8 | 20.8 | 46.374 | 249580000 | 23 |
| FASN | A0A0U1RQF0 | A0A0U1RQF0 | Fatty acid synthase | 0.2204731 | 0.31265861 | -0.533131878 | 25 | 25 | 25 | 12.9 | 12.9 | 12.9 | 273.2 | 322040000 | 58 |
| FCER1G | P30273 | FCERG | High affinity immunoglobulin epsilon receptor subunit gamma | 0.404604693 | 0.1317677 | -0.536372828 | 6 | 6 | 6 | 40.7 | 40.7 | 40.7 | 9.6674 | 708890000 | 36 |
| FERMT3 | Q86UX7 | URP2 | Fermitin family homolog 3 | 0.8350965 | -0.427945 | -0.407151825 | 52 | 52 | 52 | 73.3 | 73.3 | 73.3 | 75.952 | 60007000000 | 1051 |
| FGA | P02671 | FIBA | Fibrinogen alpha chain | -0.639254065 | -0.420576308 | 1.0598306 | 62 | 62 | 62 | 60 | 60 | 60 | 94.972 | 1.1667E+11 | 1617 |
| FGB | P02675 | FIBB | Fibrinogen beta chain | 0.315505975 | -0.61015425 | 0.2946481 | 45 | 45 | 45 | 70.3 | 70.3 | 70.3 | 55.928 | 54729000000 | 989 |
| FGG | P02679 | FIBG | Fibrinogen gamma chain | 0.25557936 | 0.073484725 | -0.32906405 | 31 | 31 | 20 | 66.9 | 66.9 | 48.6 | 51.511 | 82937000000 | 986 |
| FH | P07954 | FUMH | Fumarate hydratase, mitochondrial | 0.471844025 | 0.025737225 | -0.497581273 | 8 | 8 | 8 | 25.3 | 25.3 | 25.3 | 54.636 | 192480000 | 26 |
| FHL1 | Q5JXI8 | Q5JXI8 | Four and a half LIM domains protein 1 | 0.14670435 | 0.035204913 | -0.18190899 | 9 | 9 | 9 | 33.5 | 33.5 | 33.5 | 29.16 | 1682000000 | 85 |
| FHOD1 | Q9Y613 | FHOD1 | FH1/FH2 domain-containing protein 1 | 0.4355858 | -0.04063795 | -0.394947775 | 32 | 32 | 32 | 33.7 | 33.7 | 33.7 | 126.55 | 937870000 | 116 |
| FKBP1A | P62942 | FKB1A | Peptidyl-prolyl cis-trans isomerase FKBP1A | 0.779318975 | -0.690401725 | -0.088916875 | 2 | 2 | 2 | 25 | 25 | 25 | 11.951 | 631790000 | 38 |
| FLII | Q13045 | FLII | Protein flightless-1 homolog | 0.236561875 | -0.481900075 | 0.2453383 | 21 | 21 | 18 | 18.3 | 18.3 | 15.3 | 144.75 | 387180000 | 71 |
| FLNA | P21333 | FLNA | Filamin-A | 0.7785388 | -0.195638475 | -0.582900288 | 191 | 191 | 0 | 74 | 74 | 0 | 280.74 | 3.5948E+11 | 4345 |
| FN1 | P02751 | FINC | Fibronectin | -0.696052 | 0.144111675 | 0.551940325 | 47 | 47 | 47 | 25.4 | 25.4 | 25.4 | 262.62 | 4138000000 | 422 |
| FYB1 | O15117 | FYB1 | FYN-binding protein 1 | 0.666421325 | -0.096045575 | -0.570375403 | 18 | 18 | 18 | 25.3 | 25.3 | 25.3 | 85.386 | 1074400000 | 113 |
| G6PD | P11413 | G6PD | Glucose-6-phosphate 1-dehydrogenase | 0.919860325 | -0.64631985 | -0.2735406 | 24 | 24 | 7 | 47.6 | 47.6 | 14.4 | 59.256 | 1687800000 | 120 |
| GANAB | Q14697 | GANAB | Neutral alpha-glucosidase AB | 1.051759475 | -0.5139114 | -0.537848085 | 37 | 37 | 37 | 41.4 | 41.4 | 41.4 | 106.87 | 1806300000 | 145 |
| GAPDH | P04406 | G3P | Glyceraldehyde-3-phosphate dehydrogenase | 0.53848465 | -0.47516275 | -0.0633218 | 27 | 27 | 27 | 83.9 | 83.9 | 83.9 | 36.053 | 34463000000 | 713 |
| GC | P02774 | VTDB | Vitamin D-binding protein | -0.87923075 | 0.5002592 | 0.37897177 | 10 | 10 | 1 | 18.1 | 18.1 | 1.9 | 52.963 | 176800000 | 21 |
| GDI1 | P31150 | GDIA | Rab GDP dissociation inhibitor alpha | 0.41814195 | -0.07901095 | -0.33913129 | 19 | 12 | 12 | 45 | 32.2 | 32.2 | 50.582 | 560410000 | 73 |
| GDI2 | P50395 | GDIB | Rab GDP dissociation inhibitor beta | 0.258454693 | 0.474458525 | -0.73291317 | 22 | 22 | 15 | 51.9 | 51.9 | 39.1 | 50.663 | 2924500000 | 120 |
| GMFG | O60234 | GMFG | Glia maturation factor gamma | -0.3571133 | 0.28952182 | 0.067591418 | 5 | 5 | 4 | 40.1 | 40.1 | 33.8 | 16.801 | 626650000 | 27 |
| GNA13 | Q14344 | GNA13 | Guanine nucleotide-binding protein subunit alpha-13 | 0.460166025 | -1.013476125 | 0.553310023 | 12 | 11 | 11 | 33.7 | 31.6 | 31.6 | 44.049 | 259280000 | 44 |
| GNAI2 | P04899 | GNAI2 | Guanine nucleotide-binding protein G | 0.17216365 | -0.515440925 | 0.3432773 | 15 | 15 | 8 | 48.7 | 48.7 | 29.9 | 40.45 | 1889000000 | 164 |
| GNAQ | P50148 | GNAQ | Guanine nucleotide-binding protein G | 1.0174778 | -0.53500285 | -0.482475115 | 15 | 15 | 14 | 46.5 | 46.5 | 44.3 | 42.142 | 2146200000 | 119 |
| GNAZ | P19086 | GNAZ | Guanine nucleotide-binding protein G (z) subunit alpha | 0.8053704 | -0.4680227 | -0.337347625 | 12 | 11 | 11 | 43.7 | 41.4 | 41.4 | 40.923 | 534060000 | 58 |
| GNB1 | P62873 | GBB1 | Guanine nucleotide-binding protein G | 0.662376625 | 0.212611103 | -0.87498774 | 13 | 13 | 7 | 41.2 | 41.2 | 20 | 37.377 | 3029500000 | 99 |
| GNG5 | P63218 | GBG5 | Guanine nucleotide-binding protein G | -0.3255687 | 0.489656603 | -0.164088 | 5 | 5 | 5 | 48.5 | 48.5 | 48.5 | 7.3184 | 377030000 | 28 |
| GOT2 | P00505 | AATM | Aspartate aminotransferase, mitochondrial | 0.213366165 | 0.109689538 | -0.323055668 | 8 | 8 | 8 | 21.6 | 21.6 | 21.6 | 47.517 | 241910000 | 39 |
| GP1BA | A0A0C4DGZ8 | A0A0C4DGZ8 | Glycoprotein Ib | 0.222756465 | -0.07133315 | -0.151423275 | 14 | 14 | 14 | 23.2 | 23.2 | 23.2 | 68.954 | 8736100000 | 297 |
| GP1BB | P13224 | GP1BB | Platelet glycoprotein Ib beta chain | 0.7086062 | -0.48892225 | -0.219684075 | 6 | 6 | 6 | 24.8 | 24.8 | 24.8 | 21.717 | 5687500000 | 110 |
| GP5 | P40197 | GPV | Platelet glycoprotein V | 0.23771705 | -0.3274679 | 0.089750825 | 17 | 17 | 17 | 34.5 | 34.5 | 34.5 | 60.958 | 2534400000 | 161 |
| GP6 | Q9HCN6 | GPVI | Platelet glycoprotein VI | 0.46342305 | -0.21826919 | -0.245153675 | 7 | 7 | 7 | 24.5 | 24.5 | 24.5 | 36.866 | 469460000 | 40 |
| GP9 | P14770 | GPIX | Platelet glycoprotein IX | 0.61581565 | 0.003109625 | -0.6189253 | 6 | 6 | 6 | 30.5 | 30.5 | 30.5 | 19.046 | 2742400000 | 106 |
| GPD2 | P43304 | GPDM | Glycerol-3-phosphate dehydrogenase, mitochondrial | 0.851257875 | -0.036918923 | -0.81433886 | 27 | 27 | 27 | 38.9 | 38.9 | 38.9 | 80.852 | 979490000 | 101 |
| GPI | P06744 | G6PI | Glucose-6-phosphate isomerase | 0.936016675 | -0.65370712 | -0.282309475 | 22 | 22 | 22 | 42.3 | 42.3 | 42.3 | 63.146 | 1840100000 | 125 |
| GPX1 | A0A087WUQ6 | A0A087WUQ6 | Glutathione peroxidase | 0.7071804 | -0.55744185 | -0.149738825 | 13 | 13 | 13 | 80.7 | 80.7 | 80.7 | 21.938 | 1223500000 | 87 |
| GRB2 | P62993 | GRB2 | Growth factor receptor-bound protein 2 | -0.48352875 | 0.150486 | 0.333042965 | 12 | 12 | 12 | 50.7 | 50.7 | 50.7 | 25.206 | 654910000 | 55 |
| GSN | P06396 | GELS | Gelsolin | 0.225664025 | -0.259077825 | 0.03341375 | 50 | 50 | 3 | 56.3 | 56.3 | 3.7 | 85.696 | 43115000000 | 884 |
| GSN | Q5T0I0 | Q5T0I0 | Gelsolin | 1.0372717 | -0.5117858 | -0.525486025 | 13 | 1 | 1 | 41.9 | 3.5 | 3.5 | 28.953 | 2019700000 | 47 |
| GSR | P00390 | GSHR | Glutathione reductase, mitochondrial | 0.0430426 | -0.2538247 | 0.2107822 | 9 | 9 | 9 | 22.4 | 22.4 | 22.4 | 56.256 | 408660000 | 26 |
| GSTK1 | Q9Y2Q3 | GSTK1 | Glutathione S-transferase kappa 1 | 0.435376825 | -0.711538925 | 0.276162175 | 9 | 9 | 9 | 54.9 | 54.9 | 54.9 | 25.497 | 147120000 | 32 |
| GSTO1 | P78417 | GSTO1 | Glutathione S-transferase omega-1 | 0.942041 | -0.4786224 | -0.463418523 | 18 | 18 | 18 | 52.3 | 52.3 | 52.3 | 27.566 | 3202100000 | 109 |
| GSTP1 | P09211 | GSTP1 | Glutathione S-transferase P | 0.435086525 | -0.58544509 | 0.150358475 | 12 | 12 | 12 | 69.5 | 69.5 | 69.5 | 23.356 | 2309800000 | 107 |
| GTPBP2 | Q9BX10 | GTPB2 | GTP-binding protein 2 | 0.46472475 | -0.474371275 | 0.009646675 | 11 | 11 | 11 | 19.9 | 19.9 | 19.9 | 65.768 | 351480000 | 39 |
| GYG1 | G5E9W8 | G5E9W8 | Glycogenin 1, isoform CRA_e | -0.7136446 | 0.42151175 | 0.292132863 | 4 | 4 | 4 | 25.9 | 25.9 | 25.9 | 21.51 | 175130000 | 15 |
| HADHA | P40939 | ECHA | Trifunctional enzyme subunit alpha, mitochondrial | 0.1591673 | -0.29086925 | 0.131702 | 22 | 22 | 22 | 36 | 36 | 36 | 82.999 | 655460000 | 83 |
| HADHB | P55084 | ECHB | Trifunctional enzyme subunit beta, mitochondrial | 0.943359475 | -0.440328175 | -0.5030315 | 11 | 11 | 11 | 26.4 | 26.4 | 26.4 | 51.294 | 484380000 | 52 |
| HBA1; | P69905 | HBA | Hemoglobin subunit alpha | -0.284160825 | -0.7961623 | 1.080323025 | 10 | 10 | 3 | 71.8 | 71.8 | 18.3 | 15.257 | 42283000000 | 385 |
| HBB | P68871 | HBB | Hemoglobin subunit beta | -0.36558735 | -0.56900645 | 0.934593648 | 13 | 13 | 6 | 84.4 | 84.4 | 51 | 15.998 | 69848000000 | 510 |
| HIST1H2BM | Q99879 | H2B1M | Histone H2B type 1-M | 0.6529655 | -0.561715603 | -0.0912497 | 8 | 8 | 8 | 46.8 | 46.8 | 46.8 | 13.989 | 487440000 | 33 |
| HIST1H4A | P62805 | H4 | Histone H4 | 0.934810623 | -0.341152575 | -0.5936581 | 5 | 5 | 5 | 50.5 | 50.5 | 50.5 | 11.367 | 333650000 | 31 |
| HK1 | P19367 | HXK1 | Hexokinase-1 | 0.69684345 | 0.35244635 | -1.049289975 | 42 | 42 | 42 | 43.3 | 43.3 | 43.3 | 102.48 | 3034100000 | 265 |
| HLA-A | P10316 | 1A69 | HLA class I histocompatibility antigen, A-69 alpha chain | 0.083071975 | -0.779755933 | 0.69668404 | 20 | 20 | 0 | 52.6 | 52.6 | 0 | 40.976 | 1754700000 | 106 |
| HLA-A | P13746 | 1A11 | HLA class I histocompatibility antigen, A-11 alpha chain | 0.502503975 | -0.007345908 | -0.495158075 | 19 | 7 | 2 | 51.8 | 24.1 | 3.3 | 40.936 | 505910000 | 36 |
| HLA-B | P01889 | 1B07 | HLA class I histocompatibility antigen, B-7 alpha chain | 0.146880685 | -0.353861228 | 0.2069806 | 18 | 10 | 2 | 47.8 | 32.3 | 5.5 | 40.46 | 1597200000 | 85 |
| HLA-C | O19617 | O19617 | HLA class I antigen | 0.628966125 | -0.016015018 | -0.61295095 | 12 | 1 | 1 | 36.6 | 4.4 | 4.4 | 40.71 | 995520000 | 12 |
| HP | P00738 | HPT | Haptoglobin | -0.786831075 | 0.08608135 | 0.700749635 | 20 | 20 | 11 | 38.9 | 38.9 | 23.2 | 45.205 | 4858700000 | 215 |
| HPRT1 | P00492 | HPRT | Hypoxanthine-guanine phosphoribosyltransferase | 0.117297725 | -0.390600975 | 0.2733031 | 8 | 8 | 7 | 41.3 | 41.3 | 38.1 | 24.579 | 216600000 | 31 |
| HPSE | Q9Y251 | HPSE | Heparanase | 0.529736068 | -0.1456325 | -0.3841037 | 15 | 15 | 15 | 27.1 | 27.1 | 27.1 | 61.148 | 1126700000 | 98 |
| HPX | P02790 | HEMO | Hemopexin | 0.120195525 | 0.06868495 | -0.188880425 | 10 | 10 | 10 | 22.7 | 22.7 | 22.7 | 51.676 | 263760000 | 52 |
| HRG | P04196 | HRG | Histidine-rich glycoprotein | 0.313655325 | -0.3131516 | -0.0005038 | 11 | 11 | 11 | 22.3 | 22.3 | 22.3 | 59.578 | 443240000 | 32 |
| HSD17B12 | Q53GQ0 | DHB12 | Very-long-chain 3-oxoacyl-CoA reductase | 0.39323565 | -0.185917215 | -0.207318323 | 9 | 9 | 9 | 35.9 | 35.9 | 35.9 | 34.324 | 301570000 | 33 |
| HSD17B4 | P51659 | DHB4 | Peroxisomal multifunctional enzyme type 2 | 0.819952775 | 0.04503207 | -0.864984825 | 29 | 29 | 29 | 47.8 | 47.8 | 47.8 | 79.685 | 1470400000 | 157 |
| HSP90AA1 | P07900 | HS90A | Heat shock protein HSP 90-alpha | 0.4970407 | 0.064355095 | -0.561395825 | 32 | 32 | 21 | 39.5 | 39.5 | 24.3 | 84.659 | 5798800000 | 296 |
| HSP90AB1 | P08238 | HS90B | Heat shock protein HSP 90-beta | 0.71309845 | 0.070072375 | -0.783171075 | 28 | 14 | 11 | 37.4 | 19.5 | 16 | 83.263 | 949770000 | 87 |
| HSP90B1 | P14625 | ENPL | Endoplasmin | 0.817968225 | -0.072582125 | -0.745385925 | 46 | 46 | 43 | 51.4 | 51.4 | 49.7 | 92.468 | 5292300000 | 357 |
| HSPA1B | P0DMV9 | HS71B | Heat shock 70 kDa protein 1B | 0.56911135 | 0.45241826 | -1.02152962 | 30 | 26 | 13 | 55.2 | 49.6 | 25.7 | 70.051 | 1834600000 | 159 |
| HSPA5 | P11021 | BIP | Endoplasmic reticulum chaperone BiP | 0.126094525 | 0.085869575 | -0.211964293 | 46 | 46 | 44 | 60.1 | 60.1 | 59 | 72.332 | 10739000000 | 466 |
| HSPA8 | P11142 | HSP7C | Heat shock cognate 71 kDa protein | 0.443321275 | -0.222541623 | -0.220779575 | 44 | 42 | 16 | 66.1 | 65 | 28.9 | 70.897 | 14771000000 | 618 |
| HSPA9 | P38646 | GRP75 | Stress-70 protein, mitochondrial | 0.7180592 | 0.2657048 | -0.983763875 | 21 | 20 | 20 | 39.2 | 38.1 | 38.1 | 73.68 | 713660000 | 96 |
| HSPB1 | P04792 | HSPB1 | Heat shock protein beta-1 | -0.09244674 | -0.096869025 | 0.189315825 | 20 | 20 | 20 | 90.7 | 90.7 | 90.7 | 22.782 | 3055200000 | 160 |
| HSPD1 | P10809 | CH60 | 60 kDa heat shock protein, mitochondrial | 0.53356384 | 0.07984827 | -0.613412225 | 28 | 28 | 28 | 53.9 | 53.9 | 53.9 | 61.054 | 3910000000 | 197 |
| HSPE1 | P61604 | CH10 | 10 kDa heat shock protein, mitochondrial | 0.6587258 | 0.445486398 | -1.104212175 | 8 | 8 | 8 | 68.6 | 68.6 | 68.6 | 10.932 | 498190000 | 66 |
| HYOU1 | A0A087X054 | A0A087X054 | Hypoxia up-regulated protein 1 | 1.061106875 | -0.752955505 | -0.30815125 | 16 | 16 | 15 | 21.2 | 21.2 | 20.5 | 104.78 | 331200000 | 59 |
| IDH1 | O75874 | IDHC | Isocitrate dehydrogenase [NADP] cytoplasmic | 0.578528575 | -0.151993433 | -0.426535225 | 14 | 12 | 12 | 41.3 | 37.7 | 37.7 | 46.659 | 1093000000 | 40 |
| IDH2 | P48735 | IDHP | Isocitrate dehydrogenase [NADP], mitochondrial | 0.684313775 | 0.13735985 | -0.8216736 | 32 | 32 | 30 | 61.1 | 61.1 | 59.7 | 50.909 | 4355900000 | 230 |
| IGHA1 | P01876 | IGHA1 | Immunoglobulin heavy constant alpha 1 | -0.022081408 | 0.312576275 | -0.290494975 | 10 | 10 | 10 | 30.9 | 30.9 | 30.9 | 37.654 | 4240500000 | 213 |
| IGHG1 | P01857 | IGHG1 | Immunoglobulin heavy constant gamma 1 | -0.454714863 | -0.17177455 | 0.626489375 | 15 | 15 | 6 | 52.7 | 52.7 | 26.1 | 36.105 | 7367500000 | 337 |
| IGHG2 | P01859 | IGHG2 | Immunoglobulin heavy constant gamma 2 | 0.709473375 | -0.826314925 | 0.11684155 | 12 | 6 | 5 | 36.5 | 21.2 | 16.3 | 35.9 | 769450000 | 60 |
| IGHG4 | P01861 | IGHG4 | Immunoglobulin heavy constant gamma 4 | 0.338294515 | -0.597620685 | 0.259326125 | 12 | 7 | 5 | 42.5 | 23.5 | 15.6 | 35.94 | 398400000 | 43 |
| IGHM | P01871 | IGHM | Immunoglobulin heavy constant mu | -0.22593267 | -0.099908575 | 0.325841125 | 17 | 17 | 17 | 33.8 | 33.8 | 33.8 | 49.439 | 2586700000 | 212 |
| IGHV3OR16-9 | S4R460 | S4R460 | Immunoglobulin heavy variable 3/OR16-9 | -0.028240975 | -0.122476475 | 0.150717375 | 2 | 2 | 1 | 31.2 | 31.2 | 19.8 | 10.429 | 119890000 | 43 |
| IGKC | P01834 | IGKC | Immunoglobulin kappa constant | -0.468455025 | 0.252724478 | 0.215730425 | 9 | 9 | 9 | 93.5 | 93.5 | 93.5 | 11.765 | 4092000000 | 190 |
| IGKV3-15 | P01624 | KV315 | Immunoglobulin kappa variable 3-15 | -0.147447125 | -0.3070462 | 0.454493225 | 2 | 2 | 1 | 13.9 | 13.9 | 7.8 | 12.496 | 248890000 | 24 |
| IGKV3-20 | P01619 | KV320 | Immunoglobulin kappa variable 3-20 | -0.41589105 | -0.09741145 | 0.513302448 | 3 | 3 | 3 | 27.6 | 27.6 | 27.6 | 12.557 | 219770000 | 25 |
| IGLC3 | P0DOY3 | IGLC3 | Immunoglobulin lambda constant 3 | -0.557098808 | 0.384911295 | 0.1721876 | 7 | 7 | 3 | 67 | 67 | 20.8 | 11.265 | 7282600000 | 132 |
| IGLL5 | B9A064 | IGLL5 | Immunoglobulin lambda-like polypeptide 5 | -0.980255275 | -0.02746135 | 1.007716725 | 6 | 3 | 3 | 33.6 | 17.3 | 17.3 | 23.063 | 1978200000 | 50 |
| ILK | Q13418 | ILK | Integrin-linked protein kinase | 0.24170478 | -0.132021075 | -0.1096838 | 29 | 29 | 29 | 57.1 | 57.1 | 57.1 | 51.419 | 25701000000 | 663 |
| INF2 | Q27J81 | INF2 | Inverted formin-2 | 0.9142228 | -0.0609991 | -0.85322375 | 26 | 26 | 25 | 25.3 | 25.3 | 24.8 | 135.62 | 979110000 | 118 |
| IQGAP2 | Q13576 | IQGA2 | Ras GTPase-activating-like protein IQGAP2 | 0.624448025 | -0.265189975 | -0.359257978 | 59 | 59 | 57 | 40.8 | 40.8 | 39.7 | 180.58 | 1813400000 | 265 |
| ITGA2 | P17301 | ITA2 | Integrin alpha-2 | 0.78011805 | -0.260425873 | -0.51969195 | 16 | 16 | 16 | 17.4 | 17.4 | 17.4 | 129.29 | 405740000 | 65 |
| ITGA2B | P08514 | ITA2B | Integrin alpha-IIb | 0.858075775 | -0.064105475 | -0.793970325 | 46 | 46 | 46 | 42.2 | 42.2 | 42.2 | 113.38 | 57192000000 | 1172 |
| ITGA6 | P23229 | ITA6 | Integrin alpha-6 | 0.68859065 | 0.018647225 | -0.707237775 | 34 | 34 | 34 | 35.3 | 35.3 | 35.3 | 126.6 | 2020500000 | 214 |
| ITGB1 | P05556 | ITB1 | Integrin beta-1 | 0.292951168 | 0.365645025 | -0.658596125 | 13 | 13 | 13 | 19.2 | 19.2 | 19.2 | 88.414 | 1604800000 | 130 |
| ITGB3 | P05106 | ITB3 | Integrin beta-3 | 1.114821 | -0.529527275 | -0.58529376 | 35 | 35 | 34 | 42 | 42 | 42 | 87.057 | 44928000000 | 621 |
| ITIH1 | P19827 | ITIH1 | Inter-alpha-trypsin inhibitor heavy chain H1 | -0.587333343 | -0.00944615 | 0.5967795 | 18 | 18 | 18 | 20.1 | 20.1 | 20.1 | 101.39 | 996120000 | 122 |
| ITIH2 | Q5T985 | Q5T985 | Inter-alpha-trypsin inhibitor heavy chain H2 | -0.881384525 | 0.126525978 | 0.754858775 | 30 | 30 | 30 | 27.4 | 27.4 | 27.4 | 105.21 | 5276600000 | 285 |
| ITIH3 | A0A087WW43 | A0A087WW43 | Inter-alpha-trypsin inhibitor heavy chain H3 | -0.548327269 | 0.150653545 | 0.39767365 | 3 | 3 | 3 | 4.8 | 4.8 | 4.8 | 75.077 | 13712000 | 10 |
| ITIH4 | Q14624 | ITIH4 | Inter-alpha-trypsin inhibitor heavy chain H4 | -1.222240325 | 0.327501456 | 0.894738925 | 28 | 28 | 10 | 30.5 | 30.5 | 10.4 | 103.36 | 2384300000 | 264 |
| JAM3 | Q9BX67 | JAM3 | Junctional adhesion molecule C | 0.4212453 | 0.41075305 | -0.83199815 | 4 | 4 | 4 | 12.9 | 12.9 | 12.9 | 35.02 | 229080000 | 24 |
| JCHAIN | C9JA05 | C9JA05 | Immunoglobulin J chain | -0.59728135 | 0.3553835 | 0.24189785 | 3 | 3 | 3 | 38.6 | 38.6 | 38.6 | 8.1675 | 35149000 | 13 |
| KALRN | H7BXZ5 | H7BXZ5 | Kalirin | 0.703877703 | -0.1637851 | -0.540092575 | 16 | 16 | 16 | 6.9 | 6.9 | 6.9 | 336.17 | 239090000 | 52 |
| KIF2A | O00139 | KIF2A | Kinesin-like protein KIF2A | 0.16845629 | -0.051723975 | -0.11673237 | 20 | 20 | 19 | 27.3 | 27.3 | 26.1 | 79.954 | 541310000 | 92 |
| KNG1 | P01042 | KNG1 | Kininogen-1 | 0.273482835 | 0.0530757 | -0.326558575 | 13 | 13 | 13 | 19.6 | 19.6 | 19.6 | 71.957 | 706690000 | 70 |
| KPNB1 | Q14974 | IMB1 | Importin subunit beta-1 | -0.01046685 | -0.128173375 | 0.138640275 | 17 | 17 | 17 | 22.9 | 22.9 | 22.9 | 97.169 | 582610000 | 66 |
| KRT2 | P35908 | K22E | Keratin, type II cytoskeletal 2 epidermal | -0.362852598 | -0.220656075 | 0.583508275 | 50 | 46 | 1 | 75 | 71.2 | 4.7 | 65.432 | 10753000000 | 538 |
| LAMTOR2 | Q9Y2Q5 | LTOR2 | Ragulator complex protein LAMTOR2 | -0.56805398 | 1.17512795 | -0.607073975 | 4 | 4 | 4 | 43.2 | 43.2 | 43.2 | 13.507 | 24508000 | 8 |
| LASP1 | Q14847 | LASP1 | LIM and SH3 domain protein 1 | 0.68502 | 0.177208125 | -0.862228318 | 15 | 15 | 14 | 48.3 | 48.3 | 46 | 29.717 | 3114600000 | 124 |
| LCP2 | A0A0D9SGG1 | A0A0D9SGG1 | Lymphocyte cytosolic protein 2 | 0.7009095 | -0.061929525 | -0.638979375 | 16 | 16 | 16 | 26.3 | 26.3 | 26.3 | 60.143 | 649160000 | 54 |
| LDHA | P00338 | LDHA | L-lactate dehydrogenase A chain | 1.160879225 | -0.774044712 | -0.38683435 | 21 | 21 | 19 | 49.4 | 49.4 | 44 | 36.688 | 6151800000 | 214 |
| LDHB | P07195 | LDHB | L-lactate dehydrogenase B chain | 0.913466925 | -0.35262896 | -0.560838075 | 17 | 15 | 15 | 46.4 | 41 | 41 | 36.638 | 10259000000 | 293 |
| LGALSL | Q3ZCW2 | LEGL | Galectin-related protein | 0.50470555 | 0.421267475 | -0.925972975 | 10 | 10 | 10 | 70.3 | 70.3 | 70.3 | 18.986 | 1581800000 | 64 |
| LIMS1 | P48059 | LIMS1 | LIM and senescent cell antigen-like-containing domain protein 1 | 0.04978205 | 0.2572637 | -0.3070456 | 19 | 19 | 19 | 49.8 | 49.8 | 49.8 | 37.251 | 4657300000 | 154 |
| LRBA | P50851 | LRBA | Lipopolysaccharide-responsive and beige-like anchor protein | 1.197871075 | -0.6610374 | -0.536833968 | 39 | 39 | 39 | 15.1 | 15.1 | 15.1 | 319.1 | 738040000 | 119 |
| LTBP1 | Q14766 | LTBP1 | Latent-transforming growth factor beta-binding protein 1 | 0.80437955 | -0.48337915 | -0.321000493 | 30 | 30 | 30 | 19 | 19 | 19 | 186.79 | 5893800000 | 220 |
| LY6G6F | Q5SQ64 | LY66F | Lymphocyte antigen 6 complex locus protein G6f | 0.49753445 | -0.502631975 | 0.0050972 | 6 | 6 | 6 | 19.9 | 19.9 | 19.9 | 32.464 | 912210000 | 58 |
| LYN | P07948 | LYN | Tyrosine-protein kinase Lyn | 0.570277855 | -1.10551206 | 0.5352342 | 16 | 14 | 13 | 36.9 | 34 | 32 | 58.573 | 389300000 | 60 |
| MAOB | P27338 | AOFB | Amine oxidase [flavin-containing] B | 0.7860272 | -0.21956815 | -0.5664589 | 16 | 16 | 16 | 32.7 | 32.7 | 32.7 | 58.762 | 557800000 | 66 |
| MAPRE1 | Q15691 | MARE1 | Microtubule-associated protein RP/EB family member 1 | 1.1244785 | -0.77367035 | -0.350807995 | 14 | 14 | 13 | 43.7 | 43.7 | 43.7 | 29.999 | 647420000 | 75 |
| MAPRE2 | Q15555 | MARE2 | Microtubule-associated protein RP/EB family member 2 | -0.33292415 | -0.576238835 | 0.909162925 | 18 | 18 | 4 | 45.3 | 45.3 | 15 | 37.031 | 4090100000 | 154 |
| MAVS | Q7Z434 | MAVS | Mitochondrial antiviral-signaling protein | 0.8243974 | -0.27396345 | -0.5504339 | 8 | 8 | 8 | 24.1 | 24.1 | 24.1 | 56.527 | 358420000 | 48 |
| MDH1 | P40925 | MDHC | Malate dehydrogenase, cytoplasmic | 0.5635347 | -0.132011218 | -0.431523625 | 11 | 11 | 11 | 40.4 | 40.4 | 40.4 | 36.426 | 697450000 | 58 |
| MDH2 | P40926 | MDHM | Malate dehydrogenase, mitochondrial | 0.839946475 | -0.119383433 | -0.720562925 | 15 | 15 | 15 | 56.8 | 56.8 | 56.8 | 35.503 | 1021900000 | 105 |
| ME2 | A0A1W2PQH3 | A0A1W2PQH3 | Malic enzyme | 1.107857575 | -0.704664075 | -0.403193365 | 17 | 17 | 16 | 35.5 | 35.5 | 33.8 | 61.257 | 638210000 | 71 |
| MLEC | Q14165 | MLEC | Malectin | 0.69821735 | -0.739429675 | 0.041212375 | 10 | 10 | 10 | 39.4 | 39.4 | 39.4 | 32.233 | 349180000 | 55 |
| MMRN1 | Q13201 | MMRN1 | Multimerin-1 | 1.00382565 | 0.13461291 | -1.138438475 | 51 | 51 | 50 | 40.2 | 40.2 | 39.7 | 138.11 | 15275000000 | 642 |
| MOB1A | Q9H8S9 | MOB1A | MOB kinase activator 1A | -0.33533175 | -0.309775824 | 0.645107833 | 5 | 5 | 5 | 25.9 | 25.9 | 25.9 | 25.079 | 306560000 | 23 |
| MPIG6B | A0A140T9L8 | A0A140T9L8 | C6orf25 | 0.42905867 | -0.200465425 | -0.22859325 | 9 | 9 | 2 | 39.7 | 39.7 | 9.7 | 24.903 | 2187600000 | 89 |
| MPP1 | Q00013 | EM55 | 55 kDa erythrocyte membrane protein | 0.071213 | -0.77175235 | 0.700539525 | 13 | 13 | 13 | 30.5 | 30.5 | 30.5 | 52.296 | 326060000 | 38 |
| MPST | P25325 | THTM | 3-mercaptopyruvate sulfurtransferase | 0.291163525 | -0.181972645 | -0.109190925 | 7 | 7 | 7 | 35.4 | 35.4 | 35.4 | 33.178 | 369420000 | 22 |
| MSN | P26038 | MOES | Moesin | 0.871115375 | -0.495394058 | -0.375721095 | 66 | 66 | 52 | 74.4 | 74.4 | 56.3 | 67.819 | 11396000000 | 509 |
| MT-CO2 | P00403 | COX2 | Cytochrome c oxidase subunit 2 | 1.15369605 | -0.74737855 | -0.406317505 | 7 | 7 | 7 | 35.2 | 35.2 | 35.2 | 25.565 | 857720000 | 38 |
| MTPN | P58546 | MTPN | Myotrophin | 0.715458825 | -0.54474655 | -0.17071252 | 7 | 7 | 7 | 50 | 50 | 50 | 12.895 | 1161800000 | 52 |
| MYADM | C9JJV6 | C9JJV6 | Myeloid-associated differentiation marker | 0.6577427 | -0.33554213 | -0.3222004 | 4 | 4 | 4 | 30.8 | 30.8 | 30.8 | 15.864 | 335550000 | 36 |
| MYCT1 | H0YDV5 | H0YDV5 | Myc target protein 1 | 0.13087401 | -0.302036243 | 0.1711621 | 3 | 3 | 3 | 18.5 | 18.5 | 18.5 | 24.246 | 184710000 | 19 |
| MYH14 | Q7Z406 | MYH14 | Myosin-14 | 0.17360705 | -0.3742035 | 0.2005962 | 19 | 3 | 3 | 7.9 | 1.7 | 1.7 | 227.87 | 1905900000 | 55 |
| MYH9 | P35579 | MYH9 | Myosin-9 | 0.912330925 | -0.353166826 | -0.559164 | 230 | 230 | 185 | 76.8 | 76.8 | 61.8 | 226.53 | 2.2406E+11 | 4080 |
| MYL1 | P05976 | MYL1 | Myosin light chain 1/3, skeletal muscle isoform | 0.39525357 | -0.245496275 | -0.14975709 | 2 | 2 | 2 | 8.2 | 8.2 | 8.2 | 21.145 | 502380000 | 31 |
| MYL12A | P19105 | ML12A | Myosin regulatory light chain 12A | 0.589969275 | 0.434207175 | -1.0241765 | 12 | 12 | 5 | 66.7 | 66.7 | 29.8 | 19.794 | 17956000000 | 474 |
| MYL6 | F8W1R7 | F8W1R7 | Myosin light polypeptide 6 | 0.4915483 | -0.36658311 | -0.12496545 | 11 | 11 | 1 | 66.9 | 66.9 | 8.3 | 16.29 | 25095000000 | 278 |
| MYL9 | P24844 | MYL9 | Myosin regulatory light polypeptide 9 | 0.852858975 | -0.3458186 | -0.5070405 | 11 | 4 | 4 | 66.3 | 29.1 | 29.1 | 19.827 | 2309400000 | 70 |
| MYLK | Q15746 | MYLK | Myosin light chain kinase, smooth muscle | 0.84344193 | -0.62711355 | -0.216328473 | 19 | 19 | 19 | 11 | 11 | 11 | 210.71 | 1059500000 | 94 |
| NAP1L1 | F8W020 | F8W020 | Nucleosome assembly protein 1-like 1 | 0.9221007 | -0.037621203 | -0.8844794 | 9 | 9 | 1 | 42.5 | 42.5 | 9.2 | 24.366 | 1660700000 | 64 |
| NAPA | P54920 | SNAA | Alpha-soluble NSF attachment protein | 0.9063533 | -0.136656463 | -0.7696968 | 11 | 11 | 11 | 48.1 | 48.1 | 48.1 | 33.232 | 382710000 | 54 |
| NBEAL2 | Q6ZNJ1 | NBEL2 | Neurobeachin-like protein 2 | 0.282069 | -0.061822695 | -0.220246225 | 21 | 21 | 21 | 9.6 | 9.6 | 9.6 | 302.51 | 272100000 | 39 |
| NCK2 | O43639 | NCK2 | Cytoplasmic protein NCK2 | 0.203189575 | 0.51757785 | -0.7207673 | 9 | 9 | 9 | 25.5 | 25.5 | 25.5 | 42.915 | 187030000 | 27 |
| NCKAP1 | Q9Y2A7 | NCKP1 | Nck-associated protein 1 | 0.0803932 | -0.249758325 | 0.169364925 | 17 | 17 | 17 | 16.8 | 16.8 | 16.8 | 128.79 | 356330000 | 42 |
| NCKAP1L | P55160 | NCKPL | Nck-associated protein 1-like | -0.036445225 | 0.36710565 | -0.3306604 | 9 | 9 | 9 | 10.1 | 10.1 | 10.1 | 128.15 | 146280000 | 18 |
| NDUFA4 | O00483 | NDUA4 | Cytochrome c oxidase subunit NDUFA4 | 0.416772848 | -0.6325851 | 0.21581213 | 4 | 4 | 4 | 46.9 | 46.9 | 46.9 | 9.3697 | 134960000 | 16 |
| NEXN | Q0ZGT2 | NEXN | Nexilin | 0.42503015 | -0.211510325 | -0.213519775 | 26 | 26 | 26 | 29 | 29 | 29 | 80.657 | 1292400000 | 108 |
| NID1 | P14543 | NID1 | Nidogen-1 | 0.9876231 | 0.01023225 | -0.997855525 | 23 | 23 | 22 | 18.7 | 18.7 | 18.7 | 136.38 | 746400000 | 79 |
| NIPSNAP3A | Q9UFN0 | NPS3A | Protein NipSnap homolog 3A | 0.420586983 | -0.009461988 | -0.41112515 | 12 | 12 | 12 | 57.5 | 57.5 | 57.5 | 28.466 | 221850000 | 28 |
| NME1-NME2 | Q32Q12 | Q32Q12 | Nucleoside diphosphate kinase | 0.778343 | -0.53119065 | -0.247152348 | 15 | 15 | 7 | 62 | 62 | 21.2 | 32.642 | 1751800000 | 93 |
| NNT | Q13423 | NNTM | NAD(P) transhydrogenase, mitochondrial | 0.97070945 | -0.1370576 | -0.833651995 | 26 | 26 | 26 | 30.4 | 30.4 | 30.4 | 113.89 | 899570000 | 124 |
| NPEPPS | E9PLK3 | E9PLK3 | Aminopeptidase | 0.7735031 | -0.212621825 | -0.560881425 | 14 | 14 | 14 | 17.5 | 17.5 | 17.5 | 102.99 | 268590000 | 48 |
| NRGN | Q92686 | NEUG | Neurogranin | 0.5871239 | -0.50979595 | -0.077327975 | 3 | 3 | 3 | 26.9 | 26.9 | 26.9 | 7.6184 | 3244200000 | 56 |
| NSF | I3L0N3 | I3L0N3 | Vesicle-fusing ATPase | -0.843591525 | 0.410793253 | 0.43279816 | 6 | 6 | 6 | 8 | 8 | 8 | 82.091 | 26328000 | 11 |
| NUTF2 | P61970 | NTF2 | Nuclear transport factor 2 | 0.254113025 | -0.06664805 | -0.187464775 | 3 | 3 | 3 | 24.4 | 24.4 | 24.4 | 14.478 | 141820000 | 18 |
| P2RX1 | P51575 | P2RX1 | P2X purinoceptor 1 | -0.541500568 | -0.012156608 | 0.5536572 | 3 | 3 | 3 | 8.3 | 8.3 | 8.3 | 44.98 | 89845000 | 16 |
| P4HB | P07237 | PDIA1 | Protein disulfide-isomerase | 0.863789325 | -0.105766838 | -0.758022275 | 32 | 32 | 32 | 52.6 | 52.6 | 52.6 | 57.116 | 3681600000 | 217 |
| PAK2 | Q13177 | PAK2 | Serine/threonine-protein kinase PAK 2 | -0.80725745 | -0.06934285 | 0.876600425 | 4 | 4 | 4 | 9.2 | 9.2 | 9.2 | 58.042 | 63884000 | 14 |
| PARK7 | Q99497 | PARK7 | Protein/nucleic acid deglycase DJ-1 | -0.292429725 | 0.4371979 | -0.144768125 | 11 | 11 | 11 | 62.4 | 62.4 | 62.4 | 19.891 | 1472500000 | 103 |
| PARVB | Q9HBI1 | PARVB | Beta-parvin | 0.634078575 | -0.19690165 | -0.437176775 | 17 | 17 | 15 | 42 | 42 | 39.8 | 41.714 | 4006000000 | 243 |
| PCBP1 | Q15365 | PCBP1 | Poly(rC)-binding protein 1 | 0.856536275 | -0.1874435 | -0.6690927 | 9 | 9 | 6 | 31.2 | 31.2 | 20.2 | 37.497 | 956850000 | 56 |
| PCMT1 | P22061 | PIMT | Protein-L-isoaspartate | 0.240205375 | -0.066116775 | -0.1740884 | 5 | 5 | 5 | 31.3 | 31.3 | 31.3 | 24.636 | 278790000 | 31 |
| PDCD10 | Q9BUL8 | PDC10 | Programmed cell death protein 10 | 0.3966836 | 0.767221325 | -1.163905175 | 8 | 8 | 8 | 35.4 | 35.4 | 35.4 | 24.701 | 380600000 | 43 |
| PDCD6IP | Q8WUM4 | PDC6I | Programmed cell death 6-interacting protein | 0.630588575 | -0.270492225 | -0.360096325 | 21 | 21 | 21 | 25.1 | 25.1 | 25.1 | 96.022 | 515200000 | 67 |
| PDE5A | O76074 | PDE5A | cGMP-specific 3',5'-cyclic phosphodiesterase | 0.567479455 | -0.102399275 | -0.465080325 | 28 | 28 | 28 | 31.2 | 31.2 | 31.2 | 99.984 | 863040000 | 120 |
| PDHB | P11177 | ODPB | Pyruvate dehydrogenase E1 component subunit beta, mitochondrial | -0.435861175 | 0.2223797 | 0.213481435 | 4 | 4 | 4 | 14.2 | 14.2 | 14.2 | 39.233 | 68723000 | 17 |
| PDIA3 | P30101 | PDIA3 | Protein disulfide-isomerase A3 | 0.70067775 | -0.206913475 | -0.49376415 | 37 | 37 | 29 | 64.8 | 64.8 | 53.3 | 56.782 | 8470500000 | 405 |
| PDIA4 | P13667 | PDIA4 | Protein disulfide-isomerase A4 | 0.9671617 | -0.81482145 | -0.152340475 | 24 | 24 | 24 | 38.8 | 38.8 | 38.8 | 72.932 | 592220000 | 82 |
| PDIA5 | Q14554 | PDIA5 | Protein disulfide-isomerase A5 | 0.449032025 | 0.059599125 | -0.508631025 | 15 | 15 | 15 | 29.9 | 29.9 | 29.9 | 59.594 | 345900000 | 42 |

| | | | | | | | | | | | | | | |
|---|---|---|---|---|---|---|---|---|---|---|---|---|---|---|
| PDIA6 | Q15084 | PDIA6 | Protein disulfide-isomerase A6 | 0.519856985 | -0.094000425 | -0.4258564 | 15 | 15 | 14 | 35.9 | 35.9 | 34.5 | 48.121 | 2243300000 | 155 |
| PDLIM1 | O00151 | PDLI1 | PDZ and LIM domain protein 1 | 0.4820474 | -0.562471975 | 0.0804245 | 23 | 23 | 23 | 85.4 | 85.4 | 85.4 | 36.071 | 11529000000 | 318 |
| PDLIM5 | Q96HC4 | PDLI5 | PDZ and LIM domain protein 5 | 0.974688 | -0.20982462 | -0.7648633 | 12 | 12 | 12 | 23.5 | 23.5 | 23.5 | 63.944 | 628100000 | 48 |
| PDLIM7 | Q9NR12 | PDLI7 | PDZ and LIM domain protein 7 | 0.21918099 | -0.353923175 | 0.13474215 | 16 | 16 | 16 | 37 | 37 | 37 | 49.844 | 1243100000 | 80 |
| PEBP1 | P30086 | PEBP1 | Phosphatidylethanolamine-binding protein 1 | 0.259365725 | 0.2209626 | -0.480328405 | 6 | 6 | 6 | 44.4 | 44.4 | 44.4 | 21.057 | 246690000 | 25 |
| PECAM1 | A0A075B738 | PECA1 | Platelet endothelial cell adhesion molecule | 0.497083075 | -0.509311105 | 0.012227925 | 30 | 30 | 1 | 45.1 | 45.1 | 2.2 | 82.521 | 2837400000 | 255 |
| PF4 | P02776 | PLF4 | Platelet factor 4 | 0.409732268 | 0.12340455 | -0.533136813 | 7 | 7 | 3 | 44.6 | 44.6 | 12.9 | 10.845 | 38655000000 | 139 |
| PFKP | Q01813 | PFKAP | ATP-dependent 6-phosphofructokinase, platelet type | 0.7681264 | -0.280115975 | -0.48801045 | 31 | 31 | 29 | 42.2 | 42.2 | 38.6 | 85.595 | 1699500000 | 150 |
| PFN1 | P07737 | PROF1 | Profilin-1 | 0.838741783 | -0.422784825 | -0.415956868 | 17 | 17 | 17 | 97.1 | 97.1 | 97.1 | 15.054 | 45637000000 | 515 |
| PGAM1 | P18669 | PGAM1 | Phosphoglycerate mutase 1 | 1.037444025 | -0.7291808 | -0.308263475 | 9 | 9 | 9 | 42.9 | 42.9 | 42.9 | 28.804 | 763320000 | 41 |
| PGD | P52209 | 6PGD | 6-phosphogluconate dehydrogenase, decarboxylating | 0.642149225 | -0.3271805 | -0.314968836 | 19 | 19 | 19 | 42.7 | 42.7 | 42.7 | 53.139 | 866020000 | 73 |
| PGK1 | P00558 | PGK1 | Phosphoglycerate kinase 1 | 0.4850669 | -0.447200205 | -0.037866675 | 29 | 29 | 29 | 71.9 | 71.9 | 71.9 | 44.614 | 5720300000 | 266 |
| PGLS | O95336 | 6PGL | 6-phosphogluconolactonase | -0.114733628 | 0.258338085 | -0.143604525 | 6 | 6 | 6 | 35.7 | 35.7 | 35.7 | 27.547 | 179640000 | 29 |
| PGM1 | P36871 | PGM1 | Phosphoglucomutase-1 | 0.2247666 | -0.3376105 | 0.112843905 | 21 | 21 | 21 | 41.6 | 41.6 | 41.6 | 61.448 | 933390000 | 87 |
| PGM2 | Q96G03 | PGM2 | Phosphoglucomutase-2 | 0.21722065 | 0.0254006 | -0.242621375 | 8 | 8 | 8 | 14.7 | 14.7 | 14.7 | 68.283 | 170360000 | 23 |
| PGRMC1 | O00264 | PGRC1 | Membrane-associated progesterone receptor component 1 | 0.81139495 | 0.2004652 | -1.011860175 | 8 | 8 | 7 | 25.1 | 25.1 | 20.5 | 21.671 | 309400000 | 36 |
| PGRMC2 | O15173 | PGRC2 | Membrane-associated progesterone receptor component 2 | -0.805083725 | 0.195310905 | 0.609773048 | 5 | 4 | 4 | 26 | 22 | 22 | 23.818 | 50712000 | 10 |
| PHB | P35232 | PHB | Prohibitin | 0.3361706 | 0.608419125 | -0.944589748 | 13 | 13 | 13 | 48.9 | 48.9 | 48.9 | 29.804 | 508310000 | 61 |
| PHB2 | J3KPX7 | J3KPX7 | Prohibitin-2 | 1.074368275 | -0.602008375 | -0.472360058 | 13 | 13 | 13 | 51 | 51 | 51 | 33.239 | 459080000 | 53 |
| PIP4K2A | P48426 | PI42A | Phosphatidylinositol 5-phosphate 4-kinase type-2 alpha | 0.31243755 | -0.070852175 | -0.24158535 | 15 | 15 | 9 | 36 | 36 | 23.2 | 46.224 | 1197500000 | 106 |
| PKHD1L1 | Q86WI1 | PKHL1 | Fibrocystin-L | -0.932900925 | 0.79426365 | 0.138637125 | 4 | 4 | 4 | 1.3 | 1.3 | 1.3 | 465.73 | 27625000 | 10 |
| PKM | P14618 | KPYM | Pyruvate kinase PKM | -0.097907875 | -0.669485875 | 0.767393725 | 47 | 47 | 3 | 78.2 | 78.2 | 7.3 | 57.936 | 23829000000 | 687 |
| PLEC | Q15149 | PLEC | Plectin | 0.699613343 | -0.385335275 | -0.3142782 | 89 | 89 | 88 | 22.4 | 22.4 | 22 | 531.78 | 1336200000 | 229 |
| PLEK | P08567 | PLEK | Pleckstrin | 0.68122045 | -0.84869835 | 0.167478063 | 21 | 21 | 21 | 48.6 | 48.6 | 48.6 | 40.124 | 8406700000 | 216 |
| PLG | P00747 | PLMN | Plasminogen | -0.240657993 | 0.4018733 | -0.16121545 | 13 | 13 | 11 | 14 | 14 | 12 | 90.568 | 698050000 | 79 |
| PLIN3 | O60664 | PLIN3 | Perilipin-3 | -0.262187363 | 0.1731479 | 0.089039475 | 10 | 10 | 10 | 39.2 | 39.2 | 39.2 | 47.074 | 120260000 | 26 |
| PLS3 | A0A0A0MSQ0 | A0A0A0MSQ0 | Plastin-3 | -0.34494595 | 0.443605175 | -0.098659275 | 10 | 10 | 7 | 17.3 | 17.3 | 11.8 | 69.334 | 89425000 | 22 |
| PNP | P00491 | PNPH | Purine nucleoside phosphorylase | 0.052162735 | 0.489116875 | -0.5412796 | 17 | 17 | 17 | 73 | 73 | 73 | 32.118 | 2051600000 | 119 |
| PON1 | P27169 | PON1 | Serum paraoxonase/arylesterase 1 | -0.925790925 | 0.124133478 | 0.801657353 | 9 | 9 | 9 | 29.3 | 29.3 | 29.3 | 39.731 | 399350000 | 46 |
| PPBP | P02775 | CXCL7 | Platelet basic protein | 0.918460225 | -0.198370164 | -0.720090208 | 9 | 9 | 9 | 53.9 | 53.9 | 53.9 | 13.894 | 46454000000 | 384 |
| PPIA | P62937 | PPIA | Peptidyl-prolyl cis-trans isomerase A | 0.5483266 | -0.65009235 | 0.101765825 | 15 | 14 | 13 | 81.8 | 78.2 | 74.5 | 18.012 | 10193000000 | 201 |
| PPIB | P23284 | PPIB | Peptidyl-prolyl cis-trans isomerase B | 0.59918045 | -0.5759138 | -0.023266425 | 17 | 17 | 16 | 63.4 | 63.4 | 60.6 | 23.742 | 5110700000 | 211 |
| PPIF | P30405 | PPIF | Peptidyl-prolyl cis-trans isomerase F, mitochondrial | 0.31487225 | -0.480008525 | 0.1651362 | 13 | 12 | 11 | 72.9 | 70 | 67.1 | 22.04 | 950410000 | 62 |
| PPP1CA | P62136 | PP1A | Serine/threonine-protein phosphatase PP1-alpha catalytic subunit | 0.12225455 | -0.130281803 | 0.0080271 | 12 | 12 | 6 | 30.6 | 30.6 | 16.1 | 37.512 | 301110000 | 36 |
| PPP1CB | P62140 | PP1B | Serine/threonine-protein phosphatase PP1-beta catalytic subunit | 0.122361475 | -0.742715825 | 0.620354425 | 12 | 6 | 6 | 34.9 | 20.2 | 20.2 | 37.186 | 113270000 | 32 |
| PPP2R1A | P30153 | 2AAA | Serine/threonine-protein phosphatase 2A 65 kDa regulatory subunit A alpha isoform | 0.8200571 | -0.1863468 | -0.633710275 | 13 | 13 | 13 | 28 | 28 | 28 | 65.308 | 292710000 | 47 |
| PRDX1 | Q06830 | PRDX1 | Peroxiredoxin-1 | 0.221242125 | 0.2328396 | -0.4540814 | 15 | 15 | 11 | 63.3 | 63.3 | 48.2 | 22.11 | 1433900000 | 127 |
| PRDX2 | P32119 | PRDX2 | Peroxiredoxin-2 | -0.1723507 | 0.3916258 | -0.219275208 | 8 | 7 | 7 | 40.4 | 34.8 | 34.8 | 21.892 | 234510000 | 37 |
| PRDX3 | P30048 | PRDX3 | Thioredoxin-dependent peroxide reductase, mitochondrial | 0.394675025 | -0.064178575 | -0.33049625 | 7 | 7 | 7 | 30.9 | 30.9 | 30.9 | 27.692 | 915280000 | 51 |
| PRDX4 | Q13162 | PRDX4 | Peroxiredoxin-4 | 0.27498175 | -0.253857 | -0.02112477 | 10 | 7 | 7 | 36.2 | 29.2 | 29.2 | 30.54 | 383750000 | 36 |
| PRDX5 | P30044 | PRDX5 | Peroxiredoxin-5, mitochondrial | 0.221680948 | -0.02779955 | -0.1938811 | 9 | 9 | 9 | 55.6 | 55.6 | 55.6 | 22.086 | 739240000 | 74 |
| PRDX6 | P30041 | PRDX6 | Peroxiredoxin-6 | 0.466087215 | -0.04694875 | -0.419138575 | 18 | 18 | 18 | 75 | 75 | 75 | 25.035 | 4299200000 | 205 |
| PRKACB | P22694 | KAPCB | cAMP-dependent protein kinase catalytic subunit beta | 0.8893808 | -0.69258005 | -0.1968008 | 10 | 10 | 4 | 23.6 | 23.6 | 10.8 | 40.622 | 387700000 | 25 |
| PRKAR1A | P10644 | KAP0 | cAMP-dependent protein kinase type I-alpha regulatory subunit | 0.3536485 | -0.582832923 | 0.22918435 | 16 | 16 | 16 | 44.1 | 44.1 | 44.1 | 42.981 | 459440000 | 58 |
| PRKAR2B | P31323 | KAP3 | cAMP-dependent protein kinase type II-beta regulatory subunit | 0.4321768 | 0.197989375 | -0.630166083 | 6 | 6 | 6 | 17.2 | 17.2 | 17.2 | 46.302 | 246240000 | 24 |
| PRKCB | P05771 | KPCB | Protein kinase C beta type | 0.795410575 | -0.4959939 | -0.299416825 | 13 | 13 | 9 | 20.6 | 20.6 | 16.8 | 76.868 | 392520000 | 49 |
| PRKCSH | P14314 | GLU2B | Glucosidase 2 subunit beta | 0.527499775 | -0.608118025 | 0.0806182 | 11 | 11 | 11 | 20.8 | 20.8 | 20.8 | 59.425 | 1075800000 | 67 |
| PROS1 | P07225 | PROS | Vitamin K-dependent protein S | 0.592970275 | -0.5915008 | -0.001469725 | 13 | 13 | 13 | 19.7 | 19.7 | 19.7 | 75.122 | 733070000 | 88 |
| PSMA3 | P25788 | PSA3 | Proteasome subunit alpha type-3 | 0.0320311 | 0.16315165 | -0.1951826 | 9 | 9 | 9 | 38.4 | 38.4 | 38.4 | 28.433 | 253420000 | 35 |
| PSMA4 | P25789 | PSA4 | Proteasome subunit alpha type-4 | 0.005878 | -0.348425325 | 0.342547478 | 5 | 5 | 5 | 17.6 | 17.6 | 17.6 | 29.483 | 192660000 | 25 |
| PSMA6 | G3V295 | G3V295 | Proteasome subunit alpha type | -0.44019355 | 0.0476063 | 0.39258715 | 5 | 5 | 5 | 26.6 | 26.6 | 26.6 | 22.842 | 208770000 | 25 |
| PSMD11 | O00231 | PSD11 | 26S proteasome non-ATPase regulatory subunit 11 | 0.5912932 | -0.442967025 | -0.148325975 | 8 | 8 | 8 | 21.1 | 21.1 | 21.1 | 47.463 | 117170000 | 28 |
| PSMD3 | O43242 | PSMD3 | 26S proteasome non-ATPase regulatory subunit 3 | 0.049512375 | -0.0979933 | 0.048480825 | 6 | 6 | 6 | 14.2 | 14.2 | 14.2 | 60.977 | 67942000 | 20 |
| PSME1 | Q06323 | PSME1 | Proteasome activator complex subunit 1 | -0.123918775 | -0.01554785 | 0.139466775 | 9 | 9 | 9 | 34.5 | 34.5 | 34.5 | 28.723 | 198910000 | 26 |
| PSTPIP2 | Q9H939 | PPIP2 | Proline-serine-threonine phosphatase-interacting protein 2 | 0.8058798 | -0.3131397 | -0.492740175 | 13 | 13 | 13 | 40.1 | 40.1 | 40.1 | 38.858 | 557520000 | 41 |
| PTBP1 | A6NLN1 | A6NLN1 | Polypyrimidine tract binding protein 1, isoform CRA_b | -0.5447239 | 0.16696985 | 0.377754175 | 6 | 6 | 6 | 17.1 | 17.1 | 17.1 | 56.51 | 74326000 | 20 |
| PTGS1 | P23219 | PGH1 | Prostaglandin G/H synthase 1 | 0.650280775 | -0.173457553 | -0.476823125 | 16 | 16 | 16 | 31.1 | 31.1 | 31.1 | 68.686 | 1003400000 | 87 |
| PTPA | A6PVN5 | A6PVN5 | Serine/threonine-protein phosphatase 2A activator | 0.174956038 | -0.63483565 | 0.459879423 | 7 | 7 | 7 | 21.3 | 21.3 | 21.3 | 37.375 | 226600000 | 27 |
| PTPN1 | P18031 | PTN1 | Tyrosine-protein phosphatase non-receptor type 1 | -0.384256325 | -0.265784888 | 0.650041168 | 9 | 9 | 9 | 21.1 | 21.1 | 21.1 | 49.966 | 139370000 | 22 |
| PTPN12 | Q05209 | PTN12 | Tyrosine-protein phosphatase non-receptor type 12 | 0.743120863 | -0.040189475 | -0.702931625 | 16 | 16 | 16 | 25.9 | 25.9 | 25.9 | 88.105 | 237520000 | 34 |
| PTPN18 | E7EMB8 | E7EMB8 | Tyrosine-protein phosphatase non-receptor type 18 | 0.348094075 | -0.46004705 | 0.11195305 | 3 | 3 | 3 | 16 | 16 | 16 | 28.399 | 454040000 | 26 |
| PTPN6 | P29350 | PTN6 | Tyrosine-protein phosphatase non-receptor type 6 | 0.908606225 | 0.092679803 | -1.001285875 | 26 | 26 | 26 | 52.3 | 52.3 | 52.3 | 67.56 | 2077900000 | 128 |
| PTPRJ | Q12913 | PTPRJ | Receptor-type tyrosine-protein phosphatase eta | 0.7202569 | -0.617284975 | -0.102971775 | 17 | 17 | 17 | 16.3 | 16.3 | 16.3 | 145.94 | 527310000 | 66 |
| PTTG1IP | A8MXQ1 | A8MXQ1 | Pituitary tumor-transforming gene 1 protein-interacting protein | 0.3226145 | -0.23726325 | -0.08535125 | 3 | 3 | 3 | 11.9 | 11.9 | 11.9 | 18.331 | 260960000 | 14 |
| PYGB | P11216 | PYGB | Glycogen phosphorylase, brain form | 0.72805935 | -0.3923603 | -0.335698808 | 34 | 34 | 25 | 44.1 | 44.1 | 34.3 | 96.695 | 1828800000 | 150 |
| PYGL | P06737 | PYGL | Glycogen phosphorylase, liver form | 0.408418213 | -0.404759773 | -0.003658375 | 30 | 23 | 22 | 37.8 | 30.9 | 29.6 | 97.147 | 413390000 | 52 |
| RAB10 | P61026 | RAB10 | Ras-related protein Rab-10 | 0.70754415 | -0.64345929 | -0.064084925 | 16 | 15 | 14 | 73.5 | 68 | 64.5 | 22.541 | 2341900000 | 72 |
| RAB11B | Q15907 | RB11B | Ras-related protein Rab-11B | 0.55190488 | 0.137471075 | -0.689375945 | 16 | 16 | 16 | 68.8 | 68.8 | 68.8 | 24.488 | 5974600000 | 211 |
| RAB14 | P61106 | RAB14 | Ras-related protein Rab-14 | 0.203450063 | -0.11849765 | -0.084952225 | 12 | 12 | 12 | 73 | 73 | 73 | 23.897 | 1483900000 | 91 |
| RAB1A | P62820 | RAB1A | Ras-related protein Rab-1A | 0.162889925 | 0.103702373 | -0.2665924 | 16 | 7 | 7 | 81.5 | 38 | 38 | 22.677 | 938280000 | 59 |
| RAB1B | Q9H0U4 | RAB1B | Ras-related protein Rab-1B | 0.596168048 | -0.282017875 | -0.31415025 | 17 | 17 | 8 | 83.1 | 83.1 | 46.8 | 22.171 | 7181400000 | 261 |
| RAB27B | O00194 | RB27B | Ras-related protein Rab-27B | 0.53636505 | 0.402553025 | -0.93891795 | 16 | 16 | 16 | 70.6 | 70.6 | 70.6 | 24.608 | 4607400000 | 177 |
| RAB32 | Q13637 | RAB32 | Ras-related protein Rab-32 | 0.9239611 | -0.308927275 | -0.615033575 | 13 | 13 | 10 | 65.8 | 65.8 | 53.8 | 24.997 | 402680000 | 50 |
| RAB5C | P51148 | RAB5C | Ras-related protein Rab-5C | 0.718514075 | -0.717213675 | -0.00130027 | 9 | 9 | 5 | 50.5 | 50.5 | 32.9 | 23.482 | 784300000 | 80 |
| RAB6A | P20340 | RAB6A | Ras-related protein Rab-6A | -0.2965205 | 0.6747864 | -0.378265888 | 12 | 4 | 3 | 56.2 | 22.1 | 17.8 | 23.593 | 191610000 | 23 |
| RAB6B | Q9NRW1 | RAB6B | Ras-related protein Rab-6B | 0.5735423 | -0.162614325 | -0.4109279 | 15 | 15 | 5 | 72.1 | 72.1 | 32.2 | 23.461 | 2458600000 | 180 |
| RAB7A | P51149 | RAB7A | Ras-related protein Rab-7a | 0.7416633 | -0.14993075 | -0.591732675 | 12 | 12 | 12 | 71.5 | 71.5 | 71.5 | 23.489 | 1811600000 | 97 |
| RAB8A | P61006 | RAB8A | Ras-related protein Rab-8A | 0.381536458 | -0.877425538 | 0.49588895 | 14 | 12 | 9 | 57.5 | 52.2 | 37.7 | 23.668 | 1546100000 | 74 |
| RAC1 | P63000 | RAC1 | Ras-related C3 botulinum toxin substrate 1 | 0.179538763 | 0.461100893 | -0.640639675 | 10 | 10 | 5 | 48.4 | 48.4 | 24 | 21.45 | 1616200000 | 116 |
| RAC2 | B1AH77 | B1AH77 | Ras-related C3 botulinum toxin substrate 2 | 0.6435312 | -0.376466325 | -0.26706498 | 9 | 4 | 4 | 57.4 | 25.7 | 25.7 | 16.775 | 762020000 | 23 |
| RALB | P11234 | RALB | Ras-related protein Ral-B | 0.70806215 | -0.27893625 | -0.429125675 | 11 | 11 | 8 | 36.4 | 36.4 | 25.2 | 23.408 | 1003800000 | 61 |
| RAN | P62826 | RAN | GTP-binding nuclear protein Ran | 0.1532306 | -0.4078528 | 0.254622175 | 10 | 10 | 10 | 44.9 | 44.9 | 44.9 | 24.423 | 632590000 | 42 |
| RAP1B | P61224 | RAP1B | Ras-related protein Rap-1b | 0.632196475 | -0.043869603 | -0.588326688 | 16 | 16 | 6 | 71.2 | 71.2 | 33.7 | 20.825 | 35589000000 | 329 |
| RAP2B | P61225 | RAP2B | Ras-related protein Rap-2b | 0.371987125 | -0.53417055 | 0.162183385 | 9 | 9 | 6 | 45.4 | 45.4 | 33.3 | 20.504 | 426020000 | 40 |
| RASA3 | Q14644 | RASA3 | Ras GTPase-activating protein 3 | 0.3414725 | -0.413450725 | 0.071978 | 27 | 27 | 27 | 38.1 | 38.1 | 38.1 | 95.698 | 560030000 | 81 |
| RASGRP2 | Q7LDG7 | GRP2 | RAS guanyl-releasing protein 2 | 1.0204063 | -0.236339225 | -0.784067175 | 17 | 17 | 17 | 34 | 34 | 34 | 69.248 | 455720000 | 52 |
| RDH11 | Q8TC12 | RDH11 | Retinol dehydrogenase 11 | 0.47474929 | -0.15614385 | -0.318605255 | 12 | 12 | 12 | 43.4 | 43.4 | 43.4 | 35.386 | 1373400000 | 73 |
| RENBP | A6NKZ2 | A6NKZ2 | N-acylglucosamine 2-epimerase | 0.21412525 | 0.30015945 | -0.514284725 | 8 | 8 | 8 | 21.3 | 21.3 | 21.3 | 47.065 | 260910000 | 27 |
| RHOA | P61586 | RHOA | Transforming protein RhoA | 0.941097325 | -0.4022544 | -0.538843075 | 10 | 10 | 5 | 45.6 | 45.6 | 19.2 | 21.768 | 1818800000 | 85 |
| RNH1 | P13489 | RINI | Ribonuclease inhibitor | -0.130968385 | 0.762332765 | -0.631364448 | 5 | 5 | 5 | 15.2 | 15.2 | 15.2 | 49.973 | 127520000 | 24 |
| ROCK2 | O75116 | ROCK2 | Rho-associated protein kinase 2 | 0.6089566 | -0.266975875 | -0.3419806 | 38 | 38 | 31 | 27.7 | 27.7 | 23.1 | 160.9 | 932890000 | 133 |
| RPN1 | P04843 | RPN1 | Dolichyl-diphosphooligosaccharide--protein glycosyltransferase subunit 1 | 0.949982425 | -0.628548008 | -0.321434595 | 27 | 27 | 27 | 58.5 | 58.5 | 58.5 | 68.569 | 994000000 | 100 |
| RPN2 | P04844 | RPN2 | Dolichyl-diphosphooligosaccharide--protein glycosyltransferase subunit 2 | 0.80190035 | 0.166791853 | -0.968692075 | 14 | 14 | 14 | 31.1 | 31.1 | 31.1 | 69.283 | 437010000 | 56 |
| RSU1 | Q15404 | RSU1 | Ras suppressor protein 1 | 0.395065125 | -0.200507598 | -0.194557475 | 20 | 20 | 20 | 46.9 | 46.9 | 46.9 | 31.54 | 10804000000 | 357 |
| RTN1 | Q16799 | RTN1 | Reticulon-1 | 0.90253025 | -0.526143025 | -0.376387045 | 9 | 9 | 9 | 13.1 | 13.1 | 13.1 | 83.617 | 368450000 | 33 |
| RTN2 | K7EMR7 | K7EMR7 | Reticulon | 0.59577 | -0.5897358 | -0.00603405 | 6 | 6 | 6 | 26.2 | 26.2 | 26.2 | 29.519 | 438420000 | 41 |
| RTN4 | Q9NQC3 | RTN4 | Reticulon-4 | 0.5636052 | -0.037958025 | -0.525647125 | 9 | 9 | 1 | 12 | 12 | 2 | 129.93 | 3516100000 | 169 |
| S100A4 | P26447 | S10A4 | Protein S100-A4 | 0.449548725 | -0.50706215 | 0.057513225 | 8 | 8 | 8 | 52.5 | 52.5 | 52.5 | 11.728 | 1236600000 | 51 |
| S100A6 | R4GN98 | R4GN98 | Protein S100 | -0.154810975 | 0.161552425 | -0.006741325 | 5 | 5 | 5 | 37.6 | 37.6 | 37.6 | 9.681 | 654020000 | 36 |
| S100A9 | P06702 | S10A9 | Protein S100-A9 | 0.2011688 | 0.1233429 | -0.324511875 | 6 | 6 | 6 | 43.9 | 43.9 | 43.9 | 13.242 | 110500000 | 20 |
| SAA4 | P35542 | SAA4 | Serum amyloid A-4 protein | -0.969327275 | 0.3399818 | 0.629345425 | 4 | 4 | 4 | 27.7 | 27.7 | 27.7 | 14.746 | 104090000 | 23 |
| SACM1L | Q9NTJ5 | SAC1 | Phosphatidylinositide phosphatase SAC1 | 0.764151997 | -0.0388287 | -0.7253233 | 20 | 20 | 20 | 33.6 | 33.6 | 33.6 | 66.966 | 1146200000 | 118 |
| SAR1A | Q9NR31 | SAR1A | GTP-binding protein SAR1a | 0.5831666 | -0.057235125 | -0.525931425 | 10 | 10 | 10 | 36.4 | 36.4 | 36.4 | 22.367 | 1349000000 | 43 |
| SCAMP2 | O15127 | SCAM2 | Secretory carrier-associated membrane protein 2 | 0.237262175 | 0.0333943 | -0.270656638 | 3 | 3 | 3 | 13.7 | 13.7 | 13.7 | 36.648 | 195950000 | 21 |
| SCCPDH | Q8NBX0 | SCPDL | Saccharopine dehydrogenase-like oxidoreductase | -0.032748225 | -0.2579858 | 0.290733853 | 5 | 5 | 5 | 15.9 | 15.9 | 15.9 | 47.151 | 120050000 | 25 |
| SEC22B | O75396 | SC22B | Vesicle-trafficking protein SEC22b | -0.422891125 | -0.00109285 | 0.4239838 | 8 | 8 | 8 | 42.3 | 42.3 | 42.3 | 24.593 | 372090000 | 41 |
| SELP | Q5R345 | Q5R345 | P-selectin | 0.656454175 | 0.27543905 | -0.931893285 | 12 | 12 | 12 | 13.8 | 13.8 | 13.8 | 84.33 | 858350000 | 62 |
| SEPTIN11 | D6RGI3 | D6RGI3 | Septin 11 | -0.191411905 | 0.49132515 | -0.299913045 | 14 | 7 | 7 | 38.4 | 24 | 24 | 49.005 | 243770000 | 31 |
| SEPTIN2 | Q15019 | Sep-02 | Septin-2 | -0.032723475 | -0.53365185 | 0.5663754 | 14 | 14 | 8 | 49.3 | 49.3 | 34.9 | 41.487 | 653140000 | 54 |
| SEPTIN5 | G3XAH0 | G3XAH0 | HCG2002594, isoform CRA_c | 0.3868785 | -0.490779625 | 0.103901155 | 14 | 14 | 13 | 40.5 | 40.5 | 38.4 | 43.845 | 502450000 | 63 |
| SEPTIN6 | B1AMS2 | B1AMS2 | Septin 6 | 0.54768235 | -0.549012905 | 0.0013307 | 14 | 14 | 7 | 33.4 | 33.4 | 19.3 | 49.303 | 921460000 | 54 |
| SEPTIN7 | E7ES33 | E7ES33 | Septin 7 | 0.686629925 | -0.695621875 | 0.008991725 | 17 | 17 | 16 | 45.6 | 45.6 | 43.9 | 48.715 | 1860800000 | 111 |
| SERPINA1 | A0A024R6I7 | A0A024R6I7 | Alpha-1-antitrypsin | 0.733763225 | -0.5483699 | -0.18539325 | 25 | 25 | 1 | 55.3 | 55.3 | 3.8 | 46.708 | 3603500000 | 235 |
| SERPINA3 | P01011 | AACT | Alpha-1-antichymotrypsin | 0.27234305 | -0.4826693 | 0.210326465 | 11 | 11 | 11 | 24.3 | 24.3 | 24.3 | 47.65 | 291900000 | 38 |
| SERPINB1 | P30740 | ILEU | Leukocyte elastase inhibitor | 0.5389192 | -0.108716178 | -0.4302029 | 20 | 20 | 19 | 52.2 | 52.2 | 49.6 | 42.741 | 3328200000 | 183 |
| SERPINB6 | P35237 | SPB6 | Serpin B6 | 0.562419925 | -0.331874158 | -0.230546025 | 12 | 12 | 11 | 44.9 | 44.9 | 42.8 | 42.621 | 498360000 | 54 |
| SERPINB9 | P50453 | SPB9 | Serpin B9 | -0.1140735 | 0.716059825 | -0.601985975 | 16 | 15 | 15 | 49.5 | 46.8 | 46.8 | 42.403 | 281710000 | 34 |
| SERPIND1 | P05546 | HEP2 | Heparin cofactor 2 | -0.867371725 | 0.116918175 | 0.7504535 | 16 | 16 | 16 | 30.5 | 30.5 | 30.5 | 57.07 | 521270000 | 133 |
| SERPINE1 | P05121 | PAI1 | Plasminogen activator inhibitor 1 | 0.3155053 | 0.03205065 | -0.347556225 | 12 | 12 | 12 | 40.5 | 40.5 | 40.5 | 45.059 | 442790000 | 34 |
| SERPINF2 | P08697 | A2AP | Alpha-2-antiplasmin | -0.045212593 | -0.523009775 | 0.568222075 | 6 | 6 | 6 | 14.7 | 14.7 | 14.7 | 54.565 | 262390000 | 30 |
| SERPING1 | P05155 | IC1 | Plasma protease C1 inhibitor | 0.658859498 | -0.26482715 | -0.3940322 | 8 | 8 | 8 | 16.4 | 16.4 | 16.4 | 55.154 | 282890000 | 47 |
| SETDB1 | E9PAP1 | E9PAP1 | Histone-lysine N-methyltransferase SETDB1 | 0.6857831 | -0.52604625 | -0.159736873 | 1 | 1 | 1 | 10.8 | 10.8 | 10.8 | 12.422 | 627590000 | 10 |
| SH3BGRL | O75368 | SH3L1 | SH3 domain-binding glutamic acid-rich-like protein | 0.793503075 | 0.031713825 | -0.825217025 | 9 | 9 | 9 | 85.1 | 85.1 | 85.1 | 12.774 | 782310000 | 52 |
| SH3BGRL3 | Q5T123 | Q5T123 | SH3 domain-binding glutamic acid-rich-like protein 3 | -0.544413075 | -0.030175125 | 0.574588175 | 7 | 7 | 7 | 54.5 | 54.5 | 54.5 | 9.3804 | 6296600000 | 78 |
| SKAP2 | O75563 | SKAP2 | Src kinase-associated phosphoprotein 2 | 0.47269435 | -0.40482325 | -0.067871325 | 9 | 9 | 9 | 26.7 | 26.7 | 26.7 | 41.216 | 324800000 | 42 |
| SLC25A11 | I3L1P8 | I3L1P8 | Mitochondrial 2-oxoglutarate/malate carrier protein | 0.4165061 | -0.304553915 | -0.11195215 | 11 | 11 | 11 | 39.2 | 39.2 | 39.2 | 32.182 | 209240000 | 40 |
| SLC25A3 | Q00325 | MPCP | Phosphate carrier protein, mitochondrial | 1.004866725 | -0.6569613 | -0.347905425 | 14 | 14 | 14 | 35.6 | 35.6 | 35.6 | 40.094 | 1382700000 | 84 |
| SLC25A5 | P05141 | ADT2 | ADP/ATP translocase 2 | 0.7892635 | 0.06351137 | -0.85277485 | 17 | 17 | 6 | 48.3 | 48.3 | 19.5 | 32.852 | 3763500000 | 143 |
| SLC25A6 | P12236 | ADT3 | ADP/ATP translocase 3 | 0.330736975 | 0.348408125 | -0.6791453 | 15 | 4 | 4 | 41.9 | 13.1 | 13.1 | 32.866 | 271510000 | 21 |
| SLC2A3 | P11169 | GTR3 | Solute carrier family 2, facilitated glucose transporter member 3 | 0.702608975 | -0.5511396 | -0.151469575 | 12 | 12 | 11 | 18.8 | 18.8 | 18.8 | 53.924 | 7822500000 | 158 |
| SLC9A3R1 | O14745 | NHRF1 | Na(+)/H(+) exchange regulatory cofactor NHE-RF1 | -0.60439465 | 0.36369459 | 0.240700085 | 12 | 12 | 12 | 35.2 | 35.2 | 35.2 | 38.868 | 502330000 | 56 |
| SLC9A9 | Q8IVB4 | SL9A9 | Sodium/hydrogen exchanger 9 | -0.4070442 | 0.2118831 | 0.195161075 | 5 | 5 | 5 | 10.7 | 10.7 | 10.7 | 72.564 | 58107000 | 8 |
| SNAP23 | O00161 | SNP23 | Synaptosomal-associated protein 23 | 1.0689415 | -0.9344885 | -0.134452851 | 17 | 17 | 17 | 72.5 | 72.5 | 72.5 | 23.354 | 1533600000 | 120 |
| SNAP29 | O95721 | SNP29 | Synaptosomal-associated protein 29 | -0.10278815 | 0.0054948 | 0.097293309 | 5 | 5 | 5 | 20.5 | 20.5 | 20.5 | 28.97 | 63821000 | 18 |
| SNCA | E7EPV7 | E7EPV7 | Alpha-synuclein | 0.801465015 | -0.26265945 | -0.53880565 | 13 | 13 | 13 | 80.9 | 80.9 | 80.9 | 11.777 | 2231300000 | 144 |
| SND1 | Q7KZF4 | SND1 | Staphylococcal nuclease domain-containing protein 1 | 0.711363225 | -0.435411941 | -0.27595124 | 17 | 17 | 17 | 23.6 | 23.6 | 23.6 | 102 | 301580000 | 47 |
| SNX3 | O60493 | SNX3 | Sorting nexin-3 | 0.548043175 | -0.310731775 | -0.237311008 | 7 | 7 | 6 | 41.4 | 41.4 | 37 | 18.762 | 273150000 | 30 |
| SOD2 | P04179 | SODM | Superoxide dismutase [Mn], mitochondrial | 0.77016055 | 0.03965645 | -0.809817025 | 9 | 9 | 9 | 46.4 | 46.4 | 46.4 | 24.75 | 1210800000 | 104 |
| SPARC | P09486 | SPRC | SPARC | 0.66907535 | 0.00095735 | -0.670032425 | 14 | 14 | 14 | 32.3 | 32.3 | 32.3 | 34.632 | 1443000000 | 108 |
| SPTBN1 | Q01082 | SPTB2 | Spectrin beta chain, non-erythrocytic 1 | -0.140044975 | -0.407809775 | 0.547855075 | 47 | 47 | 44 | 23.3 | 23.3 | 22.1 | 274.61 | 1298800000 | 100 |
| SRC | P12931 | SRC | Proto-oncogene tyrosine-protein kinase Src | 0.71178984 | 0.183996025 | -0.895785913 | 24 | 24 | 17 | 53.7 | 53.7 | 42.2 | 59.834 | 3685300000 | 188 |
| SRGN | P10124 | SRGN | Serglycin | 0.34867165 | -0.215959075 | -0.132712625 | 5 | 5 | 5 | 19 | 19 | 19 | 17.652 | 809600000 | 39 |
| ST13 | P50502 | F10A1 | Hsc70-interacting protein | 0.95643325 | -0.7047006 | -0.251732525 | 7 | 7 | 7 | 20.6 | 20.6 | 20.6 | 41.331 | 224560000 | 39 |
| STIM1 | Q13586 | STIM1 | Stromal interaction molecule 1 | 1.1747832 | -0.399342315 | -0.775441125 | 18 | 18 | 18 | 28.2 | 28.2 | 28.2 | 77.422 | 1138700000 | 87 |
| STIP1 | P31948 | STIP1 | Stress-induced-phosphoprotein 1 | 0.142902575 | 0.191712695 | -0.334615098 | 21 | 21 | 21 | 41.6 | 41.6 | 41.6 | 62.639 | 822100000 | 55 |
| STK24 | B4DR80 | B4DR80 | Serine/threonine-protein kinase 24 | 0.251587825 | 0.09312975 | -0.344717889 | 12 | 12 | 8 | 34.7 | 34.7 | 26.5 | 45.835 | 324900000 | 40 |
| STOM | P27105 | STOM | Erythrocyte band 7 integral membrane protein | 0.5838584 | -0.151823775 | -0.432034675 | 20 | 20 | 20 | 63.9 | 63.9 | 63.9 | 31.73 | 20376000000 | 558 |
| STT3A | P46977 | STT3A | Dolichyl-diphosphooligosaccharide--protein glycosyltransferase subunit STT3A | 0.15942135 | -0.516400775 | 0.35697935 | 7 | 7 | 7 | 9.9 | 9.9 | 9.9 | 80.529 | 158050000 | 26 |
| STX11 | O75558 | STX11 | Syntaxin-11 | 0.5805426 | -0.178843475 | -0.401699175 | 12 | 12 | 12 | 48.8 | 48.8 | 48.8 | 33.195 | 925240000 | 77 |
| STX7 | O15400 | STX7 | Syntaxin-7 | 0.286244088 | -0.16180775 | -0.124436225 | 8 | 8 | 8 | 37.5 | 37.5 | 37.5 | 29.815 | 344900000 | 53 |
| STXBP2 | Q15833 | STXB2 | Syntaxin-binding protein 2 | 0.653325313 | -0.0303936 | -0.622931675 | 31 | 31 | 31 | 50.3 | 50.3 | 50.3 | 66.452 | 2333500000 | 170 |
| SYTL4 | Q96C24 | SYTL4 | Synaptotagmin-like protein 4 | 0.73397425 | -0.3756007 | -0.35837371 | 28 | 28 | 28 | 42.8 | 42.8 | 42.8 | 76.023 | 684030000 | 108 |
| TAGLN2 | P37802 | TAGL2 | Transgelin-2 | 0.6475213 | -0.21578135 | -0.431739918 | 20 | 20 | 19 | 77.9 | 77.9 | 70.9 | 22.391 | 57216000000 | 700 |
| TALDO1 | P37837 | TALDO | Transaldolase | 0.486568075 | 0.603691675 | -1.090259825 | 14 | 14 | 14 | 39.8 | 39.8 | 39.8 | 37.54 | 1326100000 | 75 |
| TAOK3 | Q9H2K8 | TAOK3 | Serine/threonine-protein kinase TAO3 | 0.58066985 | -0.712382425 | 0.1317125 | 6 | 6 | 6 | 7.1 | 7.1 | 7.1 | 105.4 | 102850000 | 19 |
| TARS | P26639 | SYTC | Threonine--tRNA ligase, cytoplasmic | -0.50483675 | 0.25344744 | 0.25138955 | 11 | 11 | 11 | 15.8 | 15.8 | 15.8 | 83.434 | 176220000 | 26 |
| TBXAS1 | P24557 | THAS | Thromboxane-A synthase | 0.477260625 | 0.29744625 | -0.774706925 | 21 | 21 | 21 | 41.7 | 41.7 | 41.7 | 60.518 | 1211700000 | 144 |
| TCP1 | P17987 | TCPA | T-complex protein 1 subunit alpha | -0.00892075 | -0.527563375 | 0.536484065 | 19 | 19 | 19 | 40.3 | 40.3 | 40.3 | 60.343 | 605520000 | 84 |
| TF | P02787 | TRFE | Serotransferrin | -0.1881008 | 0.624469825 | -0.4363691 | 27 | 27 | 26 | 38.8 | 38.8 | 37.8 | 77.063 | 2747900000 | 175 |
| TGFB1 | P01137 | TGFB1 | Transforming growth factor beta-1 proprotein | 0.643462475 | -0.031960225 | -0.611502375 | 13 | 13 | 13 | 32.6 | 32.6 | 32.6 | 44.341 | 1057600000 | 78 |
| TGFB1I1 | O43294 | TGFI1 | Transforming growth factor beta-1-induced transcript 1 protein | 0.00415965 | -0.312967625 | 0.308807895 | 7 | 7 | 7 | 16.5 | 16.5 | 16.5 | 49.814 | 324410000 | 23 |
| TJP2 | Q9UDY2 | ZO2 | Tight junction protein ZO-2 | 0.894168625 | -0.2813877 | -0.612780875 | 28 | 28 | 28 | 27.6 | 27.6 | 27.6 | 133.96 | 1233800000 | 96 |
| TKT | P29401 | TKT | Transketolase | 0.87675995 | -0.703869375 | -0.172890675 | 19 | 19 | 19 | 35.8 | 35.8 | 35.8 | 67.877 | 786140000 | 77 |
| TLN1 | Q9Y490 | TLN1 | Talin-1 | 0.274730925 | -0.02831005 | -0.246420875 | 210 | 210 | 180 | 77 | 77 | 70.6 | 269.76 | 3.6463E+11 | 5184 |
| TMED10 | P49755 | TMEDA | Transmembrane emp24 domain-containing protein 10 | 1.202276625 | -0.544220365 | -0.6580564 | 9 | 9 | 9 | 40.6 | 40.6 | 40.6 | 24.976 | 440810000 | 35 |
| TMEM40 | Q8WWA1 | TMM40 | Transmembrane protein 40 | 0.940832725 | -1.055802575 | 0.114969625 | 5 | 5 | 5 | 26.2 | 26.2 | 26.2 | 25.495 | 258390000 | 29 |
| TMEM63A | O94886 | CSCL1 | CSC1-like protein 1 | 0.46507335 | -0.53451995 | 0.0694466 | 8 | 8 | 8 | 9.5 | 9.5 | 9.5 | 92.125 | 161950000 | 26 |
| TMOD3 | Q9NYL9 | TMOD3 | Tropomodulin-3 | 0.9531226 | -0.563227625 | -0.38989505 | 13 | 13 | 13 | 50 | 50 | 50 | 39.594 | 811670000 | 85 |
| TMSB4X | P62328 | TYB4 | Thymosin beta-4 | 0.23997395 | 0.040561736 | -0.28053585 | 11 | 11 | 9 | 93.2 | 93.2 | 88.6 | 5.0526 | 39454000000 | 366 |
| TMX1 | Q9H3N1 | TMX1 | Thioredoxin-related transmembrane protein 1 | -0.26647705 | -0.184570125 | 0.45104715 | 7 | 7 | 7 | 23.6 | 23.6 | 23.6 | 31.791 | 232540000 | 40 |
| TOR4A | Q9NXH8 | TOR4A | Torsin-4A | 0.90907941 | -0.677001993 | -0.232077382 | 12 | 12 | 12 | 35.2 | 35.2 | 35.2 | 46.914 | 232990000 | 48 |
| TPD52L2 | O43399 | TPD54 | Tumor protein D54 | -0.40449923 | 0.83836925 | -0.433869833 | 6 | 6 | 6 | 32.5 | 32.5 | 32.5 | 22.237 | 228580000 | 36 |
| TPI1 | P60174 | TPIS | Triosephosphate isomerase | -0.378579275 | -0.517888845 | 0.896468125 | 18 | 18 | 18 | 75.5 | 75.5 | 75.5 | 30.791 | 6366100000 | 190 |
| TPM1 | H7BYY1 | H7BYY1 | Tropomyosin 1 (Alpha), isoform CRA_m | 0.412644045 | 0.165678975 | -0.578323038 | 42 | 27 | 10 | 81.9 | 62.9 | 25.4 | 28.747 | 11336000000 | 236 |
| TPM1 | Q6ZN40 | Q6ZN40 | Tropomyosin 1 (Alpha), isoform CRA_f | 0.440639795 | 0.087792 | -0.528431738 | 38 | 10 | 4 | 59.5 | 19.6 | 8 | 37.452 | 929040000 | 68 |
| TPM3 | J3KN67 | J3KN67 | Tropomyosin alpha-3 chain | 0.60502875 | -0.35638575 | -0.248642998 | 40 | 2 | 0 | 70.5 | 8.1 | 0 | 33.222 | 838690000 | 29 |

| | | | | | | | | | | | | | | | |
|---|---|---|---|---|---|---|---|---|---|---|---|---|---|---|---|
| TPM3 | Q5HYB6 | Q5HYB6 | Epididymis luminal protein 189 | 0.45106925 | -0.248624503 | -0.20244446 | 44 | 27 | 0 | 92.7 | 69.8 | 0 | 27.175 | 18260000000 | 409 |
| TPM4 | P67936 | TPM4 | Tropomyosin alpha-4 chain | 0.563655875 | -0.25792891 | -0.30572665 | 51 | 51 | 19 | 91.5 | 91.5 | 40.3 | 28.521 | 82677000000 | 823 |
| TPP1 | O14773 | TPP1 | Tripeptidyl-peptidase 1 | -0.2922073 | -0.05378735 | 0.345994725 | 4 | 4 | 4 | 9.4 | 9.4 | 9.4 | 61.247 | 152170000 | 24 |
| TPP2 | P29144 | TPP2 | Tripeptidyl-peptidase 2 | 0.933392125 | -0.983391425 | 0.04999935 | 27 | 27 | 27 | 23.9 | 23.9 | 23.9 | 138.35 | 625850000 | 61 |
| TPT1 | P13693 | TCTP | Translationally-controlled tumor protein | 0.744689833 | -0.508447585 | -0.236242128 | 7 | 7 | 7 | 40.1 | 40.1 | 40.1 | 19.595 | 623020000 | 33 |
| TREML1 | Q86YW5 | TRML1 | Trem-like transcript 1 protein | 0.14062113 | 0.158131975 | -0.2987533 | 9 | 9 | 9 | 49.2 | 49.2 | 49.2 | 32.678 | 1944300000 | 71 |
| TRIM58 | Q8NG06 | TRI58 | E3 ubiquitin-protein ligase TRIM58 | 0.1735609 | 0.102650925 | -0.276211675 | 8 | 8 | 8 | 20.4 | 20.4 | 20.4 | 54.766 | 207780000 | 25 |
| TST | Q16762 | THTR | Thiosulfate sulfurtransferase | -0.51174825 | 0.46007495 | 0.051673 | 8 | 8 | 8 | 25.3 | 25.3 | 25.3 | 33.429 | 86879000 | 17 |
| TUBA1B | P68363 | TBA1B | Tubulin alpha-1B chain | 0.12480278 | 0.297921085 | -0.4227238 | 25 | 25 | 0 | 61.4 | 61.4 | 0 | 50.151 | 38234000000 | 690 |
| TUBA4A | P68366 | TBA4A | Tubulin alpha-4A chain | 0.4798274 | -0.142054525 | -0.33777295 | 25 | 4 | 4 | 61.8 | 12.1 | 12.1 | 49.924 | 7234700000 | 149 |
| TUBA8 | Q9NY65 | TBA8 | Tubulin alpha-8 chain | 0.476113338 | 0.229548575 | -0.7056619 | 22 | 9 | 9 | 57.2 | 23.8 | 23.8 | 50.093 | 1282500000 | 95 |
| TUBB | Q5JP53 | Q5JP53 | Tubulin beta chain | -0.11286615 | -0.02253795 | 0.135404175 | 24 | 6 | 5 | 65.5 | 20.4 | 16.9 | 47.766 | 3383400000 | 88 |
| TUBB1 | Q9H4B7 | TBB1 | Tubulin beta-1 chain | 0.646058625 | -0.33527905 | -0.310779825 | 27 | 27 | 15 | 73.8 | 73.8 | 48.3 | 50.326 | 49082000000 | 777 |
| TUBB4B | P68371 | TBB4B | Tubulin beta-4B chain | 0.021600975 | -0.067637379 | 0.046036225 | 26 | 21 | 4 | 67 | 60.4 | 13 | 49.83 | 16362000000 | 399 |
| TUFM | P49411 | EFTU | Elongation factor Tu, mitochondrial | 1.2314209 | -0.390003228 | -0.8414177 | 12 | 12 | 12 | 33 | 33 | 33 | 49.541 | 442060000 | 39 |
| TWF2 | Q6IBS0 | TWF2 | Twinfilin-2 | 1.01762885 | -0.49441465 | -0.523214043 | 17 | 17 | 9 | 62.2 | 62.2 | 33.2 | 39.548 | 532450000 | 60 |
| TXN | P10599 | THIO | Thioredoxin | 0.2601695 | -0.04828175 | -0.21188755 | 7 | 7 | 7 | 51.4 | 51.4 | 51.4 | 11.737 | 507100000 | 39 |
| TXNL1 | O43396 | TXNL1 | Thioredoxin-like protein 1 | -0.32353245 | 0.3647566 | -0.041224325 | 8 | 8 | 8 | 33.9 | 33.9 | 33.9 | 32.251 | 144000000 | 23 |
| TYMP | P19971 | TYPH | Thymidine phosphorylase | 0.3690555 | -0.109808325 | -0.259247315 | 18 | 18 | 18 | 42.9 | 42.9 | 42.9 | 49.955 | 681680000 | 86 |
| UBA1 | P22314 | UBA1 | Ubiquitin-like modifier-activating enzyme 1 | 0.74750195 | -0.488800175 | -0.25870185 | 25 | 25 | 25 | 30.3 | 30.3 | 30.3 | 117.85 | 775160000 | 113 |
| UBA7 | P41226 | UBA7 | Ubiquitin-like modifier-activating enzyme 7 | -0.064903925 | 0.025094375 | 0.039809475 | 16 | 16 | 16 | 24.6 | 24.6 | 24.6 | 111.69 | 367310000 | 46 |
| UBASH3B | Q8TF42 | UBS3B | Ubiquitin-associated and SH3 domain-containing protein B | 0.527104575 | -0.23762105 | -0.289483339 | 16 | 16 | 16 | 30.7 | 30.7 | 30.7 | 72.695 | 736220000 | 51 |
| UBB | J3QS39 | J3QS39 | Polyubiquitin-B | 0.907819475 | -0.31073645 | -0.59708315 | 7 | 7 | 7 | 63.4 | 63.4 | 63.4 | 10.469 | 3048700000 | 72 |
| UBE2L3 | P68036 | UB2L3 | Ubiquitin-conjugating enzyme E2 L3 | 0.88316065 | -0.167374273 | -0.7157861 | 8 | 8 | 8 | 55.8 | 55.8 | 55.8 | 17.861 | 485920000 | 36 |
| UBE2N | P61088 | UBE2N | Ubiquitin-conjugating enzyme E2 N | 0.493015443 | -0.2679012 | -0.225114025 | 7 | 7 | 7 | 52 | 52 | 52 | 17.138 | 317160000 | 27 |
| UBE2O | Q9C0C9 | UBE2O | (E3-independent) E2 ubiquitin-conjugating enzyme | -0.503832425 | -0.3064669 | 0.8102995 | 10 | 10 | 10 | 9.5 | 9.5 | 9.5 | 141.29 | 186390000 | 24 |
| UGGT1 | Q9NYU2 | UGGG1 | UDP-glucose:glycoprotein glucosyltransferase 1 | 0.1244882 | 0.09209575 | -0.2165837 | 19 | 19 | 19 | 15.9 | 15.9 | 15.9 | 177.19 | 307640000 | 61 |
| UGP2 | A0A087WYS1 | A0A087WYS1 | UTP--glucose-1-phosphate uridylyltransferase | -0.262770375 | -0.053954725 | 0.31672496 | 19 | 19 | 9 | 39.6 | 39.6 | 18.1 | 56.967 | 941160000 | 70 |
| UNC13D | Q70J99 | UN13D | Protein unc-13 homolog D | 0.914734475 | -0.093591242 | -0.82114352 | 31 | 31 | 22 | 37.1 | 37.1 | 26.6 | 123.28 | 1955400000 | 169 |
| UQCRC2 | P22695 | QCR2 | Cytochrome b-c1 complex subunit 2, mitochondrial | -0.179451025 | -0.211742538 | 0.391193675 | 13 | 13 | 13 | 37.3 | 37.3 | 37.3 | 48.442 | 325290000 | 44 |
| USP14 | A6NJA2 | A6NJA2 | Ubiquitin carboxyl-terminal hydrolase 14 | -0.416099725 | 0.231847025 | 0.18425285 | 8 | 8 | 8 | 23 | 23 | 23 | 51.086 | 120780000 | 18 |
| VAMP3 | Q15836 | VAMP3 | Vesicle-associated membrane protein 3 | 0.769548 | -0.572637368 | -0.196910625 | 6 | 6 | 6 | 53 | 53 | 53 | 11.309 | 199840000 | 27 |
| VAMP8 | Q9BV40 | VAMP8 | Vesicle-associated membrane protein 8 | 0.141870325 | -0.4063649 | 0.264494725 | 4 | 4 | 4 | 41 | 41 | 41 | 11.438 | 231100000 | 27 |
| VAPA | Q9P0L0 | VAPA | Vesicle-associated membrane protein-associated protein A | 0.4441764 | -0.481526827 | 0.0373505 | 9 | 9 | 8 | 43 | 43 | 38.2 | 27.893 | 745380000 | 31 |
| VASP | P50552 | VASP | Vasodilator-stimulated phosphoprotein | 0.46887745 | -0.023748503 | -0.44512906 | 22 | 22 | 22 | 51.3 | 51.3 | 51.3 | 39.829 | 8955600000 | 253 |
| VCL | P18206 | VINC | Vinculin | 0.13868875 | -0.084919585 | -0.053769125 | 117 | 117 | 117 | 79.3 | 79.3 | 79.3 | 123.8 | 1.1783E+11 | 2253 |
| VCP | P55072 | TERA | Transitional endoplasmic reticulum ATPase | 0.527478475 | -0.33961589 | -0.187862575 | 38 | 38 | 38 | 44.5 | 44.5 | 44.5 | 89.321 | 2595100000 | 223 |
| VDAC1 | P21796 | VDAC1 | Voltage-dependent anion-selective channel protein 1 | 0.81561435 | -0.2870123 | -0.52860205 | 14 | 14 | 14 | 59.4 | 59.4 | 59.4 | 30.772 | 830930000 | 78 |
| VDAC2 | P45880 | VDAC2 | Voltage-dependent anion-selective channel protein 2 | 0.8161434 | 0.024924545 | -0.84106792 | 11 | 11 | 11 | 40.1 | 40.1 | 40.1 | 31.566 | 1141000000 | 96 |
| VDAC3 | Q9Y277 | VDAC3 | Voltage-dependent anion-selective channel protein 3 | 0.136229275 | 0.4486049 | -0.5848344 | 14 | 14 | 13 | 56.2 | 56.2 | 54.1 | 30.658 | 3406300000 | 126 |
| VIM | B0YJC4 | B0YJC4 | Vimentin | 0.4615867 | -0.2150061 | -0.246580675 | 12 | 11 | 10 | 25.5 | 23.9 | 21.6 | 49.653 | 137840000 | 21 |
| VPS35 | Q96QK1 | VPS35 | Vacuolar protein sorting-associated protein 35 | -0.590382775 | 0.411825848 | 0.1785567 | 9 | 9 | 9 | 14.4 | 14.4 | 14.4 | 91.706 | 148120000 | 22 |
| VTN | P04004 | VTNC | Vitronectin | 0.093940925 | -0.003666138 | -0.090274675 | 12 | 12 | 12 | 24.1 | 24.1 | 24.1 | 54.305 | 713460000 | 109 |
| VWF | P04275 | VWF | von Willebrand factor | 0.758773 | 0.0430247 | -0.801797723 | 53 | 53 | 53 | 19.8 | 19.8 | 19.8 | 309.26 | 5765000000 | 395 |
| WBP2 | K7EIJ0 | K7EIJ0 | WW domain-binding protein 2 | -0.113404935 | -0.14042035 | 0.253825478 | 4 | 4 | 4 | 25.7 | 25.7 | 25.7 | 18.224 | 214550000 | 22 |
| WDR1 | O75083 | WDR1 | WD repeat-containing protein 1 | 0.31862425 | -0.152377865 | -0.166246325 | 40 | 40 | 40 | 75.1 | 75.1 | 75.1 | 66.193 | 13469000000 | 493 |
| WDR44 | Q5JSH3 | WDR44 | WD repeat-containing protein 44 | 0.86920515 | -0.119632275 | -0.74957275 | 26 | 26 | 26 | 33.3 | 33.3 | 33.3 | 101.37 | 741200000 | 84 |
| YARS | P54577 | SYYC | Tyrosine--tRNA ligase, cytoplasmic | 0.4589253 | 0.340164535 | -0.7990898 | 16 | 16 | 16 | 29.5 | 29.5 | 29.5 | 59.143 | 203980000 | 43 |
| YWHAB | P31946 | 1433B | 14-3-3 protein beta/alpha | 0.755542125 | -0.43791925 | -0.3176229 | 17 | 7 | 7 | 58.5 | 35 | 35 | 28.082 | 1489400000 | 69 |
| YWHAE | P62258 | 1433E | 14-3-3 protein epsilon | 0.814420025 | -0.363997135 | -0.45042295 | 23 | 20 | 20 | 81.2 | 74.1 | 74.1 | 29.174 | 3551300000 | 202 |
| YWHAG | P61981 | 1433G | 14-3-3 protein gamma | 0.36637565 | -0.441083548 | 0.0747078 | 18 | 13 | 11 | 64.4 | 51.8 | 45.3 | 28.302 | 3575500000 | 139 |
| YWHAH | Q04917 | 1433F | 14-3-3 protein eta | 0.825055995 | -0.365944368 | -0.4591116 | 20 | 15 | 15 | 70.3 | 58.1 | 58.1 | 28.218 | 3813500000 | 145 |
| YWHAQ | P27348 | 1433T | 14-3-3 protein theta | 0.486682453 | 0.461285525 | -0.947967985 | 19 | 13 | 12 | 66.9 | 52.2 | 49.4 | 27.764 | 648870000 | 62 |
| YWHAZ | P63104 | 1433Z | 14-3-3 protein zeta/delta | 0.8121103 | -0.410033825 | -0.4020765 | 28 | 28 | 19 | 84.1 | 84.1 | 70.2 | 27.745 | 51807000000 | 536 |
| ZYX | Q15942 | ZYX | Zyxin | 0.444100741 | -0.392821925 | -0.051278675 | 26 | 26 | 26 | 56.3 | 56.3 | 56.3 | 61.277 | 12764000000 | 355 |
