## Supplemental Table 3 for "Proteome and functional decline as platelets age in the circulation"

Supplemental Table 3: Proteins significantly altered between young, intermediate and old platelets

| Gene Name | Protein ID | Protein Name | p value | Young Z score (average) | Intermediate Z score (average) | Old Z score (average) |
|---|---|---|---|---|---|---|
| IGLL5 | B9A064 | Immunoglobulin lambda-like polypeptide 5 | 0.0033 | -0.980255275 | -0.02746135 | 1.007716725 |
| SNP23 | O00161 | Synaptosomal-associated protein 23 | 0.0023 | 1.0689415 | -0.9344885 | -0.134452851 |
| ML12A | P19105 | Myosin regulatory light chain 12A | 0.0209 | 0.589969275 | 0.434207175 | -1.0241765 |
| SH3L1 | O75368 | SH3 domain-binding glutamic acid-rich-like protein | 0.0062 | 0.847148975 | 0.203274175 | -1.050422875 |
| B4DJV2 | B4DJV2 | Citrate synthase | 0.0200 | 0.95689195 | -0.144431975 | -0.81245995 |
| CSCL1 | O94886 | CSC1-like protein 1 | 0.0299 | 0.854480275 | -0.871299083 | 0.016818653 |
| LDHA | P00338 | L-lactate dehydrogenase A chain | 0.0016 | 1.160879225 | -0.774044712 | -0.38683435 |
| COX2 | P00403 | Cytochrome c oxidase subunit 2 | 0.0002 | 1.208486525 | -0.8837693 | -0.324717275 |
| CO5 | P01031 | Complement C5 | 0.0017 | -1.169010075 | 0.4507686 | 0.718241563 |
| TGFB1 | P01137 | Transforming growth factor beta-1 proprotein | 0.0186 | 0.6627928 | 0.359967825 | -1.022760475 |
| IGHG2 | P01859 | Immunoglobulin heavy constant gamma 2 | 0.0160 | 0.756935225 | -1.008197175 | 0.251262 |
| FIBA | P02671 | Fibrinogen alpha chain | 0.0126 | -0.639254065 | -0.420576308 | 1.0598306 |
| CXCL7 | P02775 | Platelet basic protein | 0.0405 | 0.918460225 | -0.198370164 | -0.720090208 |
| M0R009 | M0R009 | Alpha-1B-glycoprotein | 0.0197 | -0.942862 | 0.106139733 | 0.8367223 |
| RPN1 | P04843 | Dolichyl-diphosphooligosaccharide--protein glycosyltransferase subunit 1 | 0.0133 | 1.04196695 | -0.70627015 | -0.3356966 |
| RPN2 | P04844 | Dolichyl-diphosphooligosaccharide--protein glycosyltransferase subunit 2 | 0.0195 | 0.8291956 | 0.12042935 | -0.9496252 |
| ITB3 | P05106 | Integrin beta-3 | 0.0061 | 1.114821 | -0.529527275 | -0.58529376 |
| ADT2 | P05141 | ADP/ATP translocase 2 | 0.0425 | 0.805099275 | 0.05277355 | -0.857872875 |
| KPCB | P05771 | Protein kinase C beta type | 0.0164 | 1.0061022 | -0.24887048 | -0.757231833 |
| ATPB | P06576 | ATP synthase subunit beta, mitochondrial | 0.0001 | 1.167999975 | -0.176643198 | -0.991357025 |
| G6PI | P06744 | Glucose-6-phosphate isomerase | 0.0478 | 0.92121345 | -0.657599578 | -0.263613975 |
| CO8G | P07360 | Complement component C8 gamma chain | 0.0056 | -0.983261625 | 0.02736495 | 0.95589635 |
| CAN1 | P07384 | Calpain-1 catalytic subunit | 0.0149 | 1.048590525 | -0.42205305 | -0.626537405 |
| LYN | P07948 | Tyrosine-protein kinase Lyn | 0.0263 | 0.47373585 | -1.007796075 | 0.534060178 |
| ITA2B | P08514 | Integrin alpha-IIb | 0.0448 | 0.858075775 | -0.064105475 | -0.793970325 |
| A0A087X232 | A0A087X232 | Complement C1s subcomponent | 0.0079 | -0.659523588 | -0.4318083 | 1.09133185 |
| CO4A | P0C0L4 | Complement C4-A | 0.0043 | -1.124835225 | 0.689237 | 0.43559805 |
| HS71B | P0DMV9 | Heat shock 70 kDa protein 1B | 0.0220 | 0.56911135 | 0.45241826 | -1.02152962 |
| G6PD | P11413 | Glucose-6-phosphate 1-dehydrogenase | 0.0484 | 0.911229225 | -0.680610825 | -0.23061835 |
| FA5 | P12259 | Coagulation factor V | 0.0081 | 1.048784325 | -0.245233948 | -0.803550525 |
| SRC | P12931 | Proto-oncogene tyrosine-protein kinase Src | 0.0490 | 0.71178984 | 0.183996025 | -0.895785913 |
| COX41 | P13073 | Cytochrome c oxidase subunit 4 isoform 1, mitochondrial | 0.0488 | 0.039416 | 0.799426425 | -0.838842775 |
| NID1 | P14543 | Nidogen-1 | 0.0002 | 1.007460675 | 0.12864875 | -1.13610945 |
| Q5R345 | Q5R345 | P-selectin | 0.0152 | 0.735103275 | 0.286077138 | -1.021180343 |
| PGAM1 | P18669 | Phosphoglycerate mutase 1 | 0.0242 | 1.00771425 | -0.612611375 | -0.395102925 |
| HXK1 | P19367 | Hexokinase-1 | 0.0125 | 0.69684345 | 0.35244635 | -1.049289975 |
| Q5T985 | Q5T985 | Inter-alpha-trypsin inhibitor heavy chain H2 | 0.0460 | -0.881384525 | 0.126525978 | 0.754858775 |
| A0A1W2PQH3 | A0A1W2PQH3 | Malic enzyme | 0.0075 | 1.10212 | -0.568298148 | -0.533821963 |
| COF1 | P23528 | Cofilin-1 | 0.0259 | -0.942733775 | 0.158658175 | 0.784075675 |
| A6NLN1 | A6NLN1 | Polypyrimidine tract binding protein 1, isoform CRA_b | 0.0218 | -0.798815573 | -0.156575383 | 0.955390825 |
| PON1 | P27169 | Serum paraoxonase/arylesterase 1 | 0.0272 | -0.925790925 | 0.124133478 | 0.801657353 |
| PTN6 | P29350 | Tyrosine-protein phosphatase non-receptor type 6 | 0.0137 | 1.004291453 | -0.20395485 | -0.800336425 |
| E9PK01 | E9PK01 | Elongation factor 1-delta | 0.0375 | 0.9421844 | -0.68637005 | -0.255814253 |
| 2AAA | P30153 | Serine/threonine-protein phosphatase 2A 65 kDa regulatory subunit A alpha isoform | 0.0225 | 0.812887225 | 0.13003145 | -0.942918525 |
| TALDO | P37837 | Transaldolase | 0.0188 | 0.460416575 | 0.57456185 | -1.034978425 |
| GRP75 | P38646 | Stress-70 protein, mitochondrial | 0.0186 | 0.7189288 | 0.288608825 | -1.00753765 |
| A0A0C4DGS1 | A0A0C4DGS1 | Dolichyl-diphosphooligosaccharide--protein glycosyltransferase 48 kDa subunit | 0.0435 | 0.949605375 | -0.59117235 | -0.358432775 |
| GPDM | P43304 | Glycerol-3-phosphate dehydrogenase, mitochondrial | 0.0323 | 0.813045925 | 0.081157458 | -0.894203425 |
| ATPO | P48047 | ATP synthase subunit O, mitochondrial | 0.0457 | 0.858824325 | -0.7886979 | -0.070126325 |
| TCPG | P49368 | T-complex protein 1 subunit gamma | 0.0133 | 0.91580895 | -0.926627375 | 0.010818225 |
| EFTU | P49411 | Elongation factor Tu, mitochondrial | 0.0014 | 1.114380275 | -0.206287275 | -0.908093 |
| TMEDA | P49755 | Transmembrane emp24 domain-containing protein 10 | 0.0006 | 1.218011975 | -0.579017903 | -0.638994075 |
| GNAQ | P50148 | Guanine nucleotide-binding protein G | 0.0236 | 1.0174778 | -0.53500285 | -0.482475115 |
| EMD | P50402 | Emerin | 0.0499 | 0.943891725 | -0.497453425 | -0.44643845 |
| F10A1 | P50502 | Hsc70-interacting protein | 0.0032 | 1.089130875 | -0.86535865 | -0.223772225 |
| LRBA | P50851 | Lipopolysaccharide-responsive and beige-like anchor protein | 0.0011 | 1.121498575 | -0.92205065 | -0.19944795 |
| TCPD | P50991 | T-complex protein 1 subunit delta | 0.0138 | 0.785841825 | -1.010154545 | 0.22431295 |
| GDIR2 | P52566 | Rho GDP-dissociation inhibitor 2 | 0.0278 | 0.6935724 | 0.281072875 | -0.97464545 |
| ECHB | P55084 | Trifunctional enzyme subunit beta, mitochondrial | 0.0364 | 0.9565522 | -0.6523575 | -0.30419465 |
| IF4A1 | P60842 | Eukaryotic initiation factor 4A-I | 0.0397 | 0.89279075 | -0.12116854 | -0.771622275 |
| RHOA | P61586 | Transforming protein RhoA | 0.0498 | 0.941097325 | -0.4022544 | -0.538843075 |
| CH10 | P61604 | 10 kDa heat shock protein, mitochondrial | 0.0065 | 0.655959325 | 0.448780288 | -1.104739775 |
| UB2L3 | P68036 | Ubiquitin-conjugating enzyme E2 L3 | 0.0064 | 0.961357225 | 0.003713575 | -0.96507085 |
| EF1A3 | Q5VTE0 | Putative elongation factor 1-alpha-like 3 | 0.0017 | 1.093052175 | -0.162306525 | -0.930745763 |
| HBA | P69905 | Hemoglobin subunit alpha | 0.0056 | -0.284160825 | -0.7961623 | 1.080323025 |
| GSTO1 | P78417 | Glutathione S-transferase omega-1 | 0.0463 | 0.94983955 | -0.4130443 | -0.536795473 |
| MPCP | Q00325 | Phosphate carrier protein, mitochondrial | 0.0296 | 0.995723425 | -0.5473563 | -0.4483672 |
| CLH1 | Q00610 | Clathrin heavy chain 1 | 0.0144 | 0.955130525 | -0.08386499 | -0.87126538 |
| MMRN1 | Q13201 | Multimerin-1 | 0.0002 | 1.00382565 | 0.13461291 | -1.138438475 |
| NNTM | Q13423 | NAD(P) transhydrogenase, mitochondrial | 0.0201 | 0.926892 | -0.071259285 | -0.8556327 |
| STIM1 | Q13586 | Stromal interaction molecule 1 | 0.0021 | 1.1640877 | -0.492815408 | -0.671272408 |
| GNA13 | Q14344 | Guanine nucleotide-binding protein subunit alpha-13 | 0.0273 | 0.896501025 | -0.841097475 | -0.0554035 |
| ITIH4 | Q14624 | Inter-alpha-trypsin inhibitor heavy chain H4 | 0.0001 | -1.222240325 | 0.327501456 | 0.894738925 |
| GANAB | Q14697 | Neutral alpha-glucosidase AB | 0.0113 | 1.067012175 | -0.419102475 | -0.647909575 |
| MARE1 | Q15691 | Microtubule-associated protein RP/EB family member 1 | 0.0054 | 1.10636395 | -0.712499125 | -0.3938647 |
| VAMP3 | Q15836 | Vesicle-associated membrane protein 3 | 0.0172 | 0.891700625 | -0.915929725 | 0.024228875 |
| RTN1 | Q16799 | Reticulon-1 | 0.0026 | 1.1585001 | -0.624919253 | -0.533580925 |
| INF2 | Q27J81 | Inverted formin-2 | 0.0364 | 0.924796975 | -0.188287775 | -0.7365093 |
| LEGL | Q3ZCW2 | Galectin-related protein | 0.0248 | 0.602763175 | 0.404377875 | -1.007141125 |
| TWF2 | Q6IBS0 | Twinfilin-2 | 0.0070 | 1.105139175 | -0.4981927 | -0.60694655 |
| UN13D | Q70J99 | Protein unc-13 homolog D | 0.0295 | 0.9045756 | -0.087851771 | -0.81672395 |
| SND1 | Q7KZF4 | Staphylococcal nuclease domain-containing protein 1 | 0.0180 | 0.833391225 | 0.122558425 | -0.955949675 |
| GRP2 | Q7LDG7 | RAS guanyl-releasing protein 2 | 0.0281 | 0.985195275 | -0.33409651 | -0.651099093 |
| PKHL1 | Q86WI1 | Fibrocystin-L | 0.0168 | -0.999975725 | 0.23392575 | 0.766050075 |
| RHG18 | Q8N392 | Rho GTPase-activating protein 18 | 0.0494 | 0.8620322 | -0.0950281 | -0.7670042 |
| SCPDL | Q8NBX0 | Saccharopine dehydrogenase-like oxidoreductase | 0.0233 | 0.650036625 | -1.004592723 | 0.354556145 |
| C9JAI6 | C9JAI6 | CKLF-like MARVEL transmembrane domain-containing protein 5 | 0.0137 | 1.055469275 | -0.4295719 | -0.6258973 |
| J3KPX7 | J3KPX7 | Prohibitin-2 | 0.0002 | 1.129447675 | -1.036706825 | -0.092740615 |
| ESYT1 | Q9BSJ8 | Extended synaptotagmin-1 | 0.0448 | 0.853682775 | -0.05419165 | -0.799491225 |
| GTPB2 | Q9BX10 | GTP-binding protein 2 | 0.0419 | 0.4992062 | -0.96277056 | 0.463564325 |
| TMOD3 | Q9NYL9 | Tropomodulin-3 | 0.0492 | 0.9453638 | -0.444760875 | -0.50060255 |
| A0A087X054 | A0A087X054 | Hypoxia up-regulated protein 1 | 0.0401 | 0.890824575 | -0.772490775 | -0.118334075 |
| EMIL1 | Q9Y6C2 | EMILIN-1 | 0.0168 | 1.02802025 | -0.346318475 | -0.68170185 |
| Q5T0I0 | Q5T0I0 | Gelsolin | 0.0045 | 1.1057838 | -0.76324425 | -0.342539425 |
